# RNA virus ecogenomics along a subarctic permafrost thaw gradient

**DOI:** 10.1101/2025.02.13.637936

**Authors:** Akbar Adjie Pratama, Guillermo Domínguez-Huerta, James M. Wainaina, Benjamin Bolduc, Funing Tian, Jared Ellenbogen, Jiarong Guo, EMERGE Field Team, EMERGE Coordinators, Jens H. Kuhn, Kelly C. Wrighton, Kisten Küsel, Ahmed A. Zayed, Matthew B. Sullivan

## Abstract

Climate change thaws permafrost, which releases greenhouse gases partly from dormant microorganisms awakening and metabolizing organic matter. Though DNA viruses that infect these soil microbes have been studied, little is known on soil RNA viruses, which typically infect microeukaryotes. Here we identify and characterize 2,651 RNA viruses from a 4-year time series of bulk soil metatranscriptomes derived from the climatically fragile Stordalen Mire ecosystem —a long studied permafrost peatland. RNA virus diversity was structured by habitat (palsa, bog, and fen), and these patterns correlated with pH and carbon dioxide and methane emissions. Further, host prediction, virus-encoded metabolite-transforming and information-processing functions suggested roles in ecosystem-scale carbon fluxes and contribute to greenhouse gases emissions. Together, these RNA virus ecogenomic data in permafrost provide essential baseline information for integration into predictive models to support hypothesis testing.

---

Climate change accelerates permafrost thaw through reduced ice cover lessening albedo (*1*). To study this, a model ecosystem – Stordalen Mire in northern Sweden – was established decades ago and revealed that permafrost frozen for thousands of years (*2*) now thaws near-annually (*3*). Concomitant with such thaw, dormant Stordalen Mire microorganisms are also annually awakened and in turn metabolize organic matter that accelerate greenhouse gas (GHG) emissions (*4*, *5*). Globally, such processes could unlock permafrost soil carbon reserves three times greater than the carbon currently in the atmosphere (*6*, *7*), though how specific soils and their microbiomes respond are among the greatest consequential unknowns in climate models (*8*). Thus, mechanistic understanding of the interplay between thawing permafrost, microbial communities (bacteria and archaea) and their viruses, and carbon dynamics represent a key grand challenge for climate science.

Previous studies in Stordalen Mire have characterized three distinct habitats representing a thaw gradient: palsas (partially frozen), bogs (intermediate thaw), and fully-thawed fens – using metagenomic sequencing, functional genomics, and ecosystem measurements (*5*, *8–10*). These studies identified dominant microbial phyla (e.g., Acidobacteria and Actinobacteria) and demonstrated that microbial communities varied across the permafrost thaw gradient. Additionally, key microbial metabolic pathways, including polysaccharide degradation and methane metabolism, revealed their roles in altering ecosystem greenhouse gas (GHG) outputs across the thawing gradients (*5*). Among the notable findings, the syntrophy between *Candidatus* Methanoflorens and *Candidatus* Acidiflorens has been shown to be ubiquitous for regulating carbon dynamics in such soils (*5*, *9*, *11*). Further, double-stranded (ds)DNA viruses – important in marine microbial biogeochemistry (via lysis, gene transfer, and metabolic reprogramming) (*12–14*) and frequently overlooked in microbiome studies and climate change modeling (*15*, *16*) – also appear important to GHG and ecosystem processes at Stordalen Mire. Specifically, ecogenomic analyses predict that dsDNA viruses are diverse and modulate microbial ecosystem outputs via infecting key microbial hosts and encoding auxiliary metabolic genes (AMGs) (*17–19*).

Despite these advances, the roles of other biological entities, such as mobile genetic elements and RNA viruses, remain largely unexplored in permafrost ecosystems. Recent reviews suggest soil RNA viruses influence food web structure, energy flow, and nutrient cycling, potentially shaping soil health and fertility (*20*). Given the unique environmental conditions of permafrost ecosystems, RNA viruses may play an even more pronounced role in shaping microbial communities and regulating carbon cycling and GHG. Here, we address this gap by investigating RNA viruses (the focus here) and mobile genetic elements (see Companion Guo et al., mobilome) in Stordalen Mire. By leveraging four years of bulk soil metatranscriptomes, we provide the first comprehensive analysis of RNA viruses across the permafrost thaw gradient, revealing their diversity, distribution, and functional potential in regulating ecosystem processes. Specifically, we screened the data to identify 25,337 viral contigs that were dereplicated into 2,651 virus operational taxonomic units (vOTUs), which we classified and ecologically contextualized. Similar to their dsDNA counterparts, Mire RNA viruses were largely novel and likely play diverse and significant roles in ecosystem processes. Our analyses revealed that RNA virus distribution is influenced by environmental factors such as pH and CH_4_ concentrations. Moreover, detailed examination of host-virus interactions and functional potential suggested these viruses play a crucial role in regulating carbon cycling and greenhouse gas emissions at the ecosystem scale. Studying RNA viruses also offers a unique lens into the underexplored eukaryotic communities in these environments, further underscoring their importance in permafrost regions.

## Stordalen Mire hosts thousands of novel RNA viruses

To identify RNA viruses in Stordalen Mire, here we focused on RNA viruses of the kingdom *Orthornavirae*, which replicate their genomes via RNA-directed RNA polymerase (RdRp). Hence, we screened the RdRp gene across ≈630 Gbp of bulk soil RNA metatranscriptomes sequencing data (*n*=55) collected from Stordalen Mire. These samples were obtained from three canonical permafrost thaw habitats (palsa, bog, and fen) and spanned across the soil active layers over the four years (2010, 2011, 2012 (*5*, *17*) and 2016 (*10*, *19*)). To enhance the detection of divergent RNA viruses, we iteratively searched and refined RdRp hidden Markov models (HMMs), a strategy previously successful (*21*, *22*) (**fig. S1**). Together these efforts identified 25,337 RdRp-containing contigs (RdRps length mean ± s.d.: 185.47 ± 112.30) and removed 3,225 false positives (see Methods). Of the resultant 22,112 RdRps that could be used for ecological analyses, 1,702 were complete or near-complete such that they could be used in deeper taxonomic analyses (**fig. S2**).

To determine the megataxa (phyla and classes) of RNA viruses present in this ecosystem, we utilized a previously benchmarked approach based on Markov clustering algorithm (MCL) analysis of RdRps (*22*). First, because global phylogenies of the >3-billion-year-old RdRp-encoding gene are challenging (*23*), we applied machine learning to pre-sort RdRp sequences into five orthornaviraen phyla that are recognized by the International Committee on Taxonomy of Viruses, or ICTV (*Duplornaviricota, Kitrinoviricota, Lenarviricota*, *Negarnaviricota,* and *Pisuviricota*) (*23*, *24*). Note that our study did not consider the recently established sixth RdRp-encoding phylum, *Ambiviricota* (*23*, *25*), as analyses were closed before ambiviricots were described. For the machine learning pre-sorting step, we specifically applied a network-based, iterative clustering approach to the 1,702 complete/near-complete RdRp sequences and another 704,003 complete RdRp references sequences derived as follows: 6,732 from the global *Tara* Oceans project (*22*), 3,325 from coastal seawater near the Yangtze River estuary (*26*), and 695,648 from GenBank (version 245). This merged network analysis of near-complete RdRp sequences revealed a 96% adjusted rand index (98.7% agreement, the number of correct assignments per total number of assignments to ICTV taxonomy) and 18 distinct “megaclusters” (**fig. S3; Table S5**). Of the ten megaclusters that contained Stordalen Mire sequences, seven matched ICTV-taxa within the five established phyla of *Duplornaviricota, Kitrinoviricota, Lenarviricota*, *Negarnaviricota,* and *Pisuviricota* (**Fig. 1A**). In addition to the established phyla, lineages of *incertae sedis* viruses, i.e., Permutotertaviridae (‘permutotetra-like’) and ‘kitrino-like’ viruses, were included as well. These 5 phyla include a total of 20 classes and Stordalen Mire sequences are assigned to 16 of them (**Table S5**), and at vOTUs/“species”-level, RNA viruses are all different (**fig. S2F**).

**Figure 1.**
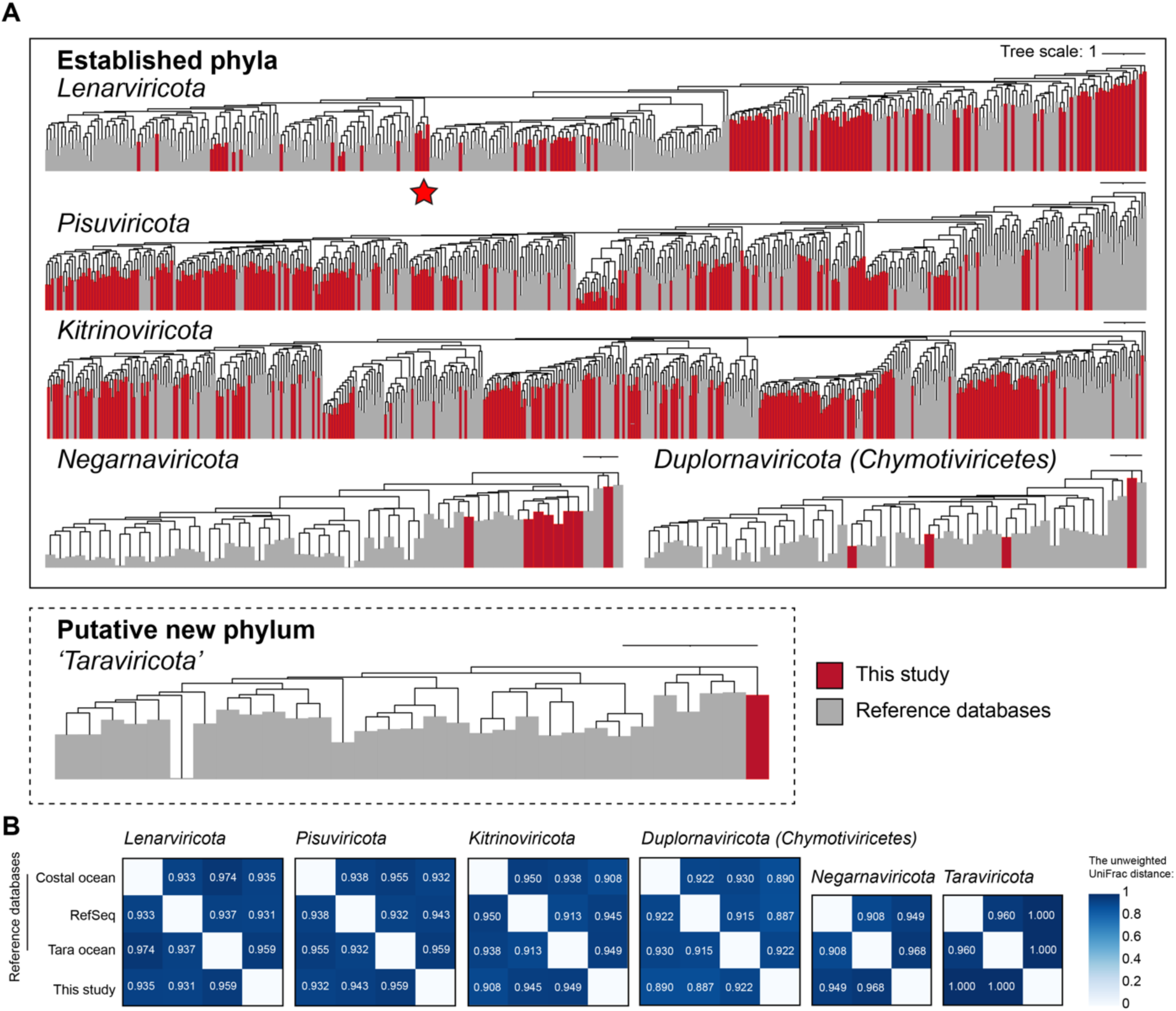
Phylum-ranked, RdRp-based orthornaviran phylogenies. **(A)** Six maximum-likelihood phylogenetic trees were built from an RdRp-guided taxonomy analysis of near-complete RdRp sequences. Five trees depict ICTV-established phyla, and one corresponds to a previously suggested phylum, ‘*Taraviricota*’ (Zayed et al.,(*22*)). The red color indicates Stordalen Mire orthornaviran sequences discovered in this study, whereas grey indicates previously known reference sequences (including open-ocean and coastal-ocean RNA viromes). The scale bar indicatesone amino acid residue substitution per site. The ⋆ represents a likely new class, Candidatus ‘Stormiviricetes’. RdRp, RNA-directed RNA polymerase; ICTV, International Committee on Taxonomy of Viruses. **(B)** Heatmap showing the unweighted UniFrac distances between RNA virus phylum-rank trees across datasets from the coastal ocean (*24*), RefSeq, *Tara* Oceans (*21*, *22*), and Stordalen Mire (this study). Phyla analyzed include *Lenarviricota, Pisuviricota, Kitrinoviricota, Duplornaviricota* (Chymotiviricetes), *Negarnaviricota*, and ‘*Taraviricota’*. Higher values (darker cells) indicate greater phylogenetic dissimilarity between datasets, whereas lower values (lighter cells) indicate greater similarity.

Beyond ICTV-recognized taxa, from our analysis, we also identified the recently reported megataxa ‘*Taraviricota*’ (*25*) of Stordalen Mire RNA viruses. This new phyla is not currently ICTV-recognized due to a lack of complete genomes, but other studies support the taxonomic assertions (*27*, *28*) (**fig. S3**; see **Table S5**). Moreover, we identified four novel virus operational units (vOTUs) encoding unclassified RdRps that clustered within *Lenarviricota* as a novel class tentatively named novel class “*Stormiviricetes*” (**fig. 1A, Fig. S3**). Their genomes ranged from 2.5 to 5.8 kb in length, and the longest one encoded two partially-overlapping open reading frames (**fig S4**). Given they are lenarviricots, we presume these are positive-sense, single-stranded RNA viruses.

To assess the similarity of Stordalen Mire RNA viruses with known reference sequences, we performed phylogenetic distance analysis using UniFrac. Phylogenetic analyses revealed permafrost virus-enriched clusters within the phyla *Lenarviricota, Kitrinoviricota, Negarnaviricota, Duplornaviricota (chrymotiviricetes),* and *Pisuviricota*, often forming phylogenetically distinct blocks (UniFrac distance >0.9) with limited reference sequences outside Stordalen (**Fig. 1B; fig. S5-S12**). Aligned to the previous literature (*29*, *30*), which found that permafrost hosts harbor phylogenetically distinct RNA viruses—including reoviruses (Duplornaviricota), hypoviruses (Pisuviricota), rhabdoviruses (Negarnaviricota), and leviviricetes—our findings reinforce the uniqueness of RNA viruses in thawing permafrost (*29*). Comparing these patterns to other soil ecosystems, we find distinct differences in RNA virus composition. For example, in agricultural soils, leviviricetes (*Lenaviricota*) are most represented (*31*), whereas soils with added root litter show increased levels of barnavirids (*Pisuviricota*), megabirnavirids, and quadrivirids (*Duplornaviricota*), along with mitovirids (*Lenarviricota*) and leviviricetes (*32*).

## Ecological patterns of Stordalen Mire RNA viruses

We next sought to assess the ecological distributions of RNA viruses. We used a recent community consensus definition (*33*) for OTUs of RNA viruses, preclustered at 90% average nucleotide identity across 80% of the shorter sequence length and 1 kb minimum contig length (materials and methods) (*21*, *22*). Clustering using these parameters collapsed the 22,112 Stordalen Mire RdRp sequences to 2,651 vOTUs (**fig. S13-S15**). Compared to other studies, this number is relatively large and suggests high soil RNA viral diversity considering it derives from a single sampling location and yet is nearly half that (2,651 vs 5,504 vOTUs) found throughout the global oceans (*22*). Consistent with such high diversity, individual samples contributed from ∼83 to 166 vOTUs on average, and accumulation curves revealed sampling was not yet saturated (**fig. S13C)**; though most large-scale RNA virus surveys of marine, soil, and engineered environments fail to saturate RNA viral diversity (*27*, *34*, *35*).

To assess this further within soils - a biome where microbiomes are considered particularly diverse (*36*, *37*) - we collated and reanalyzed all available soil metatranscriptomes using our iterative HMM RdRp search strategy. This revealed the following number of RNA virus vOTUs: 83,027 vOTUs across 288 diverse Chinese soil samples (*38*), 3,161 vOTUs across 83 Warwick-England crop agricultural soil samples (*31*), 2,112 vOTUs across 48 California rhizosphere soil samples (*39*), and 1,412 vOTUs across 16 Welsh upland peatland samples (*32*). The comparison, based on the vOTU clustering at 90% nucleotide identity and 80% coverage, across these different soil RNA studies revealed that only 62 vOTUs overlapped to Chinese soils and 28 vOTUs to those Welsh upland peatland (**fig. S16C**). Further, 98% of the Stordalen Mire vOTUs were unique across these soil studies, similar to the high proportion of study-specific unique viruses detected in other soil datasets (**fig. S6C**).

We next explored vOTU-based community structure and patterns as follows. First, we explored beta-diversity of RNA vOTU relative abundances via ordination and hierarchical clustering of Bray-Curtis dissimilarity, which revealed three distinct meta-communities that matched the major soil types (palsa, bog, and fen) (PERMANOVA, *p* = 0.001; **Fig. 2B, fig. S13-S15**). This approach added RNA viruses to habitat-specific clustering of plants, microbiomes, and dsDNA viruses at this site (*17*, *19*, *40*), similar to RNA virus community differentiation across soil types observed in other studies (*31*, *32*, *38*, *39*, *41*). This structuring in Stordalen Mire may be shaped by the distinct plant communities associated with different thawing stages, as Sphagnum mosses, hare’s-tail cottongrass (*Eriophorum vaginatum*), and common cottongrass (*Eriophorum angustifolium*) dominate the palsas, bogs, and fens, respectively (*42–47*).

**Figure 2.**
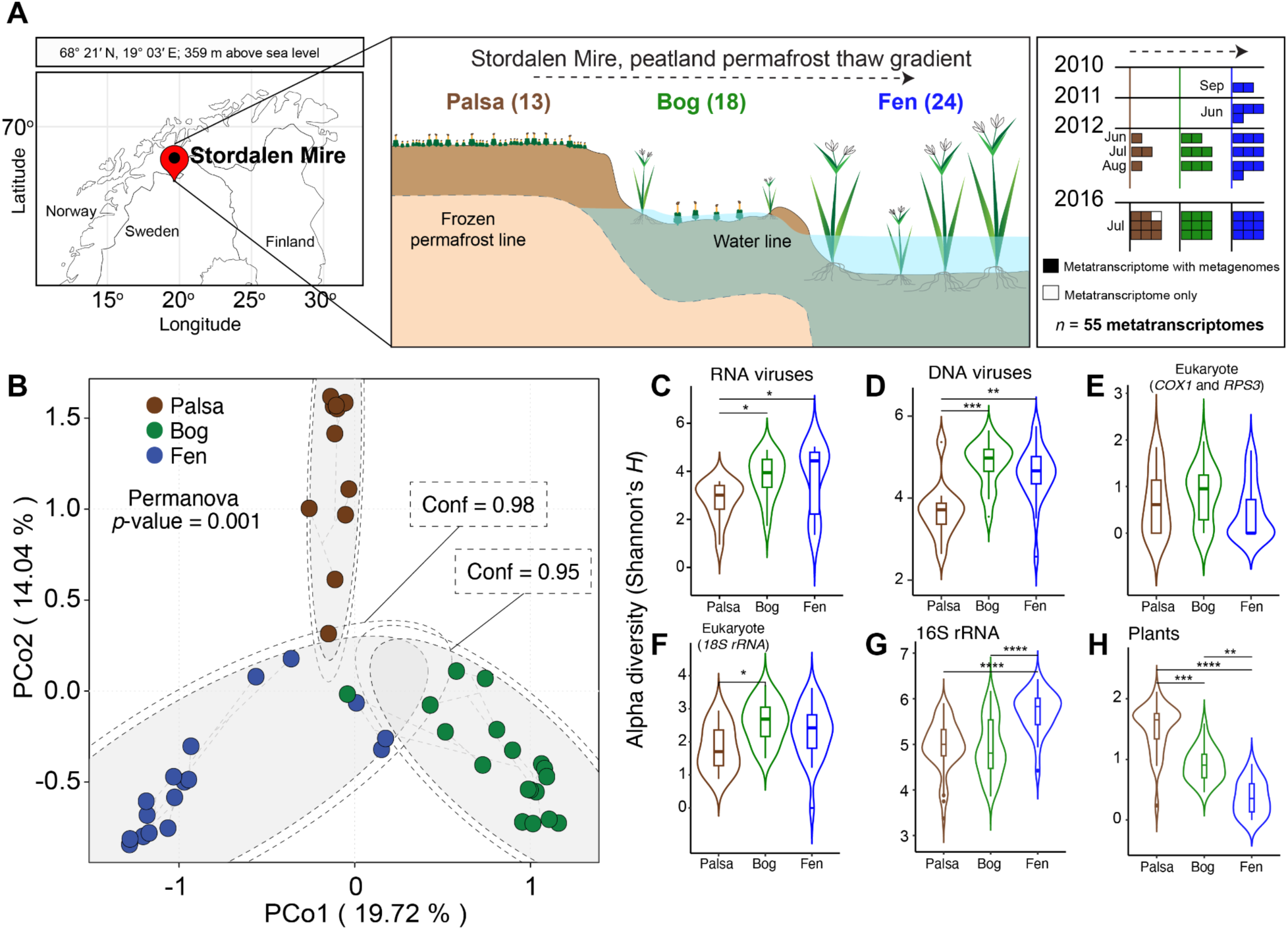
Cross-habitat soil sampling of Stordalen Mire in northern Sweden. **(A)** Soil metatranscriptomes were collected from the active layer across the peatland permafrost thawing gradient with intact palsa (approximately 30 cm), bog (intermediate thawing, approximately 60 cm), and fen (fully thawed) habitats. **(B)** Principal component analysis (PCoA) of a Bray–Curtis dissimilarity matrix calculated from all virus operational units (vOTUs; i.e., species-rank units) uncovered in this study. Dot colors correspond to the colours of the thawing gradient (brown for palsa; green for bog; and blue for fen. Alpha-diversity (Shannon’s *H*) of **(C)** RNA viruses, **(D)** dsDNA viruses, **(E)** Eukaryote and prokaryote (based on *COX1* and *RPS3*), **(F)** Prokaryote (based on 18S rRNA), **(G)** 16S rRNA, and **(H)** Plants. Violin plots with boxplots show medians and quartiles. Statistical analysis was performed using the Kruskal–Wallis test with *post hoc* Dunn test and *p*-adjusted Bonferroni. Only the significant values are shown and donated: * *p*-value ≤0.05; ** *p*-value ≤0.01, *** *p-*value ≤0.001, and **** *p*-value ≤0.0001. Fig. 2 G and H on the complementary papers.

Next, we assessed alpha-diversity (Shannon’s *H*) of RNA viruses and compared it to broader Stordalen Mire taxa including dsDNA viruses and microbial communities (eukaryotes and prokaryotes). Mire RNA viral diversity increased along the thaw gradient, with the highest diversity observed in fens followed by bogs and the lowest at palsas, just as seen for gene-marker-inferred alpha-diversity for eukaryotes assayed here (*18S rRNA*; **Fig. 2F**) and prokaryotes assayed elsewhere (16S rRNA; complementary paper). Interestingly, dsDNA virus diversity peaks in bogs but does not differ significantly from that in fens (**Fig. 2D**). However, alpha diversity measured for plant diversity exhibited distinct trends, no significant difference across palsa, bog and fen and peaks in palsas for plants, respectively (**Fig. 2F, 2H**). These differences likely reflect the varying resolution of the markers used, *COX1* and *RPS3* provide higher taxonomic resolution (*39*, *48*), whereas 18S rRNA is more conserved and better suited for broad taxonomic overviews but lacks the resolution to capture finer-scale variation (*49*). Though prior soil RNA virus studies have documented RNA virus diversity (*31*, *32*, *38*, *39*, *41*), few assessed broader communities taxa within their sites (*31*, *39*). Interestingly, while dsDNA virus diversity peaked in bogs, its Shannon’s *H* in fens was not statistically significant. Given that fens represent fully thawed, inundated permafrost systems, these results suggest that permafrost thaw is associated with increased RNA viruses, eukaryotes, and microbial diversity. However, the broader ecological consequences of these changes remain uncertain.

## Ecological drivers of orthornaviraens in thawing permafrost

We next applied ecological driver analyses to understand how biotic and abiotic factors impact Stordalen Mire RNA viral community structure across habitats and environmental factors. Myriad analyses revealed that Stordalen Mire RNA virus community structure was most impacted by pH, followed distantly by δ^13^C (CH_4_), δ^13^C (CO_2_), and total relative abundance of organisms (eukaryote) (goodness of fit and Mantel tests in **Fig 3B-C**, relative importance analysis in **fig. S17,** and redundancy analysis in **fig. S18**). Previously, prokaryotic dsDNA viruses have shown pH-influenced community structure at local and global scales (*50*). In thawing permafrost systems, pH can impact ecological parameters such as microbial activity, nutrient availability, and plant species composition by modulating the decomposition of newly thawed organic matter. Thawing also leads to organic acid accumulation in low-pH bogs, while the higher pH in fens promotes bicarbonate formation, making them more alkaline (*51*, *52*), which in turn could have an indirect effect on RNA virus communities via host composition selection (as seen prior for prokaryotes in this ecosystem, Emerson et al (*17*)). Alternatively, pH might directly alter virus community structure through conformational changes critical to virus integrity and infectivity, as known for viral pathogens (*53*). As pH emerges as a large-scale prominent driver of RNA virus community structure, supporting prior findings for prokaryotic dsDNA viruses (*50*) linking RNA virus diversity to pH, nutrients, and organic content in soil (*38*).

**Figure 3.**
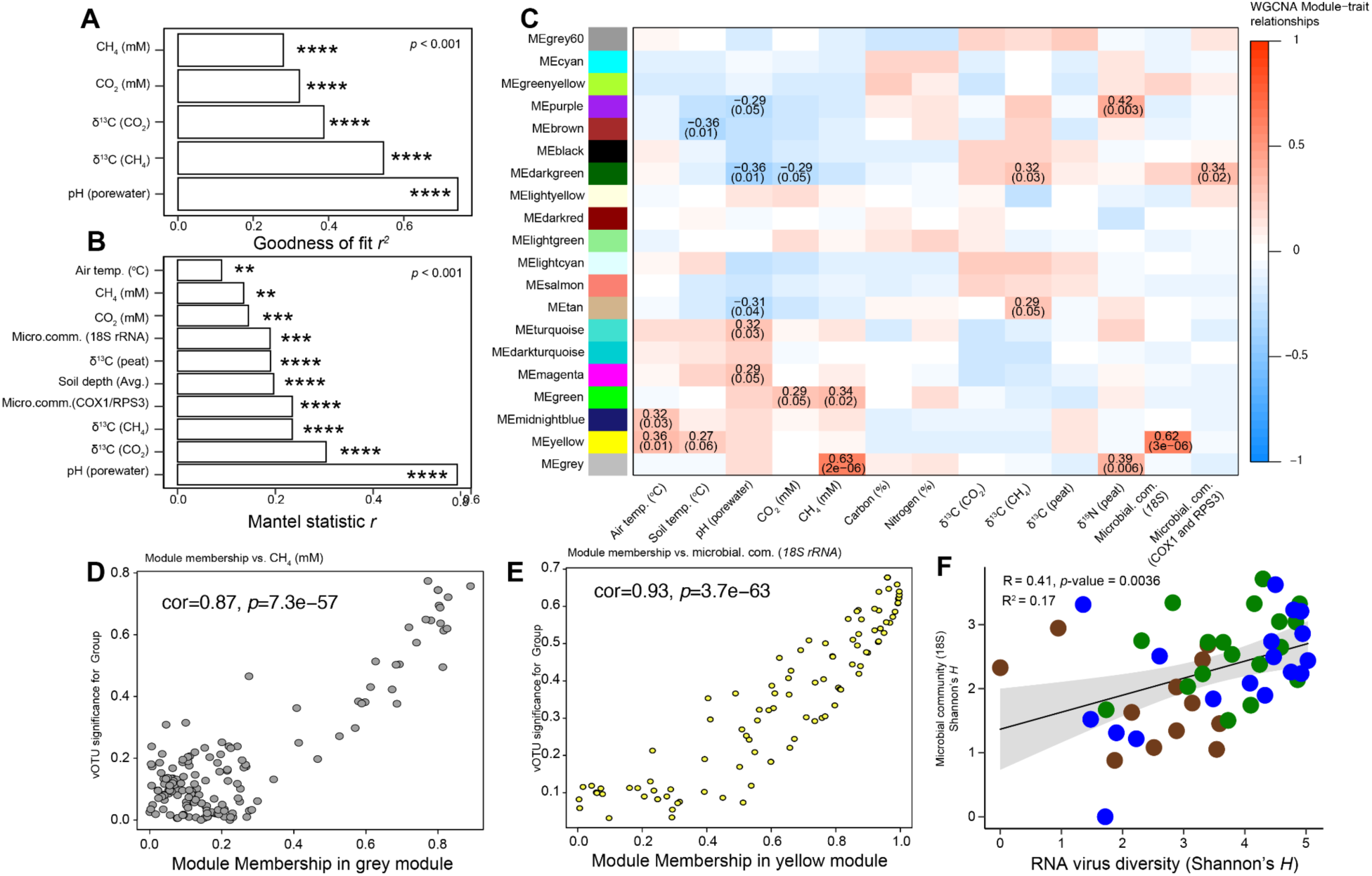
Ecological drivers of Stordalen Mire orthornaviraens. (**A, B)** The first two dimensions (with a goodness of fit *r*^2^ derived from a generalized additive model) and all dimensions (determined through a Mantel test based on Spearman’s correlation) revealed potential ecological drivers and predictors of orthornaviran beta-diversity. pH was consistently identified as the strongest predictor of beta-diversity. Only the significant values are shown and donated as follows: * *p*-value ≤0.05; ** *p* -value ≤0.01, and *** *p* -value ≤0.001. (**C**) Heat map of module-trait relationships derived from Weighted Gene Co-Expression Network Analysis (WGCNA). Each row corresponds to viral operational taxonomic units (vOTUs), grouped into modules (labeled by module colors), and each column represents an environmental trait. The color scale indicates the correlation coefficients. The correlation values and corresponding *p*-values (in parentheses) are shown within each cell; only *p*-values ≤0.05 are shown. (**D, E**) Correlation analysis between module membership in the grey and yellow modules vs. methane concentration (CH₄ mM) and microbial community (*18S rRNA*) relative abundance, respectively. Each point represents a vOTU within the grey and yellow modules. (**F**) Spearman’s correlation analysis between orthornaviran and host cellular organism diversity (Shannon’s *H*).

Having better understood drivers of community structure, we next sought to assess which vOTUs best predicted ecosystem features of interest across Stordalen Mire. To this end, we used weighted gene/genome correlation network analysis (WGCNA) to group vOTUs into modules and assess modular correlations to environmental factors as developed for ocean biota (*54*) including application for identifying key ocean DNA and RNA viruses that best predict carbon flux (*21*, *55*). Of 13 environmental traits assessed, four factors, including carbon (%), nitrogen (%), isotopic signature of carbon dioxide, and carbon in peat, are not the best predictors of RNA virus diversity. Instead, air and soil temperature, pH, CH_4_, CO_2_, and the relative abundance of microbes across the thawing permafrost were prominent predictors (**Fig. 3C**). Moreover, our analysis also showed 12 of 20 vOTU modules correlated to these environmental factors (*p* ≤ 0.05). Among these, the WGCNA gray module strongly correlated with CH₄ concentrations (corr. = 0.87, *p* = 7.3e-57; **Fig. 3D**). While dsDNA viruses have been speculated to indirectly impact methane cycling via methane-related AMGs (*17*, *19*, *56*) or microbial lysis dynamics (*57*, *58*), our result provides a connection between RNA viruses and CH₄ concentrations in thawing permafrost systems. However, the mechanism of RNA viruses influencing CH₄ in thawing permafrost remains unclear, possibly through host interactions. Toward the latter consideration, our analysis also found the WGCNA yellow module showed the strongest correlation with organismal abundance (corr. = 0.94, *p* = 4e-69; **Fig. 3E**). This extends long-standing theories such as “kill the winner,” (*59*) which suggests that host-driven shifts in microbial ecosystems could also indirectly influence RNA viruses and their ecological roles, including potential links to methane-related processes. A subsequent correlation analysis of RNA virus richness and diversity (Shannon’s *H*) with host abundance supported this hypothesis, showing a significant positive relationship (**Fig. 3F**). Together, these findings raise new questions about the ecological roles of RNA viruses in rapidly changing environments and underscore the need for future mechanistic studies to explore their contributions to methane-related biogeochemical processes.

## RNA viruses infect diverse eukaryotic hosts that are key components of food webs and biogeochemistry in Stordalen Mire

Past studies have begun to uncover RNA viruses that affect multiple trophic levels (bacteria, protists, fungi, and invertebrates), as well as inferred carbon cycling impact through host lysis (*32*, *39*). However, soil RNA virus ecological roles are largely underexplored.

To provide context for the organisms inhabiting the ecosystem, we screened for hallmark genes for eukaryotes (*COX1*, *RPS3*, 18S rRNA) and prokaryotes (*RPS3*, 16S rRNA) (**Fig. 4**, **fig. S19**) and estimated their abundances across the metagenomes (**fig. S19**). Bacteria and archaea are well known at the site due to past genome-resolved studies (*5*), but known RNA viruses more commonly infect eukaryotes (*21*, *60*), so we expected the same here. Since the eukaryotic communities at Stordalen Mire are not yet characterized, we describe their patterns here. Despite discrepancies found in the organisms inferred from the different gene screenings —likely due to bioinformatic or biological biases) (**fig. S19** and **fig. S20**—, they suggested a diverse taxonomic repertoire of protists, fungi, plants, and invertebrates with habitat-dependent abundance profiles as follows. Palsas were enriched in plants (monocotyledons of Poales, likely hare’s-tail cottongrass, *E. vaginatum* (*42*)), providing primary production that supports heterotrophs with diverse ecological roles: saprophytic and mycorrhizal (Agaricales and Helotiales) and pathogenic fungi (Mycosphaerellales), litter-decay insects (Entomobryomorph and Poduromorph springtails), rodents, and parasitic apicomplex protists (Eugregarinorida). Wetter bogs were dominated by fungi (Heliotales), tardigrades (Parachela), bacteriovorous rotifers (Adinetida, Ploima) and ciliates (Haptorida). Inundated fens were dominated by oligochaetean worms (Haplotaxida and Lumbriculidae), likely feeding on the rich bacteria and archaea biomass fueled by peatland carbon, and monocotyledons of Poales (likely, *Carex* sp. and *Eriophorum* sp.). In general, plants, fungi and herbivorous invertebrates were well distributed along the thawing gradient. In bogs and fens, prokaryotic life is grazed by bacterivorous protists and invertebrates (*61*), with protists possibly being outcompeted by invertebrates (**fig. S20**) as previously reported in inundated environments (*62*). With this mapped backdrop of potential host taxa across the Mire, we next assessed whether our newly identified RNA viruses might infect any of these organism taxa as predicted by the following *in silico* approaches. Unlike DNA viruses, RNA virus host prediction is not as well developed because orthornaviraens lack programmed endogenization into the host genome. Thus, as done previously (Zayed et al (*22*)), we combined host predictions from multiple approaches based on different types of features derived from RNA virus contigs as follows: (*i*) prior host information of known virus taxa classified using the RdRp sequences (**fig.S21,23**), (*ii*) similarity to closely-related endogenous virus elements that reside in known cellular genomes (**fig. S21**), (*iii*) prediction of alternative genetic codes that suggest possible host organisms and subcellular compartments (**fig. S21**), and (*iv*) prediction of Shine-Dalgarno motifs and CRISPR spacers that suggest bacterial hosts (**fig. S21, 22**). Where more than one line of evidence existed, the final host prediction was assigned using a hierarchical scoring approach that differentially weighted the predictions according to confidence (see materials and methods).

**Figure 4.**
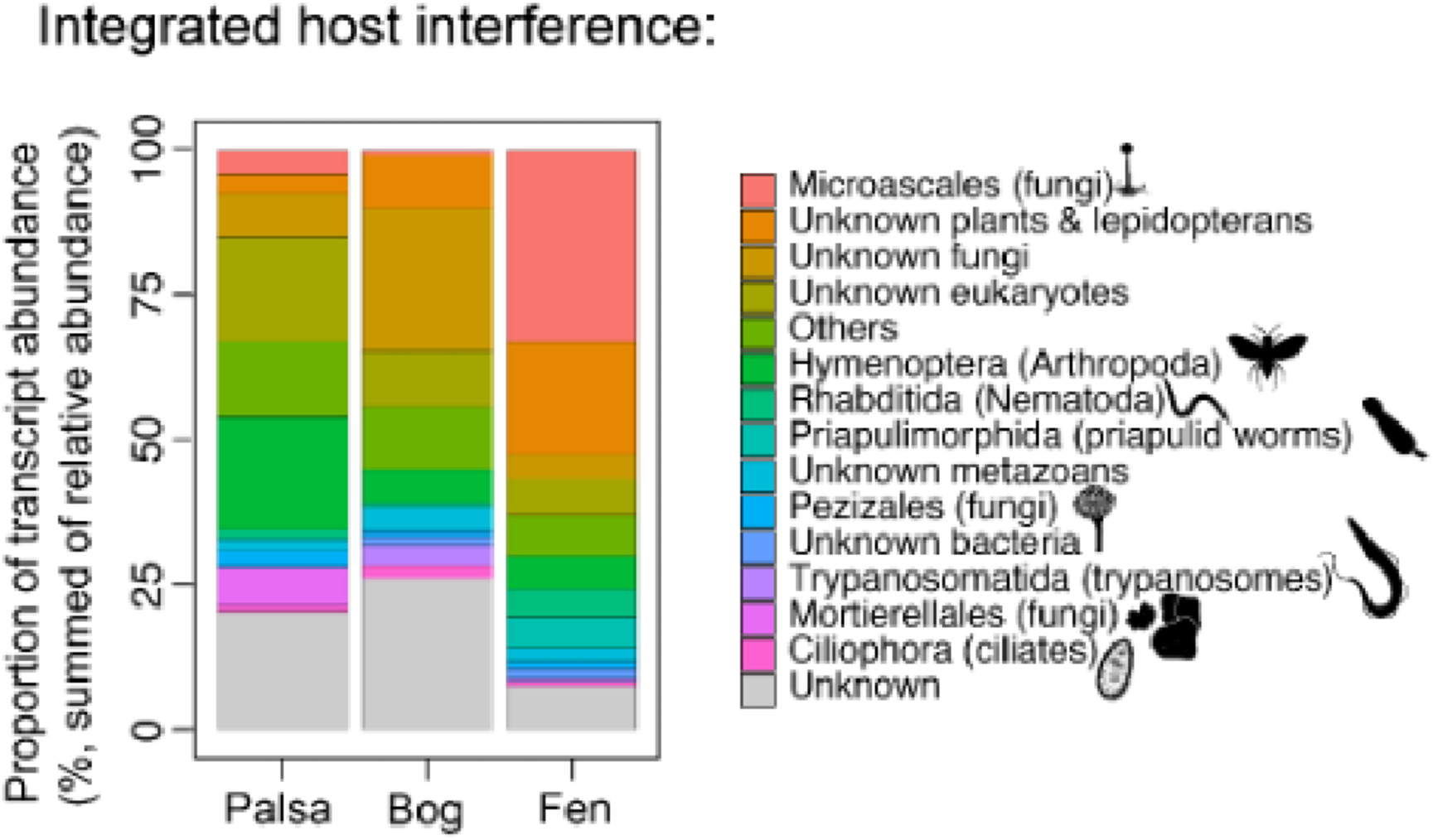
Integrated host inferences, and the co-phylogeny relationship between Stordalen Mire orthornaviraens and their predicted cellular hosts. **(A)** Taxonomic assignment of the prokaryotes and eukaryotes in Stordalen Mire identified via *COX1* and *RPS3* gene and 18S rRNA analysis. Cytochrome c oxidase subunit 1 mitochondrial gene (*COX1*); a component of the eukaryotic 40S ribosomal subunit, ribosomal protein S3 (*RPS3*); and small subunit ribosomal RNA (*18S rRNA*).

Overall, the approach worked well to provide at least some taxon rank host prediction for the vast majority (1,871 or 70.9%) of the 2,561 vOTUs (**Fig. 4C**, individual host inference approach results are in **fig. S21, S22** and **Table S6**). These efforts revealed that Stordalen Mire RNA viruses most commonly infected fungi (e.g., Microascales, Pezizales), invertebrates (e.g., Hymenoptera, Rhabditida), and protists (e.g., Ciliophora, Trypanosomatida). Similarly to what we report here, RNA viruses that putatively infect fungi are consistently found as abundant in diverse soils (*32*, *38*, *39*, *63*) and those that infect other eukaryotic hosts (protists and invertebrates) are more rare (*32*, *38*), although this fact may derived from methodological differences. Notably, by integrating RdRp assignment to *Leviviricetes* and prediction of Shine–Dalgarno motifs and CRISPR spacers, only 69 vOTUs (2.7%) putatively infect bacteria, and these were not particularly abundant as they proportionally comprised less than 2% of the vOTU abundances (**fig. S19, S20**). By contrast, prior soil surveys have observed more than an order of magnitude higher relative abundances of bacterial RNA viruses, with more than a third of the leviviricete vOTUs (*32*). However, the predominance of RNA viruses likely infecting eukaryotes is consistent with findings in the global oceans (*21*, *22*, *60*).

Beyond these qualitative inferences, we next sought to more quantitatively assess virus-host interaction biology across the Mire by assessing vOTU abundances in light of their predicted hosts (**fig. S3**). In palsas, RNA viruses commonly infected hymenopteran insects (19.4%), unknown fungi (7.5%), mortierellalian fungi (6.6%), and microascalian fungi (4.0%) (**fig. S19**, **S20**), all of which ultimately derived carbon from plants that support herbivorous, parasitic, symbiotic, and saprotrophic life around them in this habitat (*5*, *31*). In bogs, RNA viruses infected unknown fungi (24.7%), plants, lepidopterans (9.1%), hymenopterans (5.8%), and protists (ciliates and trypanosomatids, 5.6%) (**fig. S19, fig. S20**). By contrast, in the fully inundated fens, RNA viruses mostly infected microascalian fungi (33.1%), unknown plants and lepidopterans (19.5%), hymenopterans (5.7%), priapulids (5.3%), and rhabditids (4.9%) (**fig. S19, S20**). In wetter habitats like bogs and fens, peatlands provide a rich source of biogenic carbon, supporting abundant bacteria and archaea (which increase CO_2_ and CH_4_ emissions), which in turns fuels active grazing. This shift in the carbon sources could explain the higher frequency of protists in bogs and non-arthropod invertebrates in fens compared to bogs.

Extrapolating from what is known about these predicted hosts, we propose the following ways RNA virus infection might modulate key ecological roles that impact soil carbon cycling. First, impacts on primary production of the Mire ecosystem through infection of plants and photosynthetic protists. Second, infection of fungi could impact their multiple roles in rhizosphere building, saprotrophic degradation of litter and peatland and parasitism. Third, among other functions, bacterial and archaeal predation could be influenced by the infection of protists and non-arthropod invertebrates (e.g., priapulids, nematodes). Critically, because RNA viruses that infect eukaryotes are known to utilize a range of infection strategies from cryptic or persistent to acute infections, associated with different levels of impact on the hosts, this viral infection continuum (*64*, *65*) leaves open many questions about the ecological footprint of RNA viruses. Even after assuming acute infection for all RNA viruses, hence suppressing the ecological roles of their hosts, the effects on the carbon cycling often remain complex to assess. Plant RNA viruses could likely reduce photosynthetic primary production and increase dead plant material, thus decreasing CO_2_ absorption and increasing respiration by saprotrophs. However, given the diverse ecological roles of fungi, the impact of their RNA viruses across the three habitats could range from decreased CO_2_ levels by suppressing phytopathogenic and saprotrophic fungi in litter and peatlands to reduced CO_2_ assimilation due to hampered mycorrhizal fungi. Furthermore, the higher prevalence of RNA viruses infecting potential grazers in bogs and fens could reduce pressure on bacteria and archaea, thereby enhancing the mobilization of carbon from peatlands into CO_2_ and CH_4_ in these wetter habitats (*66*).

## Auxiliary metabolic genes of Stordalen Mire RNA viruses likely suppress ecological activities performed by their hosts via molecular shutoff of gene expression

Another axis for understanding the ecological impact of RNA viruses would be to assess how they might metabolically reprogram their hosts during infection. While impossible to assess in complex natural communities at any relevant scale, one window into the metabolisms viruses could impact during infection include auxiliary metabolic genes, or AMGs (*67*). While well-established for dsDNA viruses (*17–19*), previously identified AMGs in RNA viruses included diverse protein domains and motifs linked to both metabolite-transforming and information-processing pathways, requiring functional annotation with multiple databases for accurate detection. To this end, we annotated amino acid sequences encoded by RNA virus contigs using multiple databases (NCBI CDD, NCBI NR, KEGG, UniRef90, PFAM, dbCAN, MEROPS; see Methods) and revealed 52 AMGs across 51 RNA virus contigs longer than 500 nts, and 33 AMGs across ∼1.2% of the vOTUs (*n*=32) (**S8 Table**). As previously reported, AMGs encoded cellular-like protein domains and motifs (instead of complete ORFs), likely shaped by purifying selection on RNA virus genomes (facing strong streamlining constraints) after acquiring AMGs through illegitimate recombination with host mRNA (*21*, *34*). As a control, we searched the vOTU sequences against co-sampled cellular-fraction metagenomics, short-reads and long-reads, found no hits, which we interpret as consistent with the genes being RNA virus derived rather than from host genomes or endogenous virus elements (EVEs).

What kinds of AMGs were observed? Functional annotation revealed AMG-encoded proteins were associated with transcription, translation, posttranslational modifications, membrane proteins, and miscellaneous metabolic pathways (**Fig. 5**), with many functions similar to those for RNA viruses from oceanic plankton and land invertebrates (*22*, *24*, *68*). Closer examination of the AMG sequences revealed that AMGs encoded protein domains (e.g., NADAR) and motifs (e.g., motifs for SUMOylation), which could indicate the selection of specific catalytic activities with cellular proteins (**S8 Table**). To explore deeper into the effects of these AMGs on the host cells, we identified a conservative subset of 16 AMG-encoded protein sequences (indicated with double asterisks in **Fig. 5**; **S8 Table**) that were not broken by contig ends and had alignments with cdd profiles of cellular domains.

**Figure 5.**
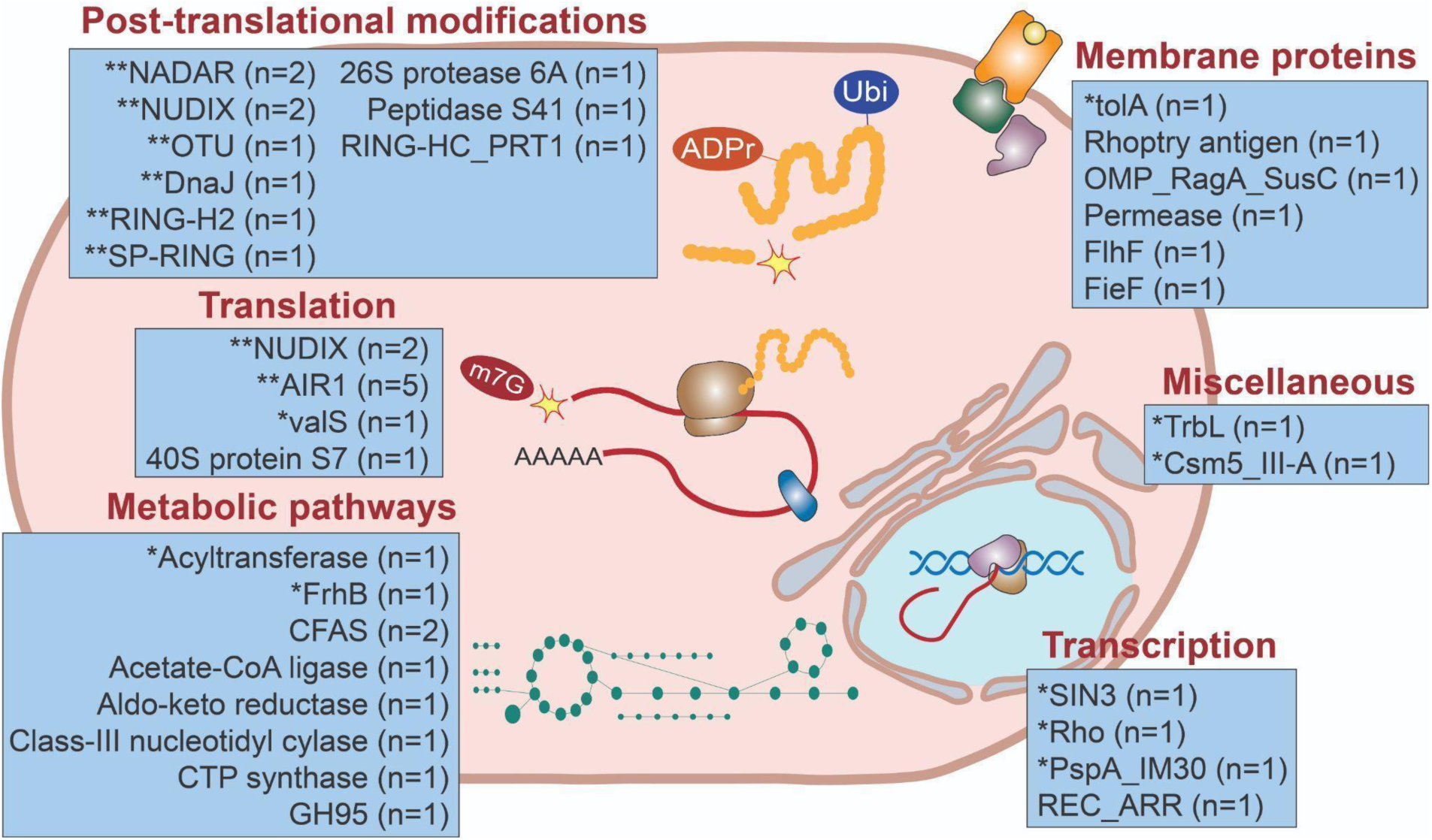
Functional diversity of AMGs carried by Stordalen Mire orthornaviraens. Schematic representation of the hypothesized effects of roles played in the manipulation of host metabolism by orthornaviraen AMGs, which are separated according to functional categories (red text) predicted from using solely functional domains (single asterisk, “permissive” set) and motifs within domains as well (double asterisk, “conservative” set). Blue boxes contain the corresponding protein annotation based on the conserved domain database (CDD), with the corresponding number of vOTUs listed in parentheses. Double asterisks indicate that AMGs that were not located at the contig ends and for which function could be identified studying motifs on the CDD, whereas those with single asterisks indicate that AMGs were not located at contig ends and for which functions could not be identified based on motifs. The absence of an asterisk indicates that AMGs were considered “broken” because given that they are located at contig ends, limiting our ability to predict their functions. AMGs that were found in viral contigs of 500–1,000 nt (**Table S8**) are not reported in this figure. RING-HC_PRT1, RING finger, HC subclass, found in thale cress (*Arabidopsis thaliana* (L.) Heynh.)) proteolysis 1 protein; valS, valine-tRNA ligase; FrhB, coenzyme F420 hydrogenase subunit beta; CFAS, cyclopropane fatty acid synthetase; GH95, glycoside hydrolase family 95; TolA, Tol proteins necessary for the uptake of all the group A colicins; TonB-linked outer membrane protein SusD/RagB family; FlhF, flagellar biosynthesis protein; FieF, divalent metal cation (Fe/Co/Zn/Cd) transporter; TrbL, conjugal transfer protein TrbL; Csm5_III-A, CRISPR system type III-A Csm effector Csm5; SIN3, transcriptional regulatory protein SIN3; Rho, transcription termination factor Rho; PspA_IM3, PspA a protein that suppresses σ^54^-dependent transcription; REC_ARR, phosphoacceptor receiver domain of type B Arabidopsis response regulators; NADAR, *Escherichia coli* swarming motility protein YbiA and related proteins; NUDIX, hydrolases of Nucleoside Diphosphate linked to another moiety (X); AIR1, arginine methyltransferase-Interacting RING finger protein.

Focusing just on these conservative 16 AMGs, activities predicted from the specific motifs found suggested translational and posttranslational interferencemechanisms as follows. *First*, five RNA viruses that encode proteins with two RNA-binding zinc knuckles (motifs of the domains AIR1 and UMSBP), that would possibly block host mRNA translation to redirect it toward virus RNA translation. Given that the only putative host of these viruses is a trypanosomid, suppression of gene expression by blocking translation in infected cells would hamper their parasitic activity on invertebrate hosts, which could increase grazing on bacteria and archaea (and hence, decrease microbial respiration-derived CO_2_ and methanogenesis) in fens. *Second*, five viruses encoded specific motifs within the domains DnaJ (HSP70-interacting motif (*69*, *70*)) and Siz/PIAS RING (CHC2- and C3H2C3-type motifs (*71*)), which putatively increase protein turnover via ubiquitination and SUMOylation, likely providing amino acids for building virus proteins. Although hosts (hymenopterans) could be assigned only to one virus, unbalanced increase of protein degradation over synthesis would lead the host cell to a catabolic state, hence diminishing the associated ecological roles. *Finally*, seven viruses encoded NUDIX hydrolases, and two viruses encoded NADAR. Both domains have been proposed to counter the antiviral defense based on NAD^+^-dependent ADP ribosylation of virus macromolecules (*72*): NUDIX cleaves the required NAD^+^ for this system and NADAR deconjugates ADP-ribosyl moieties from virus proteins and RNAs. Only one virus-encoding NUDIX was assigned to a specific host taxon (dinoflagellates of the order Suessiales). Depletion of NAD^+^, a cofactor required in many essential cellular processes, would disrupt the functions of dinoflagellates (photosynthesis and grazing), with concomitant impact on biogeochemical cycling by decreasing CO_2_ absorption or increasing CH_4_ methanogenesis. Together, these interference mechanisms boost virus fitness while decreasing host fitness with subsequent suppression of their ecological roles in the Stordalen mire with diverse impacts on carbon cycling. In summary, Stordalen More RNA viruses likely impact ecosystem processes through infection of biogeochemically important microbial eukaryotic hosts and AMGs that specifically target processes underpinning the cellular physiology that dictates biogeochemical cycling. However, quantifying these impacts awaits perturbation experiments and/or modeling designed leveraging these initial empirical observations.

## CONCLUSIONS

Our study provides a baseline for understanding ecological patterns and the footprint of soil RNA viruses in thawing permafrost, emphasizing their potential implications amid Arctic environmental and climate change. By exploring diversity patterns, we posit a near-future scenario in which permafrost thawing and proliferation of palsas will conceivably activate and expand the existing RNA virus communities, leading to increased diversity and complexity compared to current levels. We evaluated the potential roles of RNA viruses to modulate the mineralization rates of peatland carbon into CO_2_ and methane through infection. We suggest that pathogenesis and killing along with AMG-driven suppression of gene expression by RNA viruses have the potential to impact greenhouse gas emissions. Fungal hosts assume extremely diverse ecological roles, from being saprotrophs feeding on plant litter or peatland to biotic interactions (e.g., being parasites of plants or invertebrates and mutualists in lichens). Infection of saprotrophic, parasitic, and mutualistic fungi and heterotrophic protists has the potential to increase CO_2_ emissions from peatland. It could also increase methanogenesis by infecting (*i*) grazers (e.g., priapulids, nematodes, arthropods, ciliates), (*ii*) invertebrates stimulating microbiota, and (*iii*) anaerobic protists and ciliates (in particular, those associated with methanogenic archaeal endosymbionts). The manner, intensity, and direction of such effects in diverse hosts need to be addressed to allow the enable footprint of RNA viruses to be integrated into predictive ecological models. This knowledge will empower us to improve the accuracy of ecosystem models and improve the prediction of the ecological implications of climate change.

## ACKNOWLEDGMENTS

This material is based upon work supported by the U.S. Department of Energy, Office of Science, Office of Biological and Environmental Research, under Award Number DE-SC0023307 to M.B.S. Funding was provided by DOE grants to MBS. This research is a contribution of the EMERGE Biology Integration Institute, funded by the National Science Foundation, Biology Integration Institutes Program, Award #2022070. The authors thank the Swedish Polar Research Secretariat and SITES (Swedish Research Council grant 4.3-2021-00164) for the support of the work done at the Abisko Scientific Research Station. This work was supported in part through a Laulima Government Solutions, LLC, prime contract with the U.S. National Institute of Allergy and Infectious Diseases (NIAID) under Contract No. HHSN272201800013C. J.H.K. performed this work as an employee of Tunnell Government Services (TGS), a subcontractor of Laulima Government Solutions, LLC, under Contract No. HHSN272201800013C. K.C.W. funding was provided by EMERGE (#2022070), NSF Cyverse [#2149505, Collaborative Research: Updating iVirus -- the CyVerse-powered analytical toolkit for viruses of microbes], and was partially funded by the Genomic Science Program of the United States Department of Energy Office of Biological and Environmental Research, grants #DE-SC0023456.

We thank Anya Crane (National Institutes of Health National Institute of Allergy and Infectious Diseases [NIAID] Division of Clinical Research [DCR] Integrated Research Facility at Fort Detrick [IRF-Frederick], Frederick, MD, USA) for editing the manuscript. The views and conclusions contained in this document are those of the authors and should not be interpreted as necessarily representing the official policies, either expressed or implied, of the U.S. Department of Health and Human Services or the institutions and companies affiliated with the authors, nor does mention of trade names, commercial products, or organizations imply endorsement by the U.S. Government.

KK, and A.A.P is currently funded by the Deutsche Forschungsgemeinschaft (DFG, German Research Foundation) under Germany’s Excellence Strategy – EXC 2051 – Project-ID 390713860. G.D.H is currently funded by the Spanish Ministry for Ecological Transition and the Demographic Challenge.

## Data availability

The sequencing data described in this article, including metagenomic and metatranscriptomic data also described in Woodcroft et al., Emerson, et al, Sun et al. and McGivern et al. (*5*, *10*, *17*, *19*), are submitted under NCBI BioProject accession number PRJNA386568. The metagenomics assembled genomes (MAGs) used in this study were acquired from a previous study (Woodcroft et al.,(*5*)) and deposited at DDBJ/ENA/GenBank under the accession numbers provided in **Table S6**.

## SUPPLEMENTARY

### Methods

#### Sampling, purification of nucleic acids, library preparation, and short-read sequencing

A total of 55 metatranscriptomics were used in this study (**S1 Table**). Twenty-eight metatranscriptomics were collected during 2010, 2011, and 2012 (*5*, *11*); and 27 metatranscriptomics samples were collected in 2016 (*10*) (≈0.63 terabases [Tb] of data). A detailed description of active permafrost sampling strategies and protocols was previously described (*5*, *11*). Briefly, triplicate samples were taken from the active layer across the thaw permafrost gradient comprising palsa (intact permafrost), bog (partially thawed), and fen (fully thawed). The soil cores were taken at different depths of 2.5–48.5 cm (**S2 Table**). Soil cores were kept at saturation with three volumes of LifeGuard solution (MoBio Laboratories) and stored at −80°C until processing.

The protocols for extraction and purification of RNA for 2010–2012 samples were described previously (*5*). The total RNA of samples was subjected to removal of residual DNA using DNAse I (Roche) following the manufacturer’s instructions. Next, ScriptSeq Complete (Bacterial)-low input kits (Epicentre) were used for library preparation. To check the quality of RNA and libraries during processing, Agilent 2100 Bioanalyzer and Agilent 2200 Tapestation (Agilent Technologies) with Qubit (ThermoFisher Scientific) were used. Before the deeper NextSeq (Illumina) sequencing, the resulting metatranscriptomic libraries were assessed on either a HiSeq (Illumina) or MiSeq (Illumina) to evaluate library quality. For 2016 samples, the TruSeq Stranded mRNA Library Prep (Illumina cat. No. 20020594, 20020595) was used for library preparation. Using this, polyA-containing mRNA molecules were purified in two rounds with oligo-dT magnetic beads. During the second elution process of the polyA RNA, the RNA was fragmented and primed for cDNA synthesis.

#### Metatranscriptome assembly, virus identification, and evaluation of completeness of putative RdRps

The description of the metatranscriptome read assembly, virus identification, and taxonomic classification methods have been detailed previously (*21*, *22*). Briefly, the raw-read RNA sequences downloaded from the National Institutes of Health (NIH) National Center for Biotechnology Information (NCBI) Sequence Read Archive (SRA) database (https://www.ncbi.nlm.nih.gov/sra) were first processed to eliminate adapter and low-quality sequences by using Trimmomatic v0.36 (*73*) with default parameters. PhiX contamination was also removed by BBDuk (bbduk.sh; ktrim=*r*, minlen=40, minlenfraction=0.6, mink=11, tbo tpe k=23, hdist=1, hdist2=1, ftm=5). Next, *de novo* assembly was performed using MEGAHIT v1.1.3 (*74*) with default settings. To avoid problems associated with alternative genetic code and non-canonical translation events, EMBOSS Transeq was used (instead of protein prediction with prodigal) to yield RdRp footprints.

Approaches to identify the RNA virus genomes or transcripts from assembled metatranscriptomes and the authentication of virus RdRp were described previously (*22*). To improve the identification of the divergent RdRp, the Hidden Markov Model (HMM) was applied over 10 iterations to search for virus RdRp domains (**fig. S1**). To evaluate the authenticity, thus true positive hits of virus RdRp sequences, competitive HMM searches were conducted of 70 RdRps derived from the lineages defined previously (*24*), eight recently discovered lineages (*22*), and 16,268 PfamA (v33.0) non-RdRp HMMs. Further, the putative RdRp hits with the best match to an RdRp HMM and a bit-score ≥30 was kept as true positives for proteins containing the virus RdRp domain. In addition, to confirm that the RNA virus contigs were not from endogenous viruses, the identified sequences were searched against the available metagenomes.

To estimate RdRp domain completeness, based on previous efforts (*22*), the extraction of footprint amino acid sequences of the RdRp domain was performed considering the challenging cases of the usage of alternative genetic codes and non-canonical translation events (e.g., ribosomal frameshifting). Extraction and reconstruction of RdRp domain sequences were performed by running another iteration of the search for the translated amino acid against the 70 RdRp profile HMMs, and the hits with a bit score ≥30 and with aligning regions of ≥90% of the average length of the best-matching HMMs (to be considered as “complete” or “near-complete” virus RdRp domain sequences). The 3,560 extracted RdRps that were shorter than 100 amino acids were further inspected for RdRp annotations, having detected only 18 of them for which no virus signal was apparent, hence further supporting that 99.5% of hits are RdRp-encoding sequences derived from RNA viruses. Lastly, to estimate genome completeness for the recovered viral contigs, CheckV (*75*) was used with its default parameters (**fig. S2**), which compared amino acid identity and HMMs of complete viral genomes derived from NIH NCBI GenBank and environmental samples.

#### Virus operational taxonomic units (vOTU) and calculations of vOTU relative abundances and diversity metrics

To analyze the diversity and ecological pattern of soil RNA viruses, they were first designated as vOTUs or an approximate “species-rank” ecological unit; the detailed analysis of the RNA virus sequence space to establish a “universal” cutoff has been reported previously (*21*, *22*). Using the previously recommended cutoff, 90% average nucleotide identity (ANI) across 80% of the shorter sequence length (*21*, *22*), resulted in 9,567 total vOTUs, of which 2,561 were ≥1 kb (**S3 Table**). Further, to assess the robustness of this cutoff in the soil system, an investigation of the ecological patterns was performed across an ANI range of 60–100% (and 80% allele frequency [AF]), including consensus cutoff recommended by (*33*), 95% ANI and 85% AF (**fig. S3**).

To generate the relative abundance table of the designated vOTUs, the previously described approach was followed (*21*, *22*). Before the mapping, virus contigs were trimmed off their polyA/polyT sequences; this was done to avoid inflated abundance and better estimate horizontal coverage for polyA-tailed viruses, respectively. Further, a quality-controlled sequence read of metatranscriptomic samples to the vOTUs was recruited, and trimmed mean abundances were calculated with CoverM (https://github.com/wwood/CoverM). The trimmed mean abundance table was then normalized by dividing abundances by the number of quality-controlled reads per sample per 1 Tb to get the final relative abundance table (**Table S4**).

The ecological analysis was done using samples collected in 2012 and 2016 to minimize bias introduced by samples variably from 2010 and 2011. Different diversity metrics were calculated, including α-diversity (Shannon’s H, Simpson, Inverse–Simpson, Richness, and Perilous J Evenness) and β-diversity (Bray–Curtis dissimilarity method), using R vegan package (*76*). vOTUs that were not present in any samples and samples with no vOTUs present were excluded. The abundance table of vOTUs was log-transformed (function ‘decostand’, method = ‘log’) before the calculation of the Bray–Curtis dissimilarity analysis (function ‘vegdist’, method ‘bray’). This was then used to generate principal coordinate analysis (pCoA, function ‘capscale’ of the vegan package). The emerging metacommunities in pCoA were statistically verified using permanova tests (function ‘adonis’) and the confidence intervals were plotted using the standard deviation method (function ‘ordiellipse’) at 95 and 97.5% confidence limits. The same Bray–Curtis dissimilarity matrices were used for hierarchical clustering (function ‘pvclust’, method.dist =‘’co’’ and method.hclust = ‘average’) using 1,000 bootstrap iterations. Only the approximately unbiased (AU) bootstrap values were reported on the dendrograms. The heatmaps of the hierarchical clustering analysis were generated using the heatmap3 package (*77*) (**fig. S3–S5**).

#### Comparing Stordalen Mire RNA virus to other soil RNA virus studies

Metatranscriptome sequences were downloaded from recently published soil RNA virus studies, which included examination of California grassland (*39*), grassland and coastal soils (*32*), crop field soil (*31*), Kansas grassland (*41*), and various soil types across China (*38*). Similar pipelines, including metatranscriptome assembly, virus identification, and evaluation of completeness of putative RdRps, were run as described above. To compare the virus sequences across different soil studies, the identified RNA virus contigs were clustered at 90% ANI and 80% AF to get vOTUs. For RdRp-level comparison, only those “complete”/“near-complete” (≥90% of the average length of the best-matching HMMs) were used. The RdRps were clustered using USEARCH (-cluster_fast) (*78*) at 75% Identity of Amino Acid (IAA), representing taxonomic rank between family and genera (*24*). Subsequently, those clustered at 75%, were considered shared among studies. These results were plotted in UpSet using the UpSetR package in R (*79*) (**fig. S6**). To compare the identified orthornavirans from this study to other soil studies, taxonomy assignment was done using RdRp-scan (*80*).

#### RdRp-based taxonomic annotation of RNA viruses

The taxonomic classification analysis was conducted using both the network-based, iterative clustering approach and the phylogenetic described previously (full details in Zayed et al., 2022) (*22*) (**fig. S7**). Briefly, a total of 705,705 complete and near-complete RdRp domains—including 1,702 from the current Stordalen Mire soil study; 6,732 from a *Tara* Ocean study (*21*, *22*); 3,325 from coastal ocean RNA viromes (*24*); and 695,648 from GenBank [version 245]—were used in the analysis. First, these sequences were pre-clustered at 50% amino acid identity using Uclust v10.0.240 (usearch -cluster_fast -id 0.50 -sort length) (*78*). Then, these centroid sequences (*n* = 6,553) were used in pairwise comparisons using blastp v2.10.0+ (-gapopen 9 - gapextend 1 -word_size 3 -threshold 10) (*81*). Further, the e-values for each pair were extracted and negative-long10-transformed in MCL v14-137 (-stream-mirror -stream-neg-log10 -stream-tf ‘ceil(200)’) (*82*). The transformed e-values were used in an MCL network for iterative clustering, changing the granularity parameter at each iteration (range 0.1–8). From this point, ‘adj.rand.index’ value was used to evaluate all the cluster sets from MCL and concurrently compared to the previously established ICTV taxonomy (**Table S5**). The clusters with the highest rand index value at the phylum and class ranks were picked and the taxonomic delineations were then assigned for sequences within each cluster.

Moreover, phylogenetic trees were generated for each assigned taxonomic network-derived cluster (at the phylum and class ranks) as established previously (*22*). Briefly, from each resultant network-derived major cluster, sequences from each cluster were aligned separately using the E-INS-I strategy over 1,000 iterations in MAFFT v7.017 (*83*). Aligned sequences were subsequently trimmed using Trimal (*84*). Sites with ≥20% gaps were removed, and the alignments were manually inspected. Before phylogenetic analysis, sequences were screened for possible recombination events using 3Seq (*85*), with a recombinant event determined by a Bonferroni-corrected *p-*value cutoff of 0.05. Phylogenetic relationships of sequences within a cluster were first assigned the appropriate evolutionary model using ModelFinder (*86*). Then, a subsequent maximum-likelihood phylogenetic tree was generated using bootstrap support generated for 1,000 iterations in IQ-TREE (*87*). Given the high number of reference sequences within the phylum-level phylogenetic trees, a subset of these trees was created to aid in visualization. Redundant reference sequences were reduced while maintaining the diversity captured in the original tree using Treemer, applying stop option -X (50–200) depending on the phylum (*88*). For **Fig. 1**, sequences used to build the trees were preclustered at 50% identity, and clades supported by 100% bootstrap values were collapsed.

To taxonomically assign sequence to a family rank, a two-pronged approach was used. First, the class-rank phylogenetic trees were used to establish the placement of sequences on the tree, requiring a sequence to fall within a clade or sister clade to be assigned to the same taxon. Then, sequences were assigned tentative names identical to the reference sequences of the established clades with a ‘-like’ suffix added. Further, the prior analysis was complemented by the taxonomic assignment from the conservative MCL clusters. Subsequently, these two approaches were combined and the more specific taxonomic assignment of the two methods (e.g., without the ‘-like’ suffix or higher-resolution taxonomic assignment) was picked as the final putative classification for the novel sequences.

#### Host inference of hosts and cellular organisms’ analysis

To infer the virus-host associations, a few independent approaches—including RdRp-based taxonomy, endogenous virus elements (EVEs), RdRp-based against the cellular non-redundant (nr) database, alternative genetic code, Shine–Dalgarno motifs, and CRISPR-Cas spacers analysis—were used. The following is the detailed approach: (*i*) The RdRp-based taxonomy approach was used to infer the potential virus–host associations. For this approach, the known RNA virus taxa (phyla, classes, and families) that were assigned to the vOTUs after clustering the RdRp domain protein sequence similarity network were used to retrieve the information on putative hosts. (*ii*) EVEs looked for closely-related RdRp-encoded cellular genomes, and the RdRp hits (from virus contigs) suggested that a virus contig can have a putative host taxonomically related to one of the cellular genomes (*68*). All available assembly genomes from cellular organisms in GenBank version 205 (last accessed May 2021) were downloaded; this included archaea (4.89 Gbp), bacteria (234 Gbp), fungi (157 Gbp), invertebrates (618 Gbp), plants (683 Gbp), protozoa (41.6 Gbp), and vertebrates (2.44 Tbp). A more sensitive diamond (version 0.9.36) blastx algorithm against the aforementioned databases was used (--evalue 1e-20–-sensitive–-out–-outfmt 6). The threshold for the search was set to 100 amino acids for length and 1 × 10−20 for e-value. The results were also checked manually to accurately assign the putative host taxonomy. (*iii*) Host information was inferred from cellular nr databases. The query of virus RdRp footprints was then used against this database. The threshold for the search was set to 100 amino acids for length and 1 × 10^−5^ for e-value. Only the top match hits were used, and the results were manually checked. (*iv*) Using genetic code, the translated sequences were searched once more against the 84 RdRp profile HMMs, and those with a bit score ≥30 and aligning regions of ≥90% of the average length of the best-matching HMMs were considered proteins containing “complete” virus RdRp domain sequences. Sequences containing more than one hit in the same or different frames (presumably non-canonical translation cases) were resolved manually by joining the aligning regions into the same protein sequence. (*v*) Shine–Dalgarno motifs, following the same approach described by Menardo et al., 2018 (*88*). (*vi*) To predict potential prokaryotic hosts using the CRISPR-Cas spacer approach, blastn was used against ≈12 million spacer sequences (*89*), and only blastn hits with 0 or 1 mismatch over the whole spacer length were considered. Furthermore, spacers were extracted from 1,529 previously described metagenomics-assembled genomes (MAGs) (*5*) using MinCed (https://github.com/ctSkennerton/minced). A minimum of two repeats was required for a CRISPR to be included in the analysis. Further, as in the preceding analysis, only blastn hits with 0 or 1 mismatch over the whole spacer length were considered. The best hit sequence alignment was then visualized using mView (https://www.ebi.ac.uk/Tools/msa/mview/). Moreover, the taxonomic assignment of the identified orthornavirans, which was performed using RdRp-scan (*80*), was also utilized to infer the hosts. Additionally, to identify the RNA virus infecting bacteria, bacteriolytics analysis was conducted using HMM searches, utilizing the HMM profiles previously provided (*34*) and only considering hits with a bit score ≥30 for further analysis (**fig. S9–S12**). Leviviricetes require bacterial pili to attach to and infect cells (*90*). Therefore, we performed competitive searches using HMMsearch with nine type-IV pilus (T4P) profiles (PF13681.9, PF11104.11, PF16074.8, PF17456.5, PF11773.11, PF18222.4, PF04350.16, PF04351.16, and PF04964.17) and 13 non-T4P profiles containing the keyword “pilus” (PF07434.14, PF09589.13, PF14855.9, PF09476.13, PF16976.8, PF09460.13, PF13629.9, PF13400.9, PF16968.8, PF09977.12, PF05946.15, PF06340.14, and PF10671.12). The best match to a T4P HMM profile and a bit score ≥50 were considered true positives for proteins containing the bacterial T4P domain. To ensure a true positive match, random hits were manually checked across the datasets with BLASTP using RefSeq and nr databases. The integrated host interference analysis can be seen in **S7 Table**.

Next, to get a more comprehensive picture of the eukaryotic community and due to the lack of this information in Stordalen Mire, a further investigation focused on cellular organism community structure using HMM marker genes—cytochrome c oxidase subunit 1 mitochondrial gene (*COX1*); a component of the eukaryotic 40S ribosomal subunit, ribosomal protein S3 (*RPS3*) (*39*); and small subunit ribosomal RNA (SSU rRNA). For *COX1* and *RPS3* HMM profiles were used to screen the metatranscriptomic data. Hits with an e-value = 10–5 and a bit score ≥30 were extracted, and the top match for each marker gene was retained. The *COX1* and *RPS3* sequences were dereplicated and clustered at 98 and 99% amino acid identity, respectively, using USEARCH (-cluster_fast) (*78*). The reconstruction analysis was then performed for the second time by running the translated contigs against the marker genes HMMs. The hits with a bit score ≥30 and aligning regions of ≥90% of the average length of the best-matching HMMs (to be considered as “complete” or “near-complete” virus RdRp domain sequences) were retained. Further, a manual check was also done to verify the taxonomy of the identified community. Additionally, SSU rRNA—including 28S and 18S (eukaryotes); 23S, 16S, and 23S (bacteria); and 16S (archaea)—were extracted from assembled contigs and subsequently blasted against the Silva database (*91*) to determine their cellular organism taxonomy. The vOTUs of the cellular organisms were then extracted from the aforementioned dereplicated marker genes.

#### Ecological drivers’ analysis

To examine the relationships between the relative abundance of orthornaviraens in different habitats and environmental factors, a redundancy analysis (RDA) was performed using the vegan package in R (*76*) and Hellinger. To ensure a suitable ratio of explanatory variables to samples, variance inflation factors (VIF) were used and included soil depth (avg.), air temperature (°C), soil temperature (°C), porewater pH, carbon dioxide (CO_2_; mM), methane (CH_4_; mM), carbon (C; %), nitrogen (N; %), δ^13^C of CO_2_, δ^13^C of CH_4_, δ^15^N of peat, relative abundance of cellular organisms (*COX1* and *RPS3*), and relative abundance of cellular organism (SSU rRNA) in the analysis.

The environmental variables were fitted to the first two dimensions of the PCoA using a generalized additive model (function envfit; permutations = 9,999 and na.rm = TRUE) and R vegan package (*76*). Then, they were correlated with all the pCoA dimensions using a mantel test (function mantel; permutations = 9,999 and method = “spearman”) after scaling (function scale) and calculating their distance matrices (function vegdist; method = “Euclidean” and na.rm = TRUE). Furthermore, Spearman’s correlation analysis was performed (function cor; use = “pairwise.complete.obs”) and plotted using ggplot2. Finally, Spearman’s pairwise correlation was performed with vegan package in R. The “rcorr” function and the Hellinger transformed data were used as input. Correlograms were generated using the corrplot package to visualize the correlation results; only variables with significant correlations highlighted at a significance level of 0.05 were shown.

To quantify the relative contributions of environmental variables to the changes in virus community (the first coordinate of a pCoA; pCo1), a relative-importance analysis test was performed. This analysis utilized the “relaimpo” v2.2–348 package (*92*), using commonly used multiple-linear regression models (i.e., LMG, First, Last, Betasq, and Pratt) with 1,000 bootstraps. The relaimpo package provided methods for estimating the contributions of explanatory variables within a multi-linear regression model. The Lindeman–Melinda–Gold model (LMG) calculated the marginal contribution of each predictor variable while accounting for the presence of all other predictor variables in the model. The model “last” (also called usefulness) calculated the importance of each variable as the reduction in the residual sum of squares (RSS) when that variable was added last to a model that already contained all the other predictor variables. The model “first” calculated the importance of each variable as the reduction in the RSS when all the other predictor variables were added to a model that already contained that variable. Model “Betasq” calculated the importance of each variable as the square of its standardized regression coefficient. Model “Prat” calculated the importance of each variable as the sum of squared correlations between that variable and all other variables in the model (**fig. S16**).

#### Functional annotation of RNA virus genomes

Functional aspects of the protein sequences encoded by RNA virus nucleotide sequences (partial or complete genomes and transcripts) at 95% ANI and 80% AF were studied by integrating annotations from several tools. Nucleotide sequences of contigs coding RdRp were searched against (*i*) the conserved domain database (CDD) by using CDD search (*93*), HMMER (*94*), and a minimum e-value of 0.01 and (*ii*) against the NCBI nr database using blastx and a minimum bit score of 50. Protein sequences predicted from the RdRp-coding contigs by using Prodigal (apply corresponding genetic codes previously inferred) were (*i*) searched against KEGG, UniRef, CAZY, MEROPS, Pfam, and VOG databases by using DRAM (*95*). Virus coding sequences whose functional annotation suggested a “cellular” origin were considered “auxiliary metabolic genes” (AMGs) after manual inspection if none of the following cases was accomplished: (*i*) the putative cellular domains overlapped typical RNA viral domains (e.g., RdRp, capsid, helicase, protease), (*ii*) the putatively cellular domain was equivalent to known RNA viral domains, (*iii*) blastx hits also included viral proteins (to avoid poor annotations in the NCBI nr database), or (*iv*) blastx hits were not hypothetical proteins of unknown functions. To determine redundancies and avoid different annotation names for the same gene function, AMG sequences not at the contig ends were considered “complete” and compared against each other through blastp (cutoffs: e-value of 0.001, 20% IAA, and query coverage of 20%). To avoid the possibility of these AMGs being derived from sequenced genomic DNA carrying EVEs, RNA virus contigs were searched against all the available assembled and co-sampled DNA sequencing data (as described in the sections on metatranscriptome assembly, virus identification, and evaluation of completeness of putative RdRps). The final list of RNA virus AMGs can be seen in **Table S8**.

## SUPPORTING INFORMATION

**Fig. S1.**
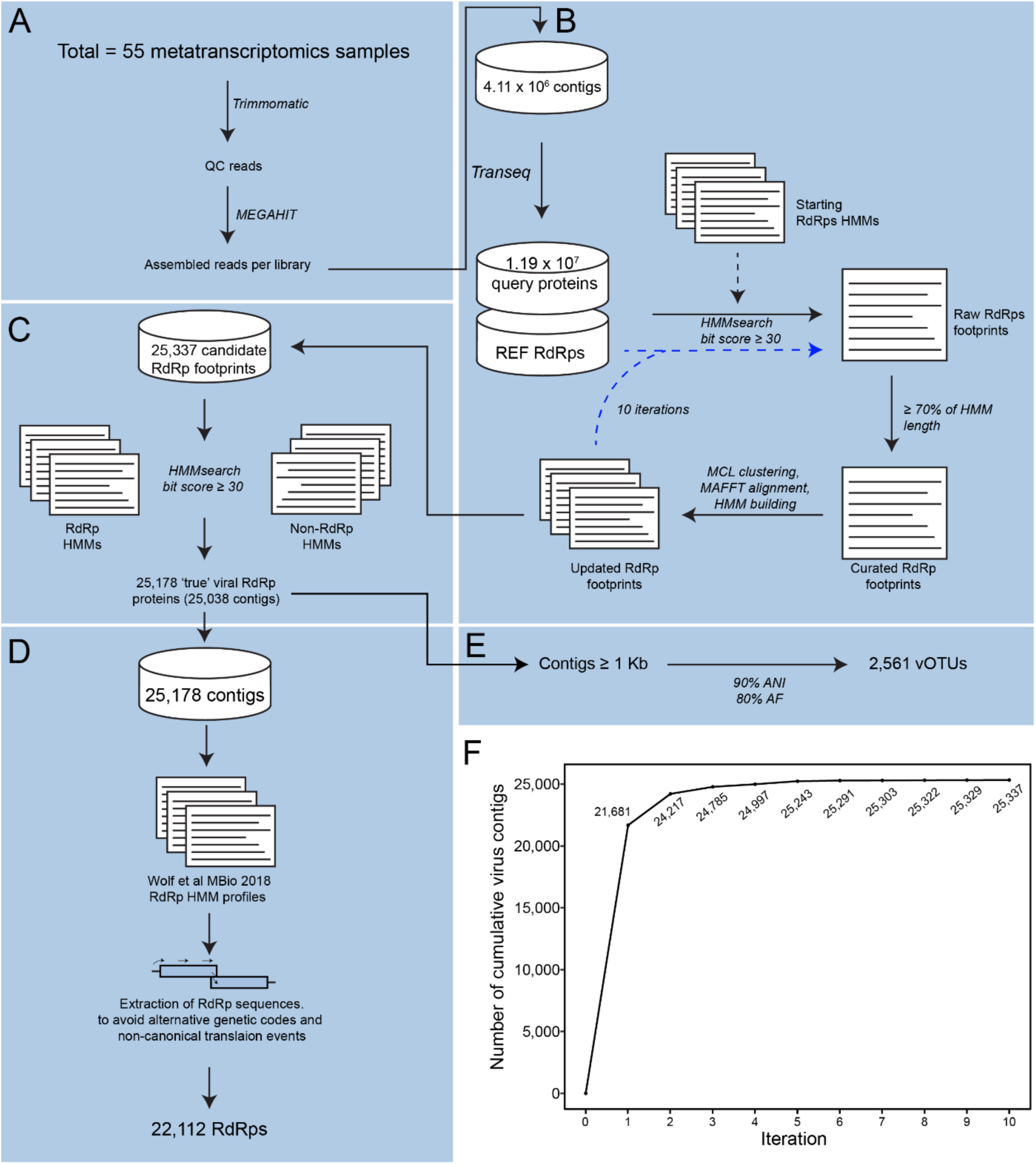
Bioinformatics workflow and Stordalen Mire orthornaviraen identification. (**A**) The metatranscriptomes from Stordalen Mire permafrost, collected in 2010, 2011, 2012, and 2016, were preprocessed, including quality-trimming and assembly into contigs. (**B**) The deduced encoded protein sequences were then used to identify orthornaviraens. To capture highly divergent RdRps, a search-and-update Hidden Markov Model (HMMs) approach was used over ten iterations (dotted blue lines). (**C**) Competitive HMM profiles were used to evaluate the authenticity of RdRp hits. (**D**) Extraction and reconstruction of RdRp domain sequences were performed by running another iteration of searching and deducing protein sequences against the RdRp profile HMMs. (**E**) For ecological analysis, virus operational taxonomic units (vOTUs), i.e., approximate species-rank ecological units, at the 90% Average Nucleotide Acid (ANI), and an 80% alignment fraction (AF) were applied. (**F**) Virus contigs were detected as a result of 10 iterations of HMMsearch.

**Fig. S2.**
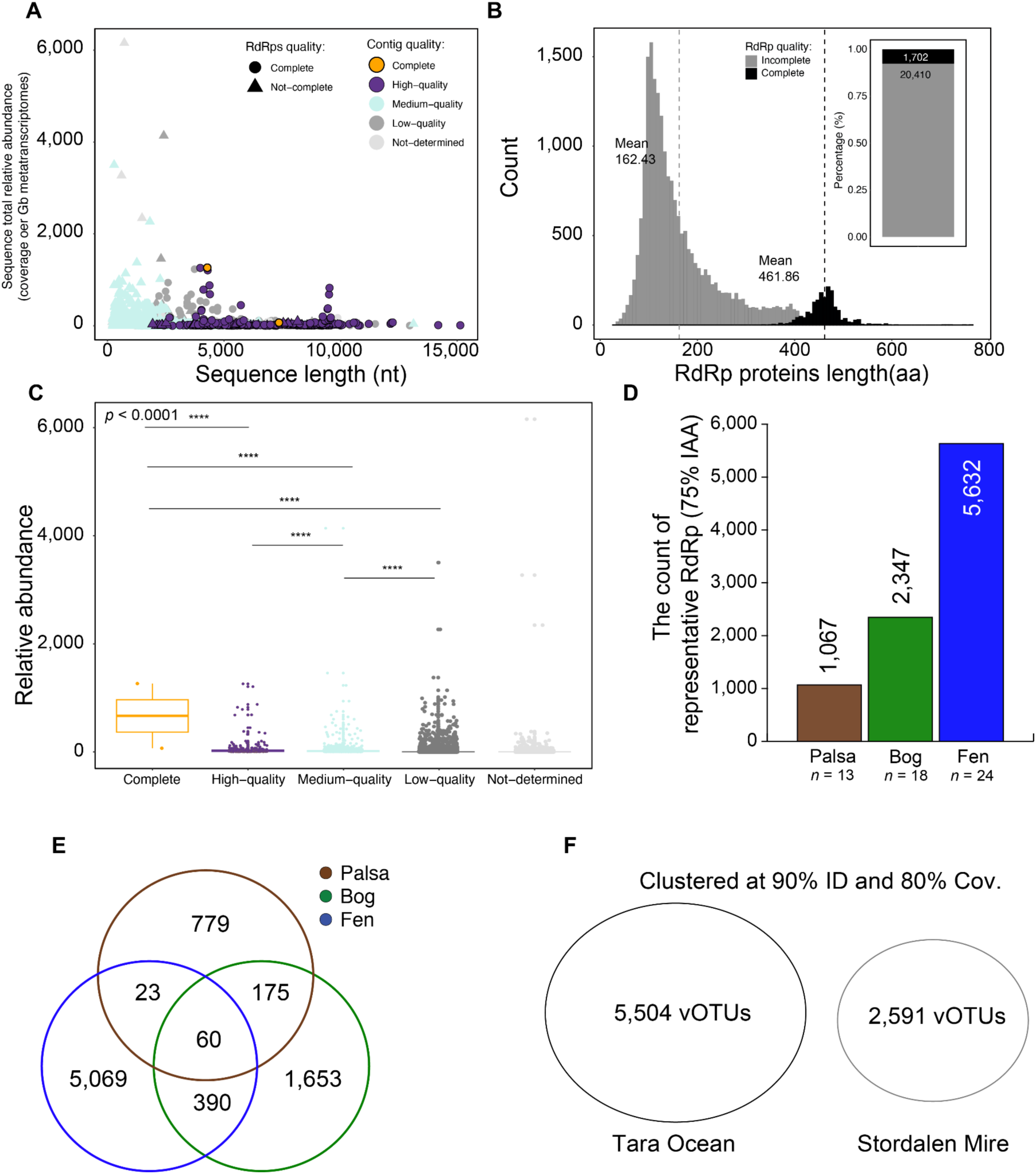
Completeness of Stordalen Mire orthornaviran genomes and RdRp domains. (**A**) Lengths of 22,112 analyzed contigs encoding virus RdRps (*x* axis) and their cumulative coverage across Stordalen Mire (*y* axis), indicating completeness of virus genome sequences and RdRp domains. Contig quality analysis was performed using CheckV. (**B**) Histogram depicting the length distribution of complete and incomplete RdRp domain sequences. The inset stacked barplot indicates the percentage of complete RdRp domains. (**C**) Stordalen Mire orthornaviran contig quality (based on CheckV). Statistical analysis was performed using Kruskal–Wallis analysis, with *post hoc* Dunn-test and *p*-adjusted: Bonferroni. Only *p*-values ≤ 0.0001 are shown (****). (**D**) Counts of RdRp representative footprints per habitat at the 75% amino acid identity (AAI) level. (**E**) Counts of RdRp footprint representatives across habitats at 75% AAI (number of samples as indicated in D). (**F**) Clustering of 5,504 *Tara* ocean RNA viruses and 2,591 vOTUs of RNA viruses from Stordalen Mire at 90% identity and 80% coverage. No overlapping of viruses at the “species-level”.

**Fig. S3.**
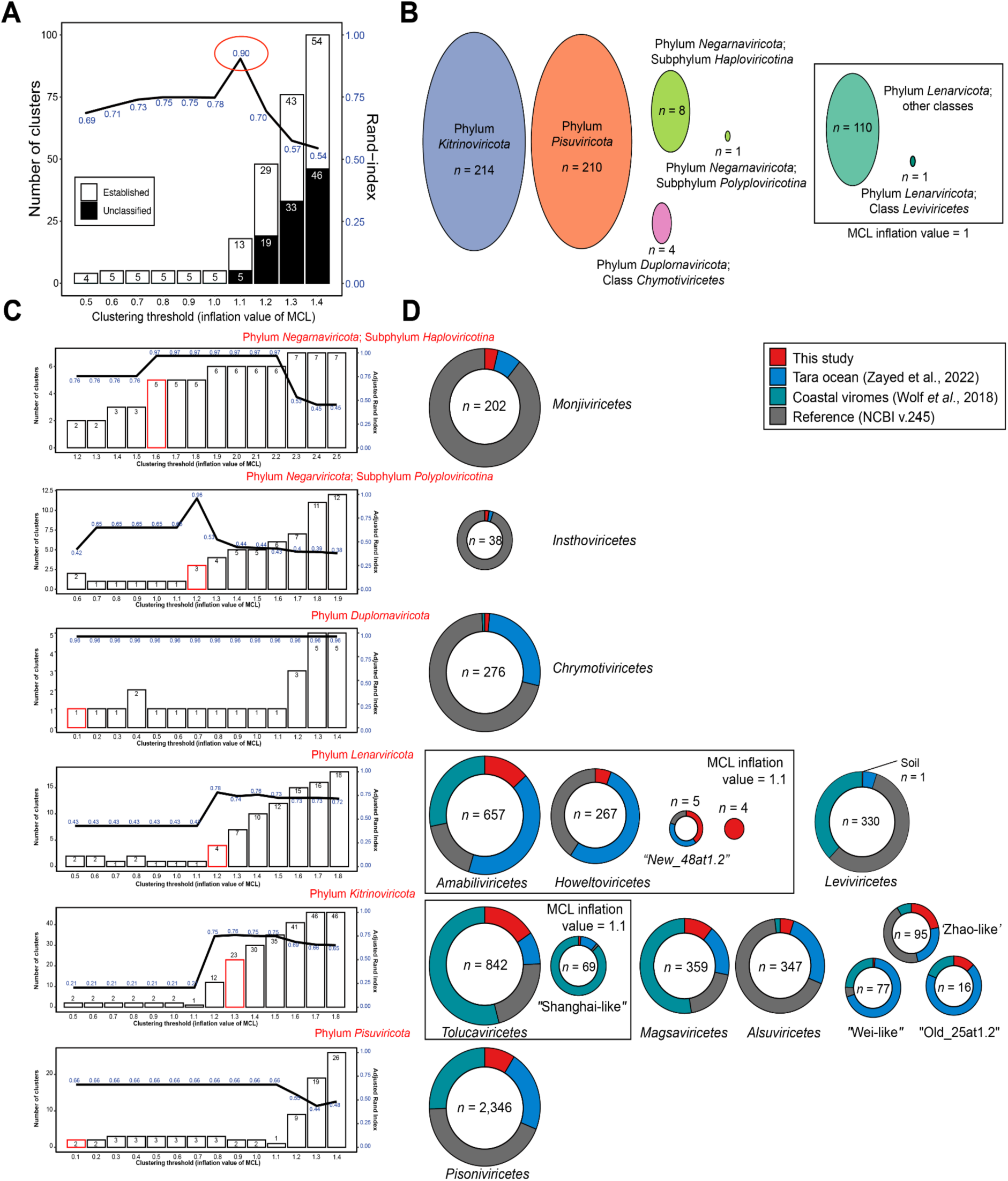
Establishment of RdRp domain-based phylum- and class-rank clusters. (**A**) adjusted rand index (line) of the network-guided and phylogeny-based megataxonomy at different clustering thresholds. Stacked bars represent the number of taxonomic clusters of near-complete RdRp domains (at least 90% of the domain) at the depicted clustering thresholds. Only sequences representing established taxa (black) were used for calculating the agreement percentage. The highest agreement is shown at an inflation value of 1.1, forming a total of 18 clustered phyla; five were previously established by Wolf et al. (**B**) Established taxa at the Markov Clustering Algorithm (MCL) inflation value of 1.1, represented by ellipses in distinct colors. The box depicts taxa that were exclusively joined at lower inflation values. (**C**) adjusted rand index (line) of the network-guided and phylogeny-based at phylum/subphylum rank. (**D**) Pie charts represent the proportion of orthornaviraens of the “complete”/“near-complete” RdRp domains (of the network-guided phylogeny analysis) at class rank clusters, including sequences from this study (red), and references (NCBI—grey, *Tara* Ocean—blue, and coastal ocean viromes—green). The box depicts taxa that were exclusively joined at lower inflation values.

**Fig. S4.**
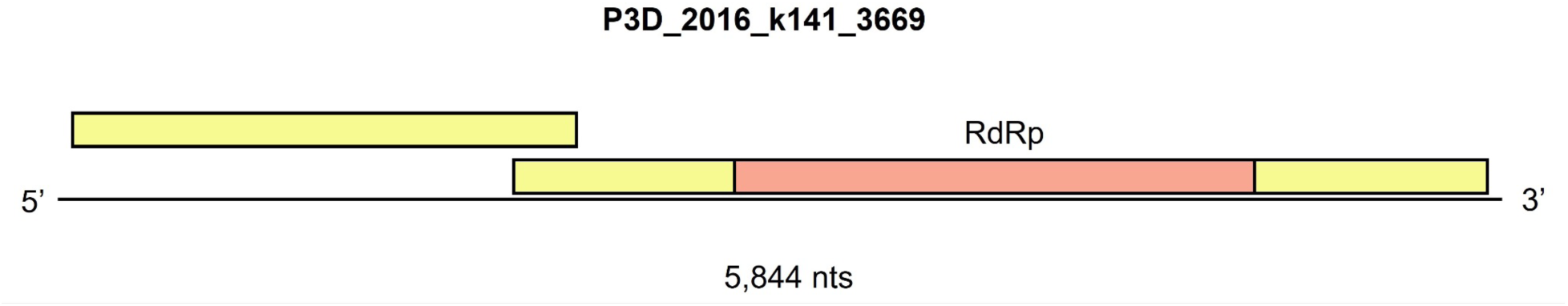
Genome of novel class ‘*Stormiviricetes*’. The longest genome of the newly identified class of ‘*Stormiviricetes*”. Yellow boxes represent ORFs in 5’→3’ sense and coral boxes represent the RdRp domain.

**Fig. S5.**
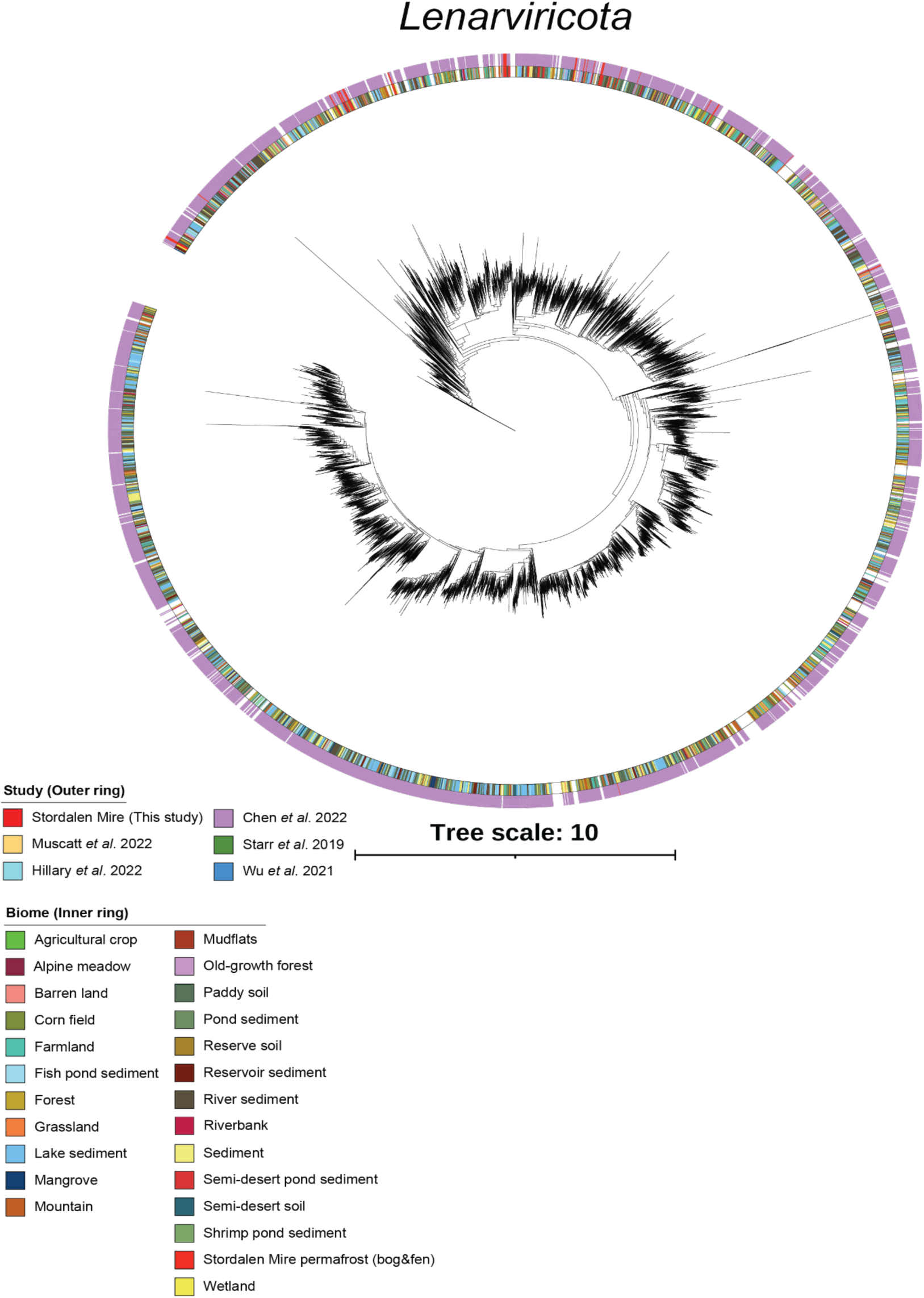
Thawing permafrost lenarviricots. RdRp-based phylogenies across RNA virus studies. A maximum-likelihood phylogenetic tree was built from the RdRp-guided taxonomy analysis of near-complete RdRp domain sequences. The scale bar indicates one amino acid residue substitution per site. Sequences used to build the trees were preclustered at 40% identity, and clades supported by 100% bootstrap values were collapsed. The inner ring represents the biomes of these viruses whereas the outer ring represents soil RNA virus studies.

**Fig. S6.**
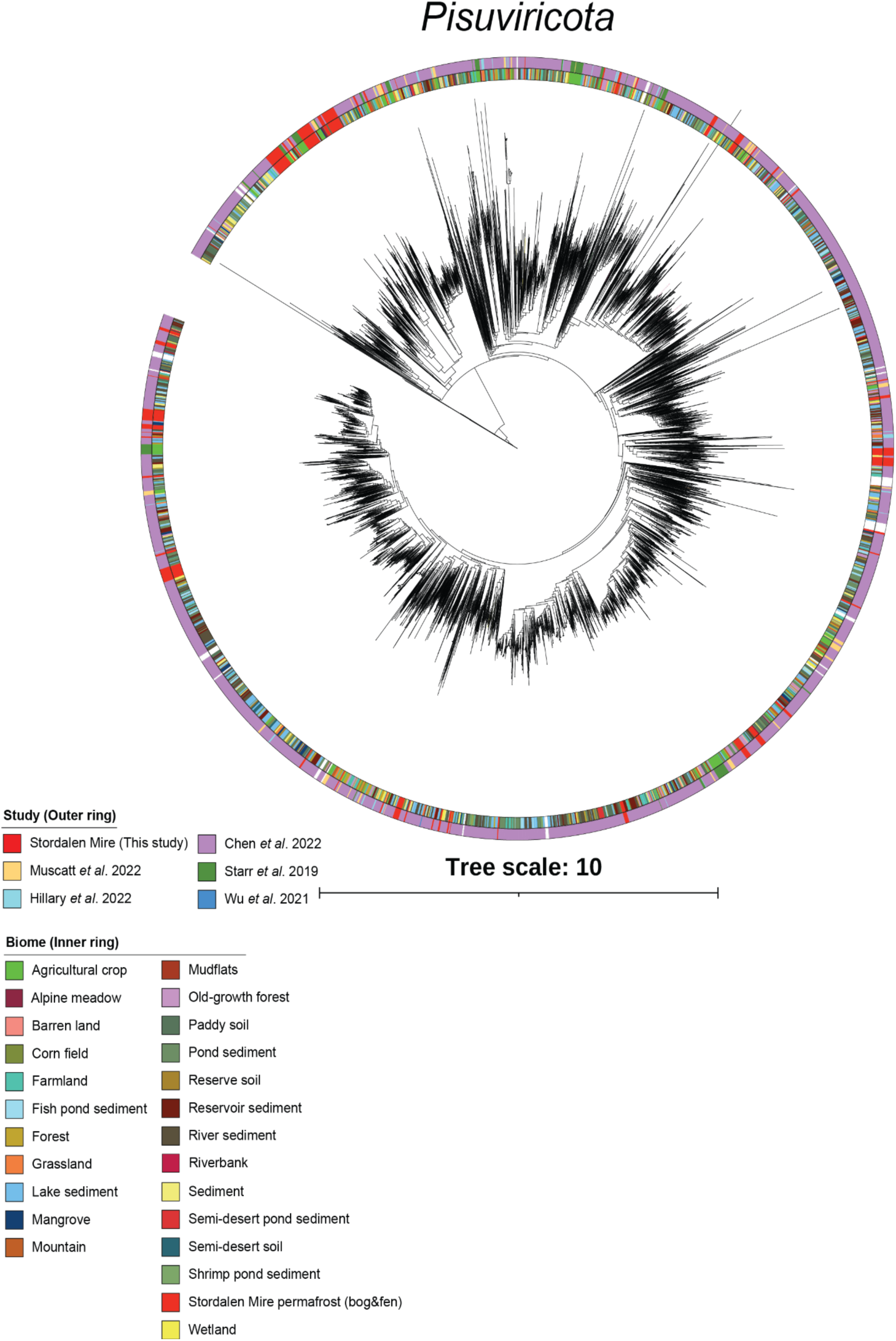
Thawing permafrost pisuviricots. RdRp-based phylogenies across RNA virus studies. A maximum-likelihood phylogenetic tree was built from the RdRp-guided taxonomy analysis of near-complete RdRp domain sequences. The scale bar indicates one amino acid residue substitution per site. Sequences used to build the trees were preclustered at 40% identity, and clades supported by 100% bootstrap values were collapsed. The inner ring represents the biomes of these viruses whereas the outer ring represents soil RNA virus studies.

**Fig. S7.**
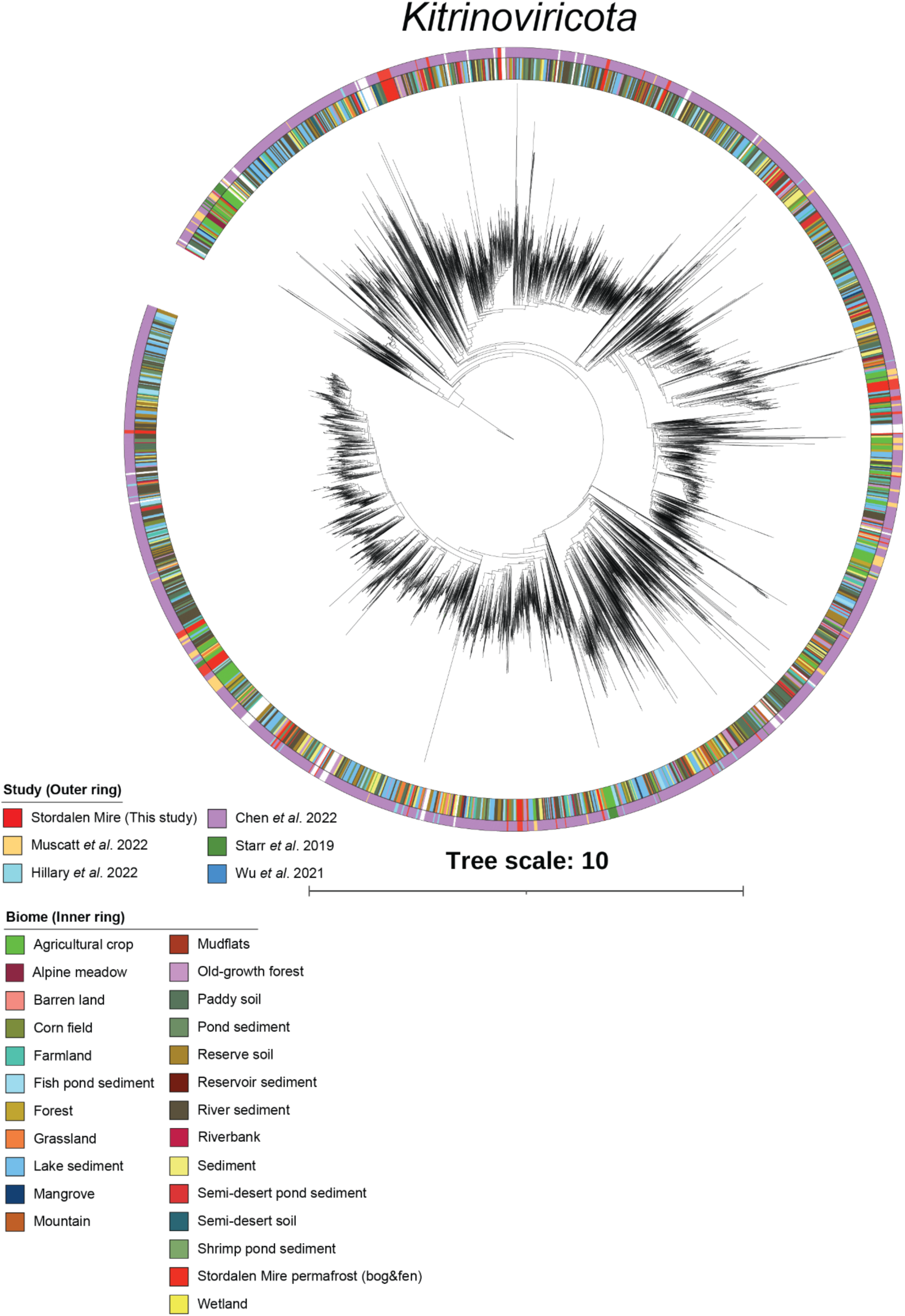
Thawing permafrost kitrinoviricots. RdRp-based phylogenies across RNA virus studies. A maximum-likelihood phylogenetic tree was built from the RdRp-guided taxonomy analysis of near-complete RdRp domain sequences. The scale bar indicates one amino acid residue substitution per site. Sequences used to build the trees were preclustered at 40% identity, and clades supported by 100% bootstrap values were collapsed. The inner ring represents the biomes of these viruses whereas the outer ring represents soil RNA virus studies.

**Fig. S8.**
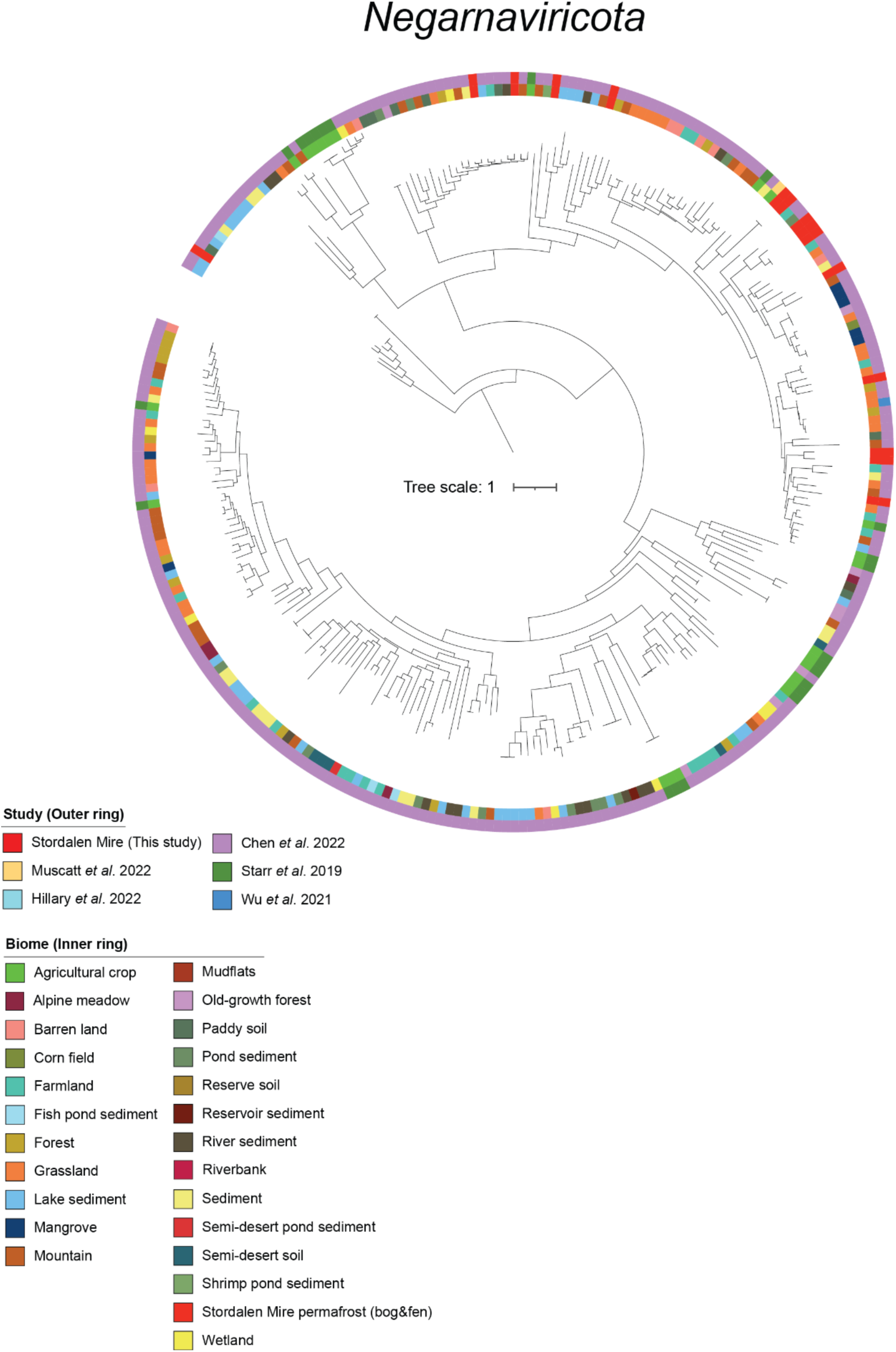
Thawing permafrost negarnaviricots. RdRp-based phylogenies across RNA virus studies. A maximum-likelihood phylogenetic tree was built from the RdRp-guided taxonomy analysis of near-complete RdRp domain sequences. The scale bar indicates one amino acid residue substitution per site. Sequences used to build the trees were preclustered at 40% identity, and clades supported by 100% bootstrap values were collapsed. The inner ring represents the biomes of these viruses whereas the outer ring represents soil RNA virus studies.

**Fig. S9.**
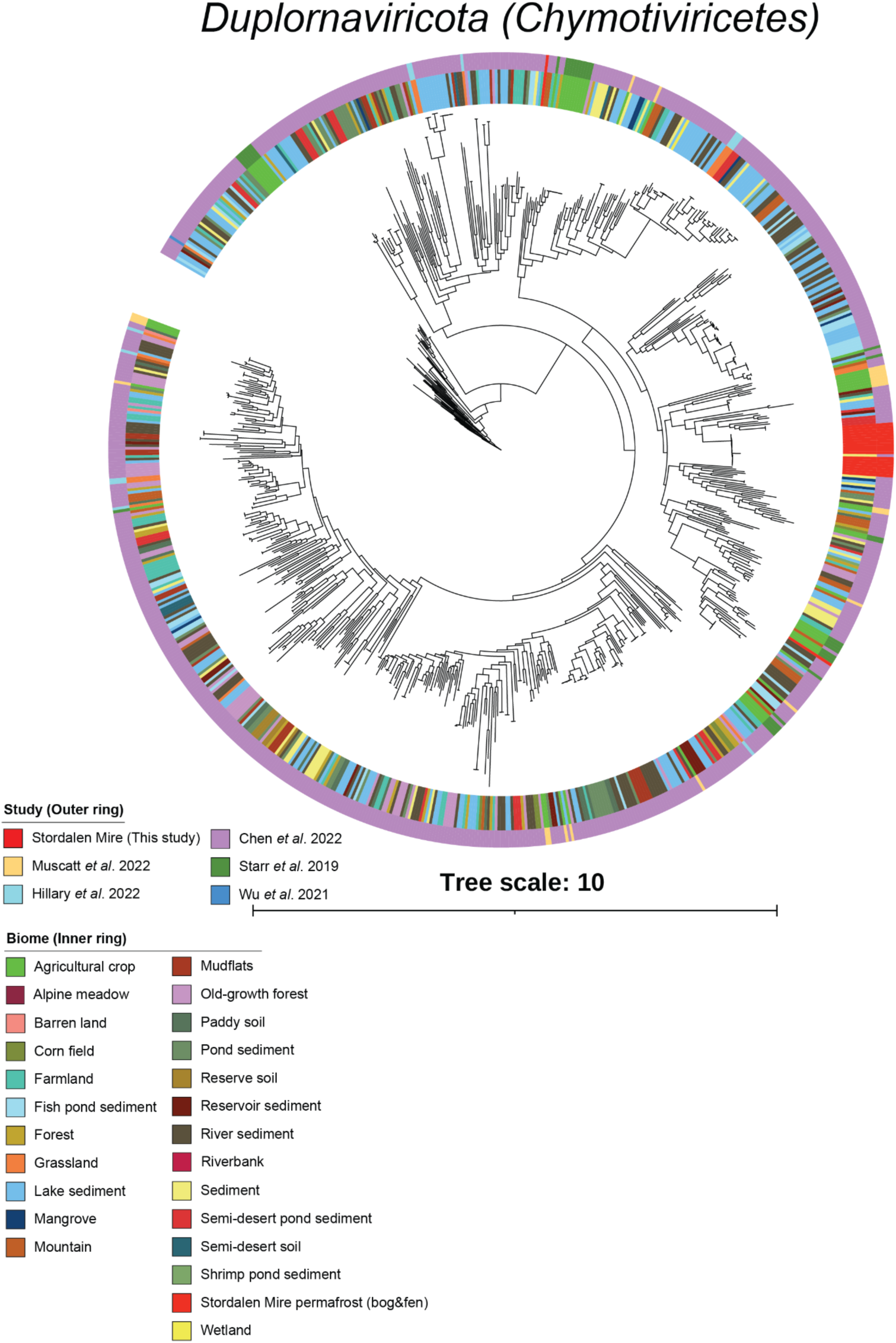
Thawing permafrost duplornaviricots (chrymotiviricetes). RdRp-based phylogenies across RNA virus studies. A maximum-likelihood phylogenetic tree was built from the RdRp-guided taxonomy analysis of near-complete RdRp domain sequences. The scale bar indicates one amino acid residue substitution per site. Sequences used to build the trees were preclustered at 40% identity, and clades supported by 100% bootstrap values were collapsed. The inner ring represents the biomes of these viruses whereas the outer ring represents soil RNA virus studies.

**Fig. S10.**
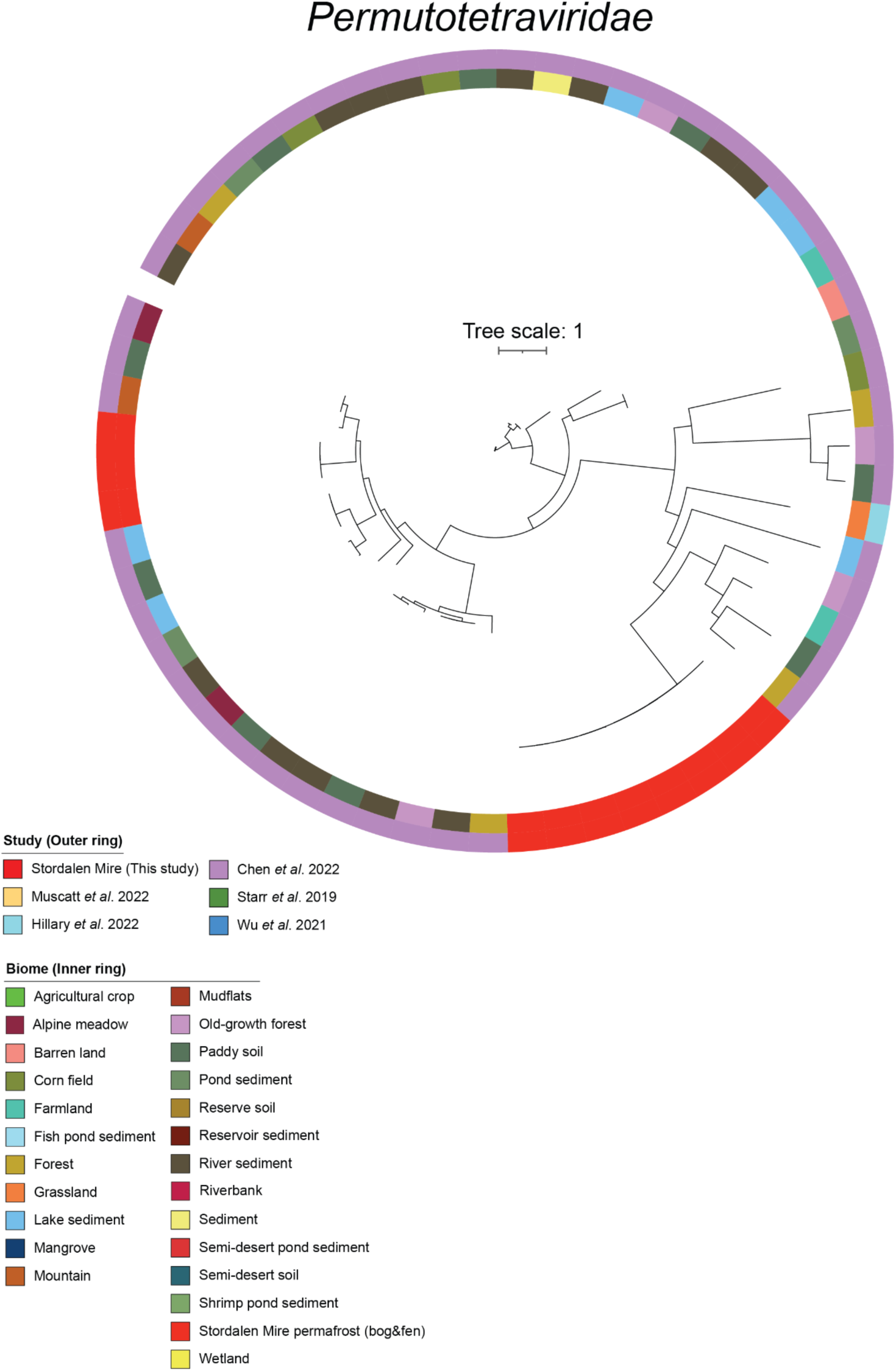
Thawing permafrost permutotetravirids. RdRp-based phylogenies across RNA virus studies. A maximum-likelihood phylogenetic tree was built from the RdRp-guided taxonomy analysis of near-complete RdRp domain sequences. The scale bar indicates one amino acid residue substitution per site. Sequences used to build the trees were preclustered at 40% identity, and clades supported by 100% bootstrap values were collapsed. The inner ring represents the biomes of these viruses whereas the outer ring represents soil RNA virus studies.

**Fig. S11.**
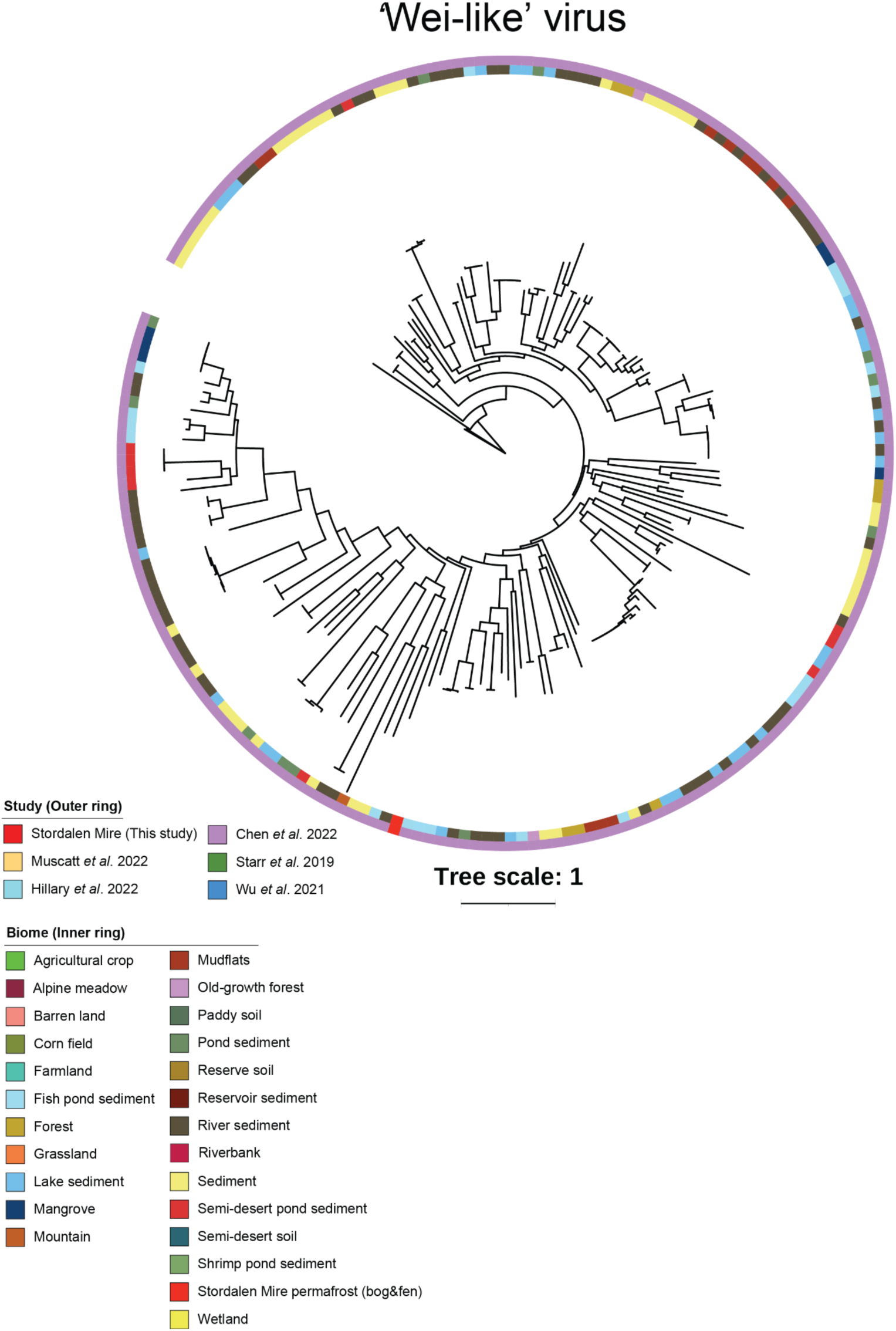
Thawing permafrost ‘wei-like’ viruses. RdRp-based phylogenies across RNA virus studies. A maximum-likelihood phylogenetic tree was built from the RdRp-guided taxonomy analysis of near-complete RdRp domain sequences. The scale bar indicates one amino acid residue substitution per site. Sequences used to build the trees were preclustered at 40% identity, and clades supported by 100% bootstrap values were collapsed. The inner ring represents the biomes of these viruses whereas the outer ring represents soil RNA virus studies.

**Fig. S12.**
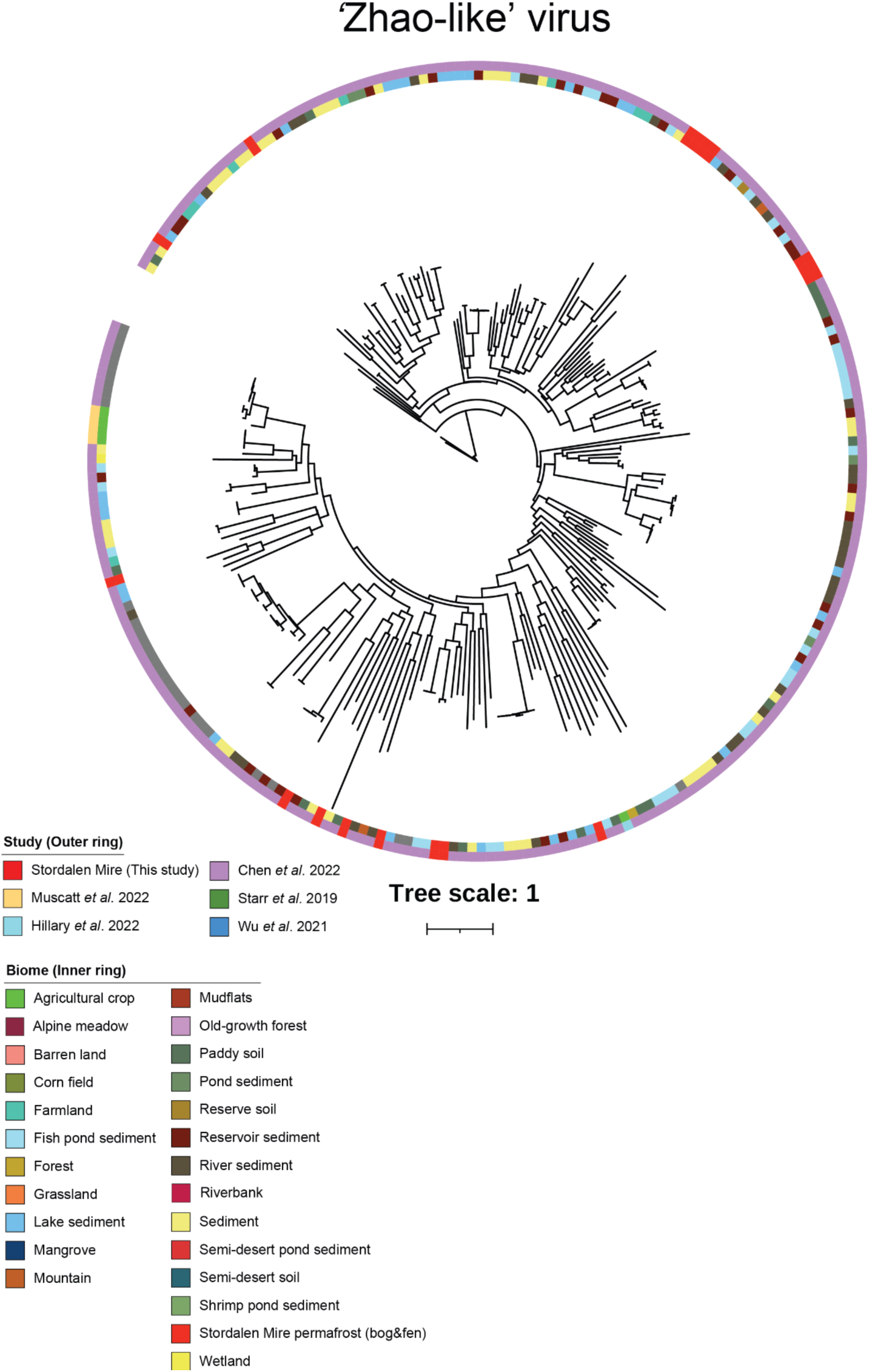
Thawing permafrost ‘Zhao-like’ viruses. RdRp-based phylogenies across RNA virus studies. A maximum-likelihood phylogenetic tree was built from the RdRp-guided taxonomy analysis of near-complete RdRp domain sequences. The scale bar indicates one amino acid residue substitution per site. Sequences used to build the trees were preclustered at 40% identity, and clades supported by 100% bootstrap values were collapsed. The inner ring represents the biomes of these viruses whereas the outer ring represents soil RNA virus studies.

**Fig. S13.**
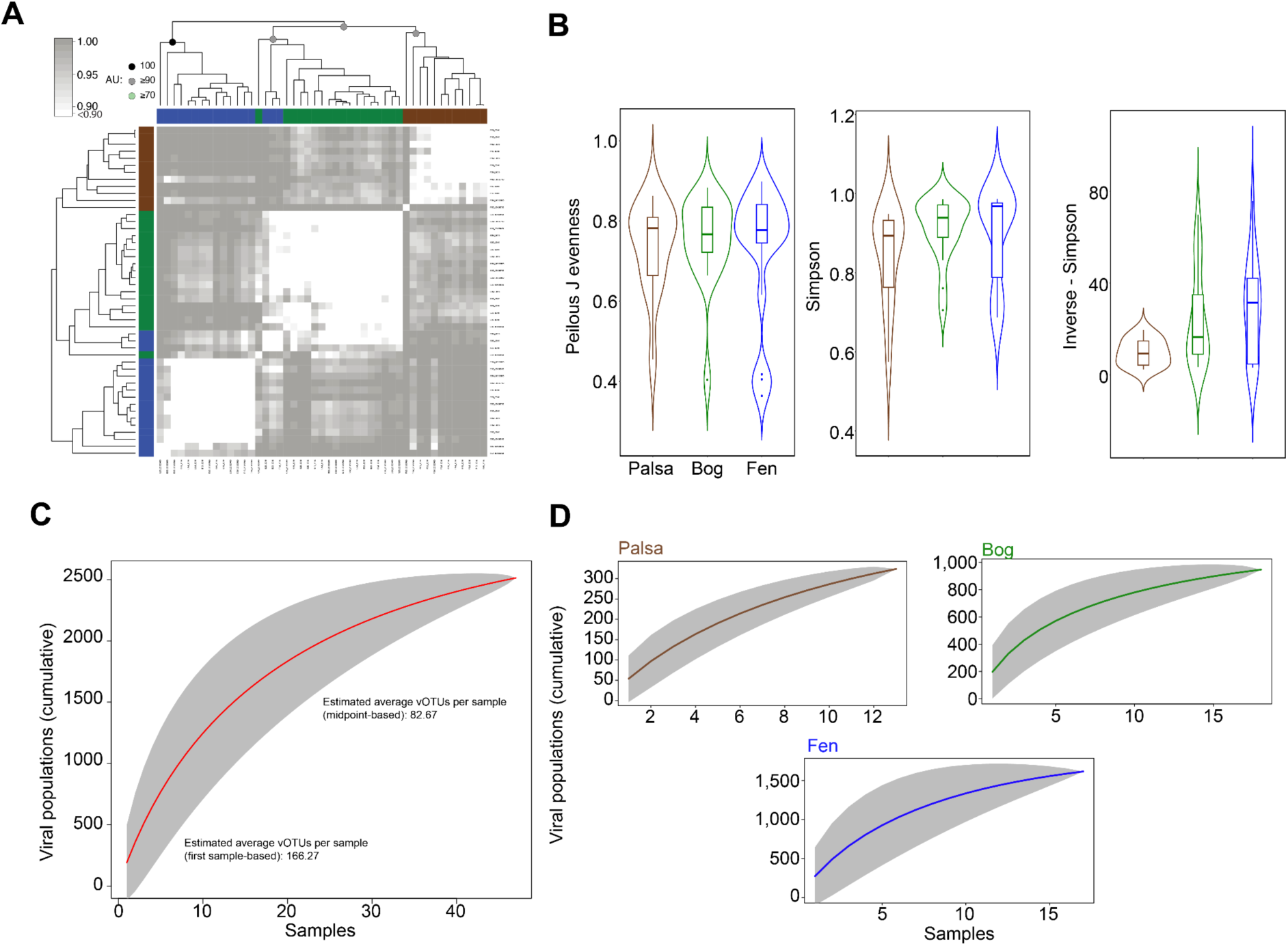
Stordalen Mire orthornaviraen ecology and diversity analysis. **(A)** Correlation-based hierarchical clustering of a Bray-Curtis dissimilarity matrix calculated from a randomly subsampled set from the vOTUs. The hierarchical clustering analysis structured orthornaviraens into three distinct meta-communities (brown; palsa, green; bog, and blue; fen) with an approximately unbiased (AU) bootstrap value ≥90. **(B)** Violin plots depict the diversity analysis of different metrics, including perilous J evenness, Simpson, and Inverse-Simpson. Diversity metrics were calculated using the “vegan” package in R. Statistical analysis was performed using Kruskal-Wallis analysis, with *post hoc* Dunn-test and *p*-adjusted: Bonferroni. Only the significant values are shown and donated as follows, *: *p*-value ≤0.05; **: *p*-value ≤ 0.01. **(C)** An accumulation curve of orthornaviraens in metatranscriptomes (*n* = 49, restricted to 2012 and 2016 data). Means are represented by red circles and 200 randomizations of sample order are shown in teal. Estimation of average vOTUs per sample from species accumulation curves. The first sample-based estimate (∼166 vOTUs) represents an upper bound, while the midpoint-based estimate (∼83 vOTUs) provides a more conservative average. and **(D)** Accumulative curve for palsa (*n* = 13), bog (*n* = 18), and fen (*n* = 18) (restricted to 2012 and 2016 data). Means are represented by brown, green, and blue circles (for palsa, bog, and fen, respectively), and the confidence interval (95%) of the sample order is shown in grey.

**Fig. S14.**
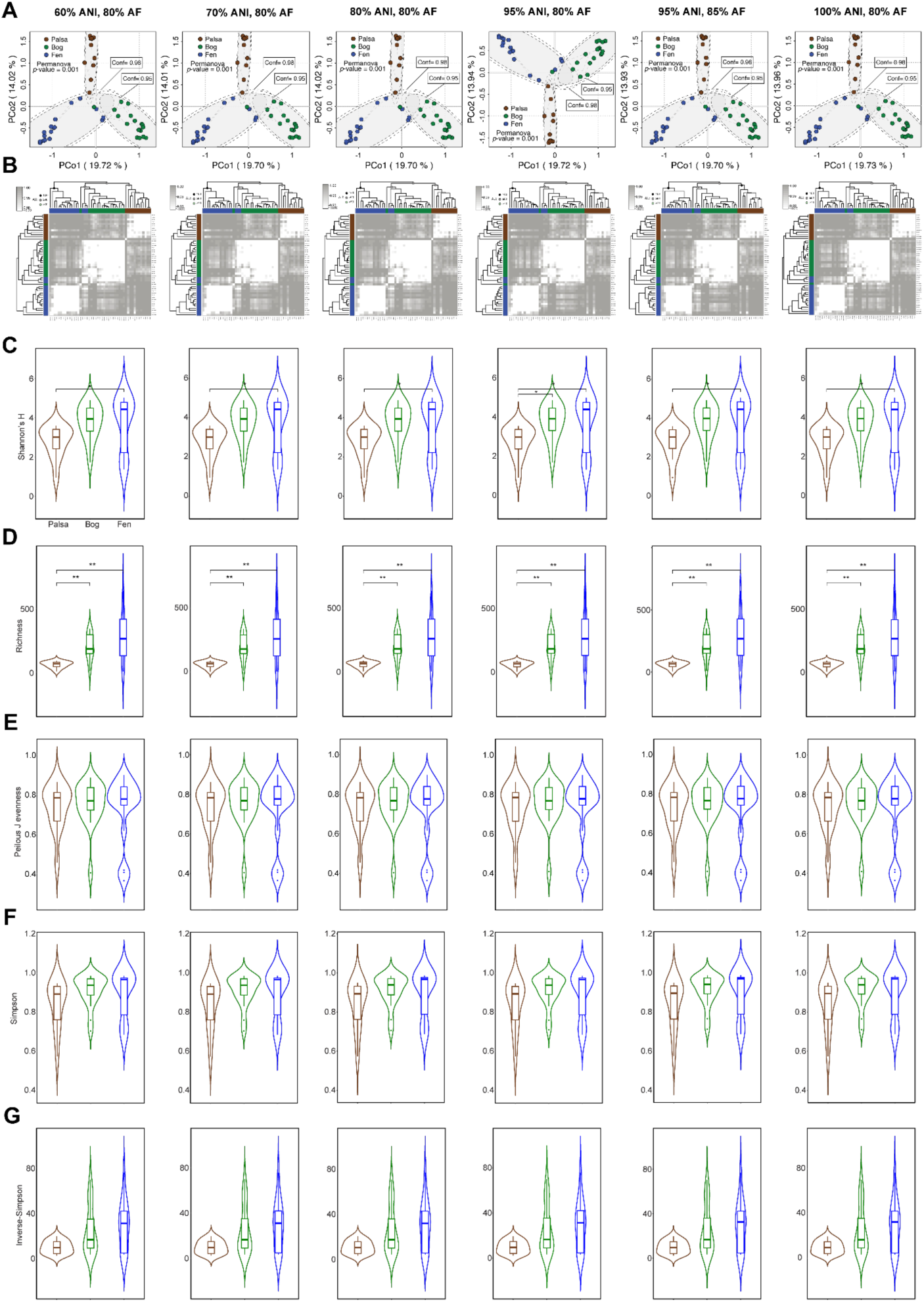
Sensitivity analyses for the robustness of ecological inferences under different vOTU definitions. (A) Principal component analysis (PCoA) of a Bray-Curtis dissimilarity matrix calculated from all vOTUs in this study. Dot colors correspond to Stordalen Mire habitats as in previous figures. Correlation-based hierarchical clustering of a Bray-Curtis dissimilarity matrix calculated from a randomly subsampled set from vOTUs. Hierarchical clustering analysis sorted orthornaviraens into three distinct meta-communities (brown, palsa; green, bog; and blue, fen) with an approximately unbiased (AU) bootstrap value ≥90. (**C, D, E, F, G**) Violin plots (with boxplots) depict the diversity analysis of different metrics, including Shannon’s *H*, richness, peilous J evenness, and Simpson and Inverse-Simpson. Statistical analysis was performed using Kruskal-Wallis analysis, with *post hoc* Dunn-test and *p*-adjusted: Bonferroni. Only significant values are shown and donated as follows, *: *p*-value ≤0.05; **: *p*-value ≤0.01.

**Fig. S15.**
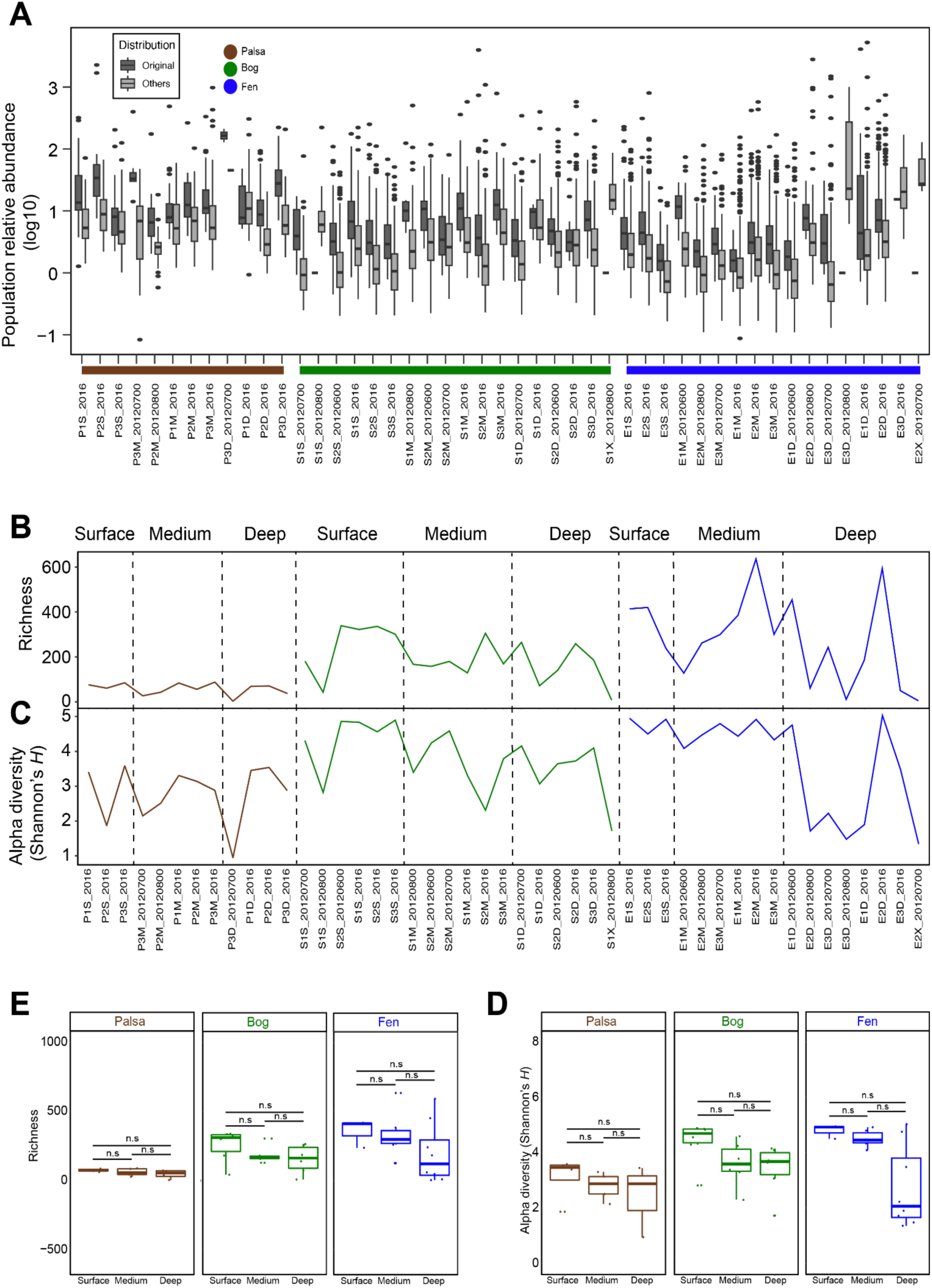
Ecological patterns of Stordalen Mire orthornaviraens across habitats. (A) Relative abundance of vOTUs in original samples, from which vOTUs were assembled and compared in respect to abundance compared to other samples. (**B, C, D**, and **E**) Depth profiles of richness and alpha diversities (Shannon’s *H*) of each sample, respectively. Samples were ordered based on depth, thaw gradient progress (palsa, bog, and fen) and sampling order. Statistical significance was evaluated with ANOVA (ns: not significant).

**Fig. S16.**
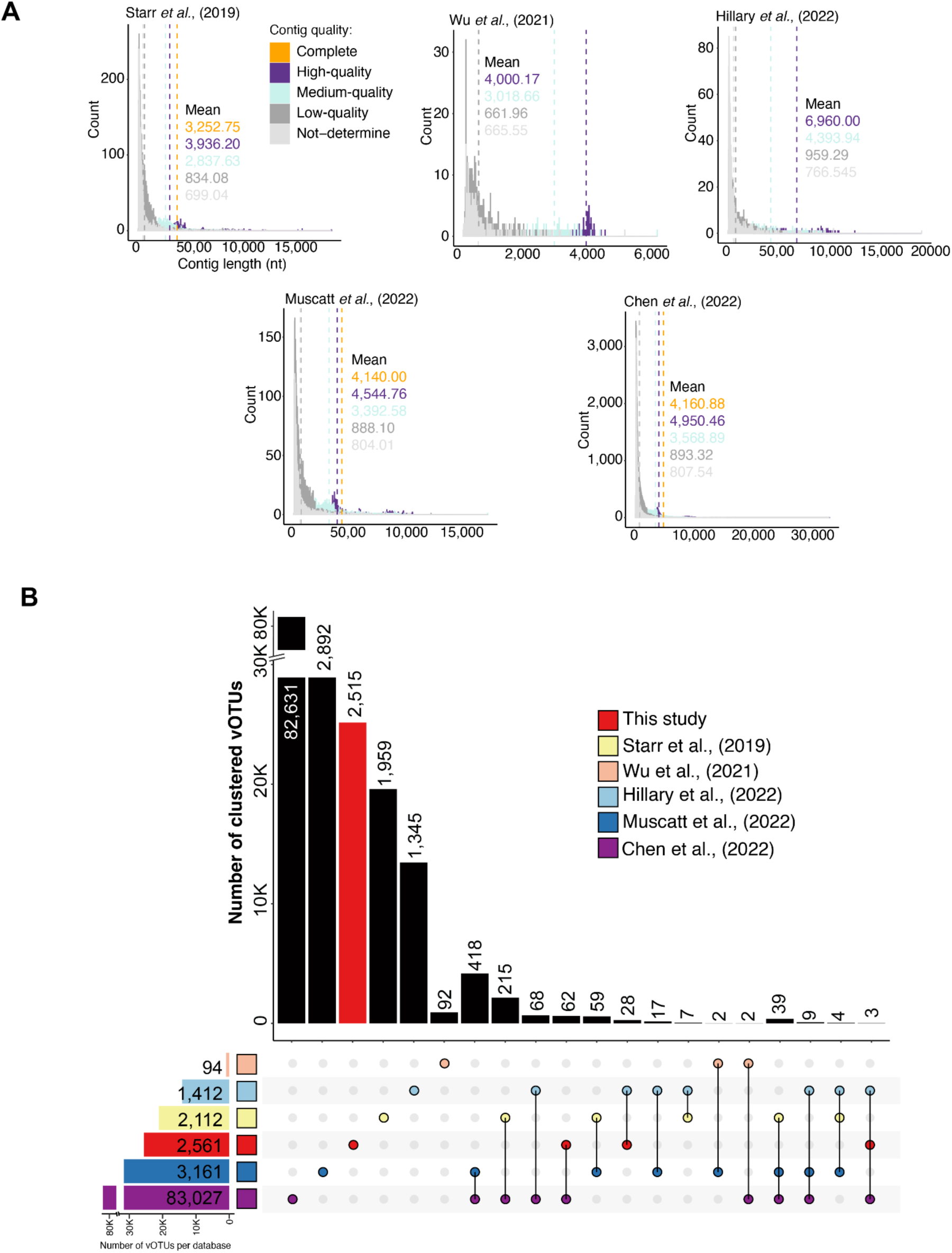
Comparison of orthornaviraen counts and RdRps across soil studies. (A) The color dotted lines represent the length means of virus genomes based on contig quality. (B) UpSet plot depicts the number of shared and unique vOTUs (90% ID and 80% coverage).

**Fig. S17.**
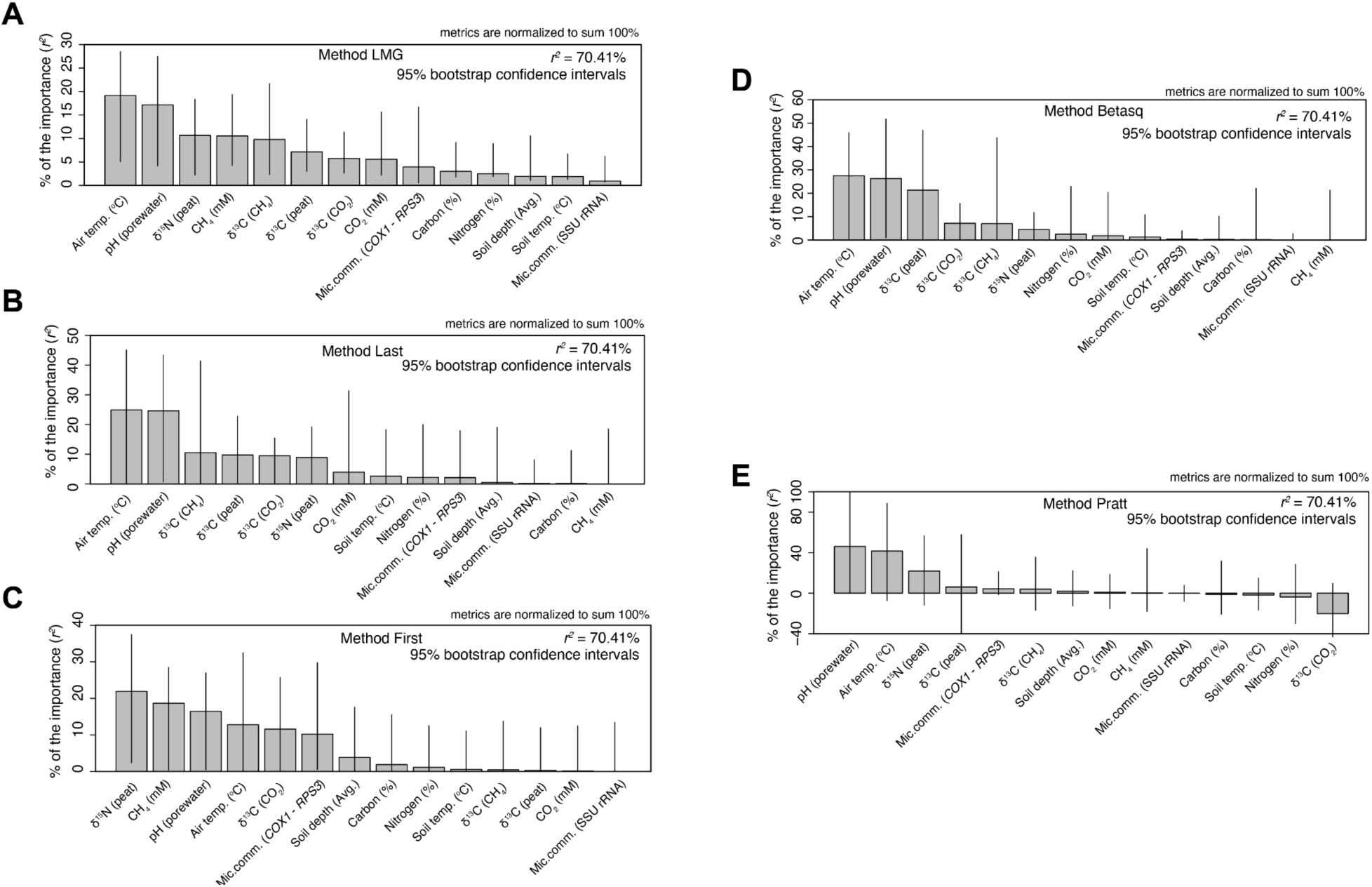
pH is a significant predictor of orthornaviraens in the Stordalen Mire thawing permafrost. Relative importance of environmental variables as predictors of estimated virus community (the first coordinate of a PCoA – Pco1). Analyses were performed using R package “relaimpo” v2.2-3 (*84*). This package enables univariate linear regression analyses using five different methods (**A**) LMG, (**B**) Last (**C**) First, (**D**) Betasq, and (**E**) Pratt, and 1,000 bootstraps for confidence estimates. *R*^2^ denotes the proportion of response variance explained by the model. Bars show the mean proportion of response variance (i.e., relative contribution) of each variable based on 1,000 bootstrap replicates. Error bars indicate the 95% bootstrap confidence intervals of the response variance (in percentages). Only samples from 2011 and 2012 were used in the analysis, as no metadata were available for 2016.

**Fig. S18.**
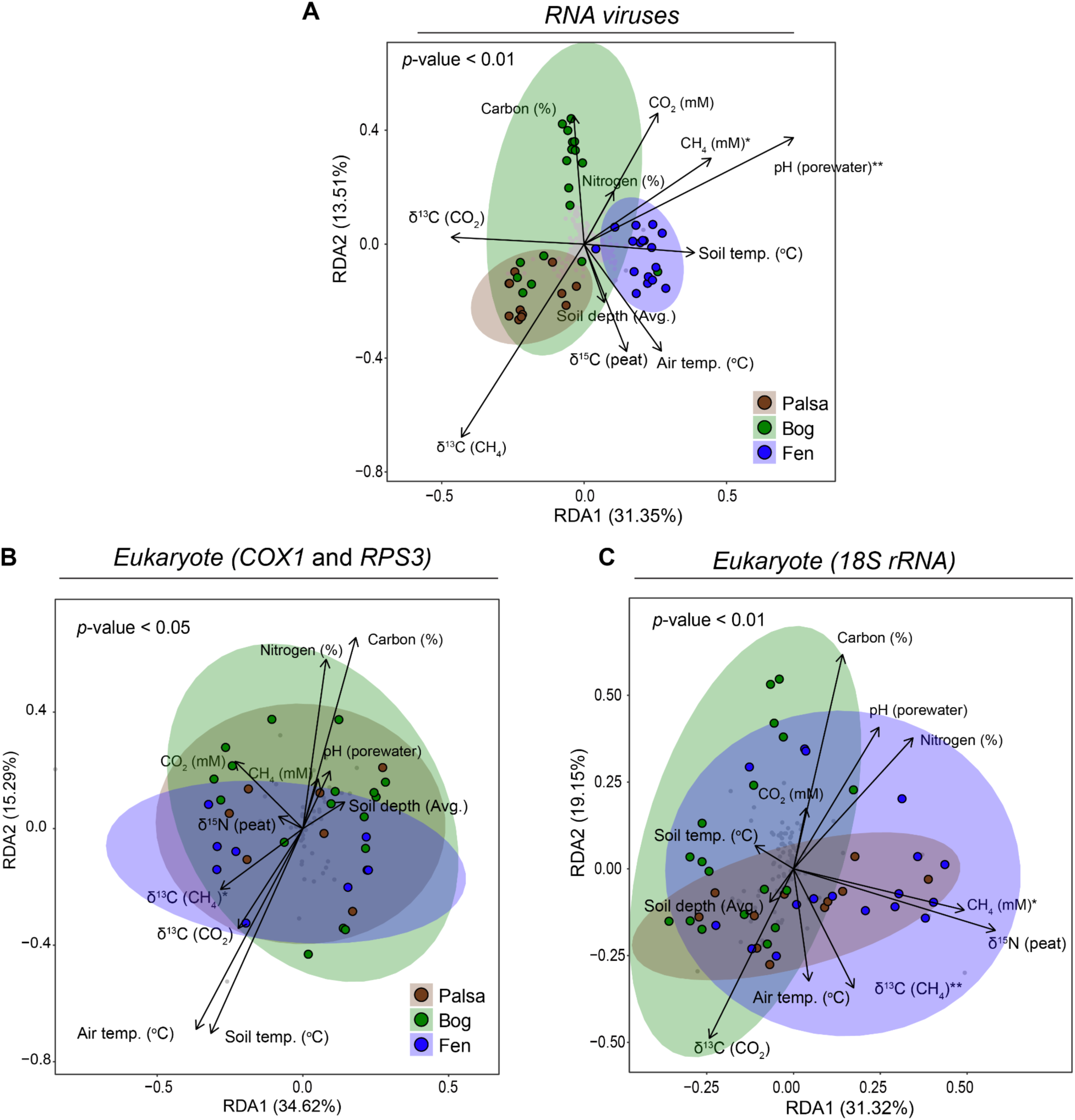
Redundancy Analysis (RDA) of RNA viruses and Stordalen Mire eukaryotes and prokaryotes. Redundancy analysis (RDA) of (**A**) RNA viruses, (**B**) Eukaryotes (based on *COX1* and *RPS3*) and (**C**) prokaryotes (based on 18S rRNA) of the relationship between the relative abundance of organisms across habitats and abiotic factors. Significance codes:; **: 0.01; *: 0.05.

**Fig. S19.**
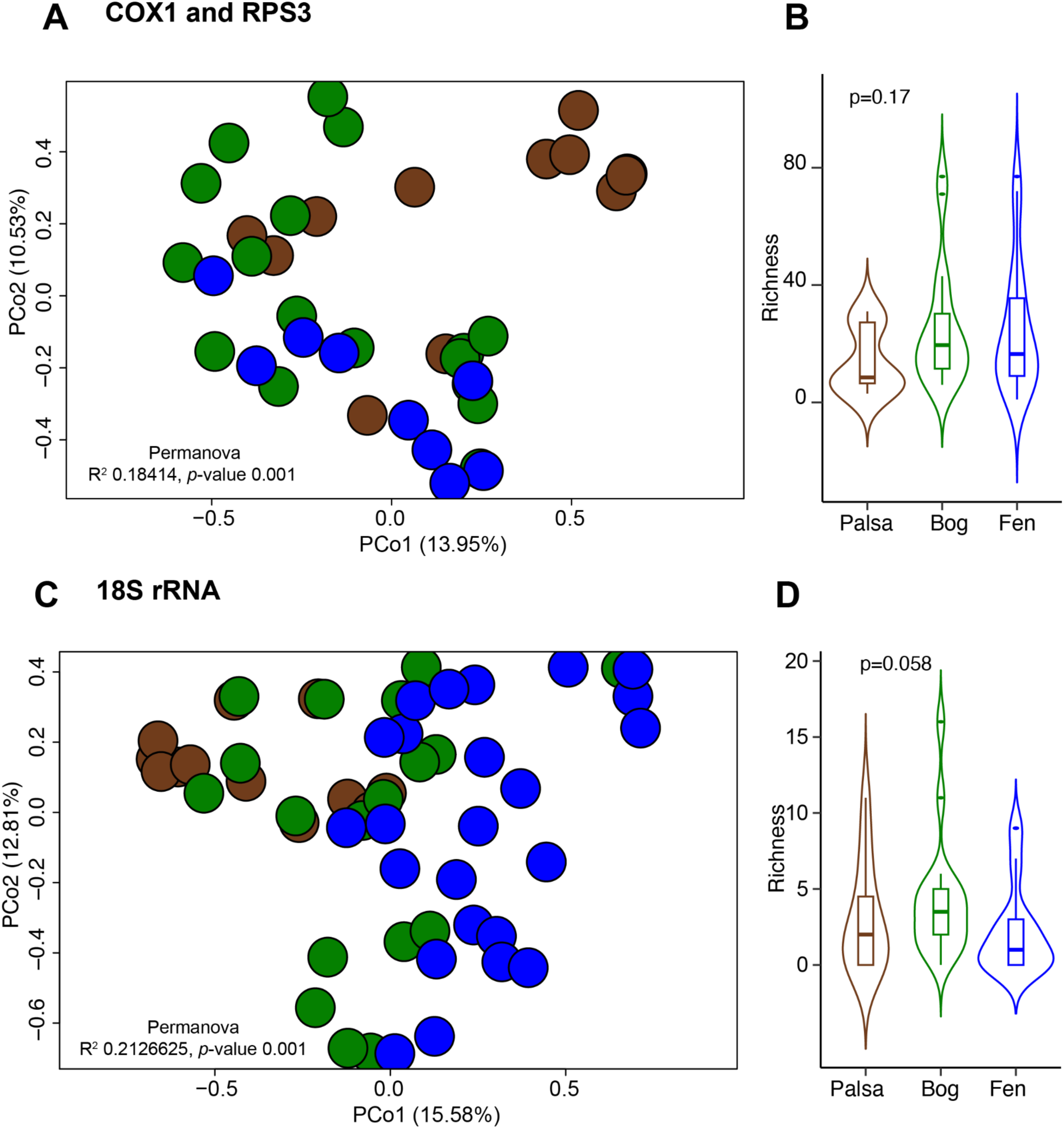
Organismal ecology. (**A** and **C**) Principal component analysis (PCoA) of a Bray-Curtis dissimilarity matrix calculated from all organism OTUs in this study deduced via marker gene and 18S rRNA sequencing, respectively. Violin plots (with boxplots) depict the Richness for (**B**) *COX1* and *RPS3* genes and (**D**) 18S rRNA, respectively. Statistical analysis was performed using Kruskal-Wallis analysis, with *post hoc* Dunn-test and *p*-adjusted/and ANOVA (for those with normal distribution data): Bonferroni.

**Fig. S20.**
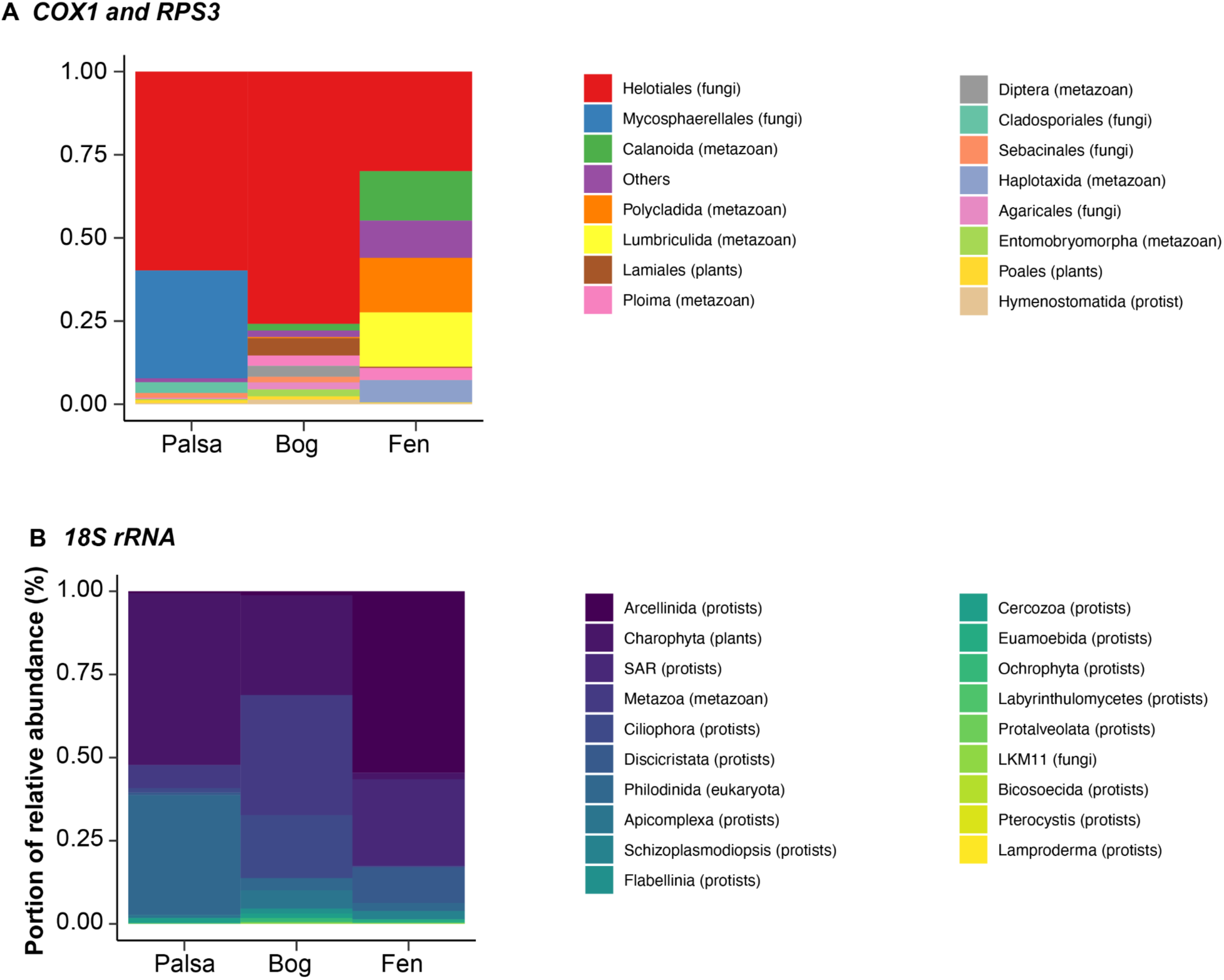
Microbial organism taxonomy across thawing gradients. The stacked-bar plots depict the proportion of relative abundance (%) based on metagenomics mapping. Taxonomy assignment at the order-level of organisms in Stordalen Mire identified by **(A)** *COX1* and *RPS3* genes, and (**B**) organisms identified by 18S rRNA gene.

**Fig. S21.**
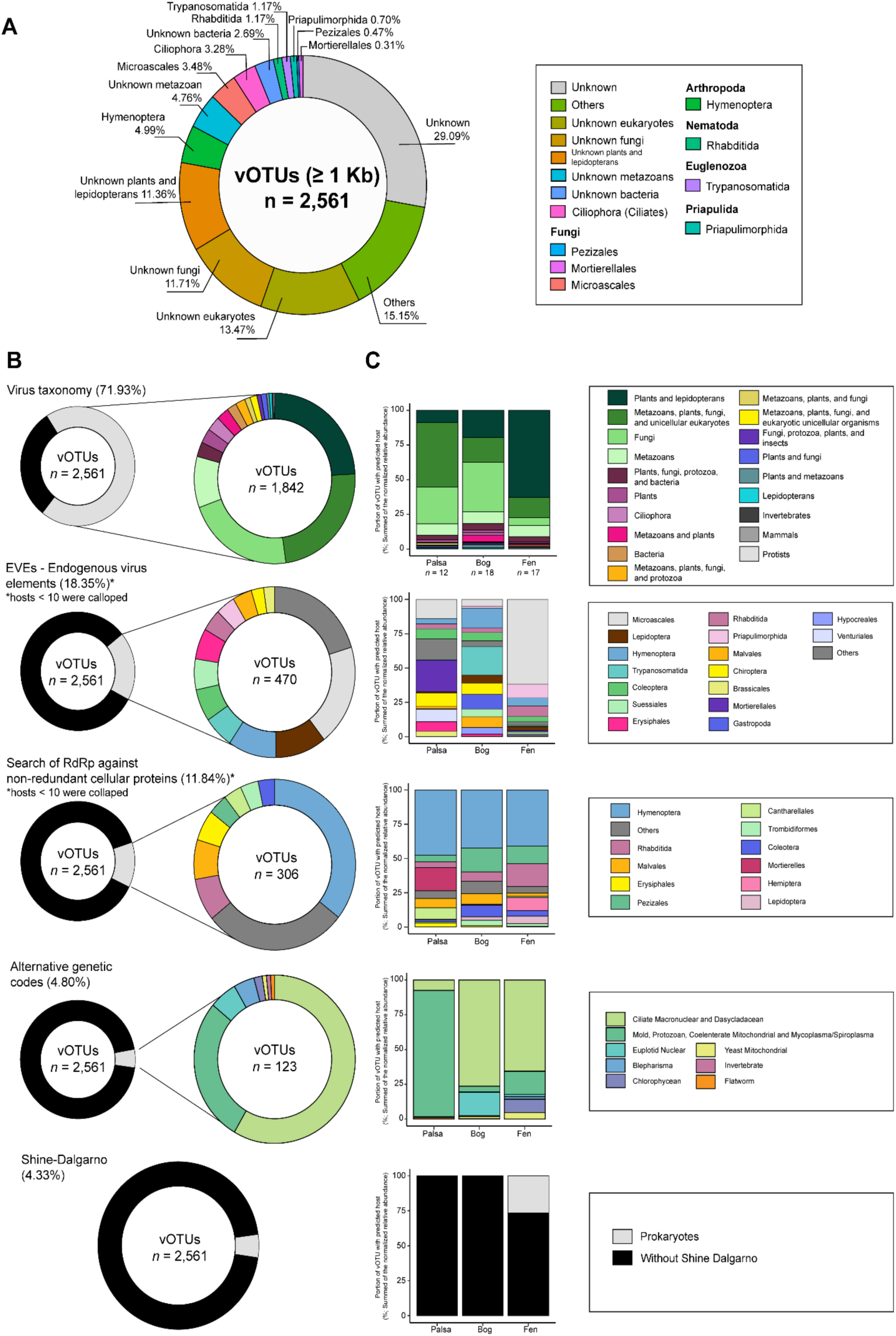
Inferred hosts of Stordalen Mire orthornaviraens. (**A**) Percent of inferred host based on the integrated approach. See analysis of inferred host using individual approach in panel B. (**B**) Host inferred analysis, including virus RdRp-based taxonomy, endogenous virus elements (EVEs), virus RdRps against non-redundant cellular proteins, alternative genetic codes and Shine-Dalgarno sequences. (**C**) Stacked bar plots depicting the percentage of vOTUs with the taxonomic affiliation of the inferred hosts per habitat. Only the top 10 taxonomy assignments are shown.

**Fig. S22.**
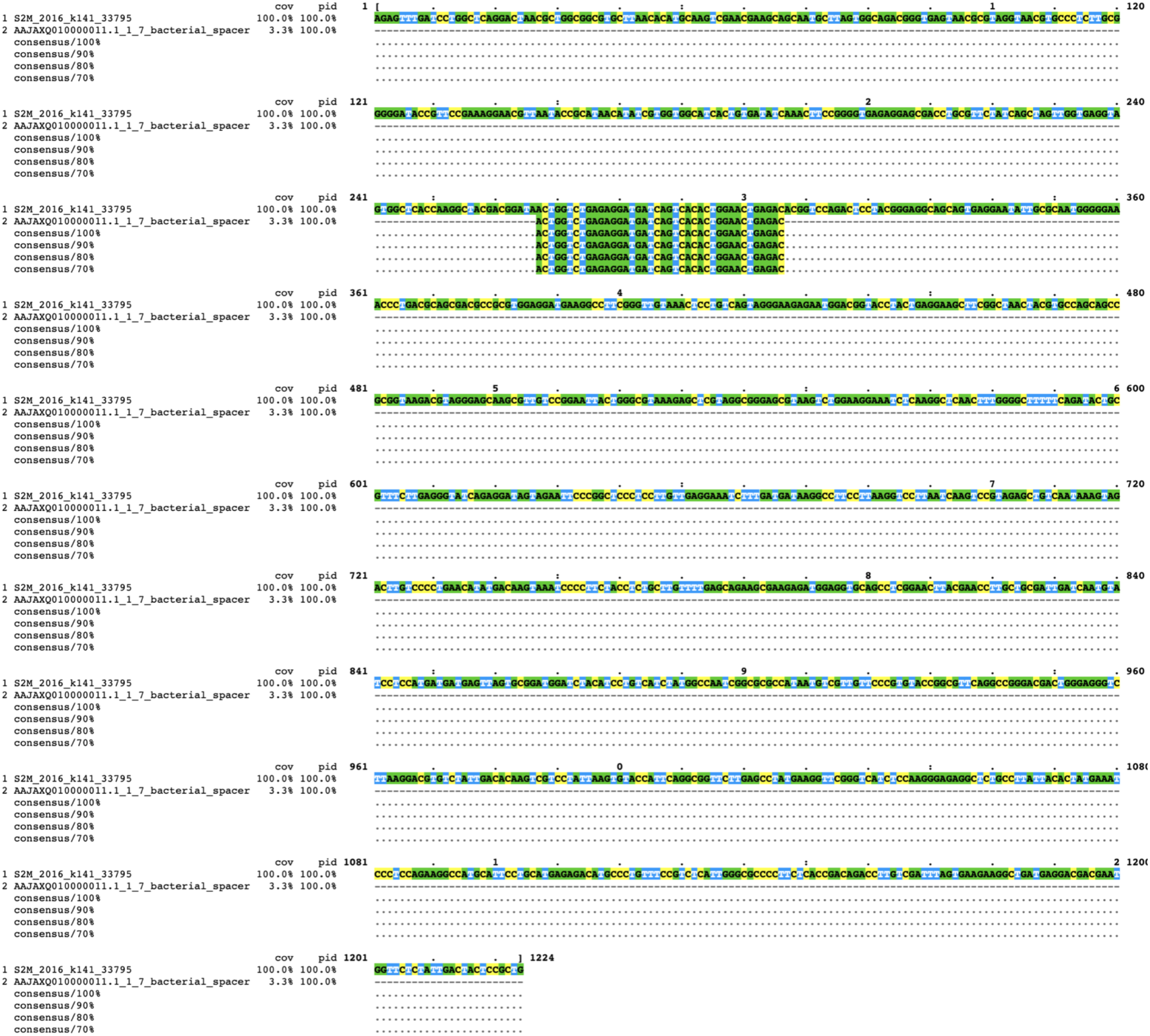
Search of CRISPR-Cas spacers across orthornaviran genomes. Global alignment of nucleotide sequences of an orthornaviraen contig (S2M_2016_k141_33795) and a CRISPR spacer of *Campylobacter lari* (AAJAXQ010000011.1) visualized using MView.

**Fig. S23.**
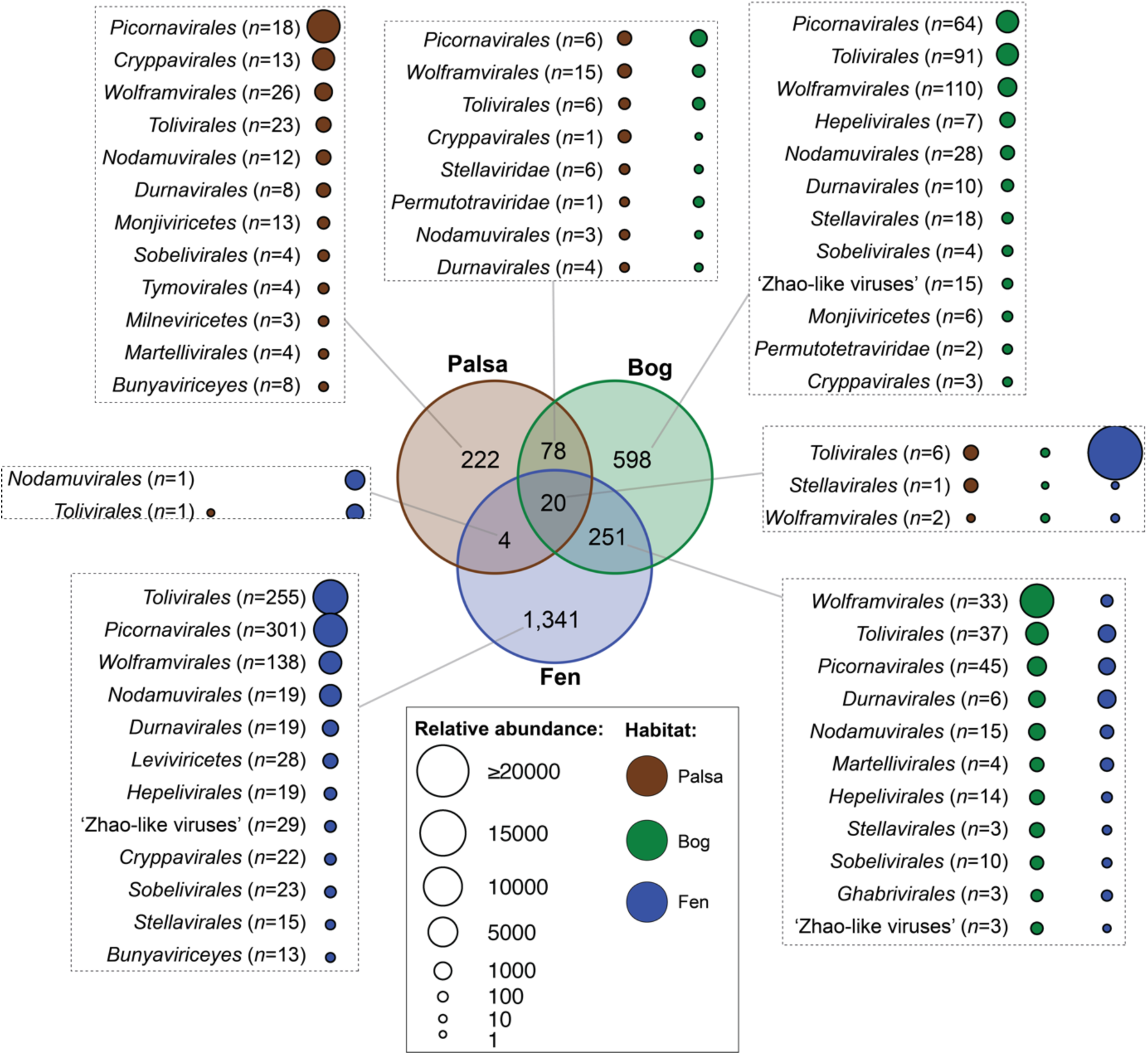
Venn diagram of Stordalen Mire orthornaviraens across habitats. A modified RdRp-scan (*80*) approach was used to assign order-ranked virus taxonomy to RdRp footprint sequences longer than 200 amino acid residues encoded by vOTUs. Dot sizes represent the summed relative abundances of vOTUs detected across habitats.

## Supplementary Tables

**S1 Table. Accession numbers Stordalen Mire metatranscriptomes used in this study.**

**S2 Table. Metadata: sampling IDs, locations, depth, biochemical measurements.**

**S3 Table. List of final RNA virus footprints and contigs.**

**S4 Table. Normalized relative abundances of vOTU.**

**S5 Table. MCL Network-based taxonomy assignment**

**S6 Table. Metagenomics assembed genomes were used in this analysis. Acquired from Woodcroft et al. 2018. Nature 560:49–54; 10.1038/s41586-018-0338-1**

**S7 Table. Integrated host interference analysis**

**S8 Table. Final auxiliary metabolic genes in RNA viruses**

