## Supplementary Material for "RNA virus ecogenomics along a subarctic permafrost thaw gradient"

### SUPPORTING INFORMATION

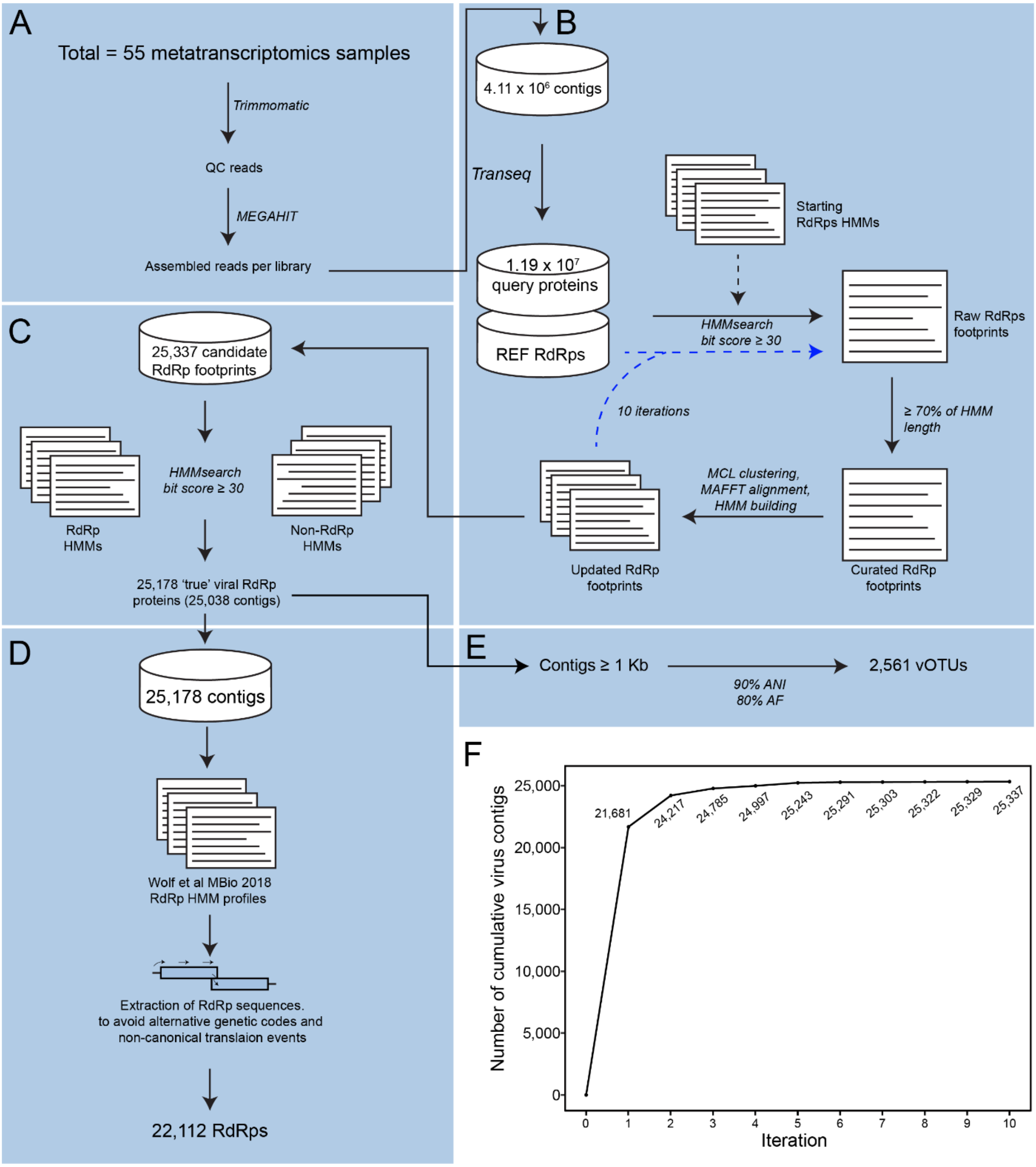


**Fig. S1. Bioinformatics workflow and Stordalen Mire orthornaviraen identification.** (**A**) The metatranscriptomes from Stordalen Mire permafrost, collected in 2010, 2011, 2012, and 2016, were preprocessed, including quality-trimming and assembly into contigs. (**B**) The deduced encoded protein sequences were then used to identify orthornaviraens. To capture highly divergent RdRps, a search-and-update Hidden Markov Model (HMMs) approach was used over ten iterations (dotted blue lines). (**C**) Competitive HMM profiles were used to evaluate the authenticity of RdRp hits. (**D**) Extraction and reconstruction of RdRp domain sequences were performed by running another iteration of searching and deducing protein sequences against the RdRp profile HMMs. (**E**) For ecological analysis, virus operational taxonomic units (vOTUs), i.e., approximate species-rank ecological units, at the 90% Average Nucleotide Acid (ANI), and an 80% alignment fraction (AF) were applied. (**F**) Virus contigs were detected as a result of 10 iterations of HMMsearch.

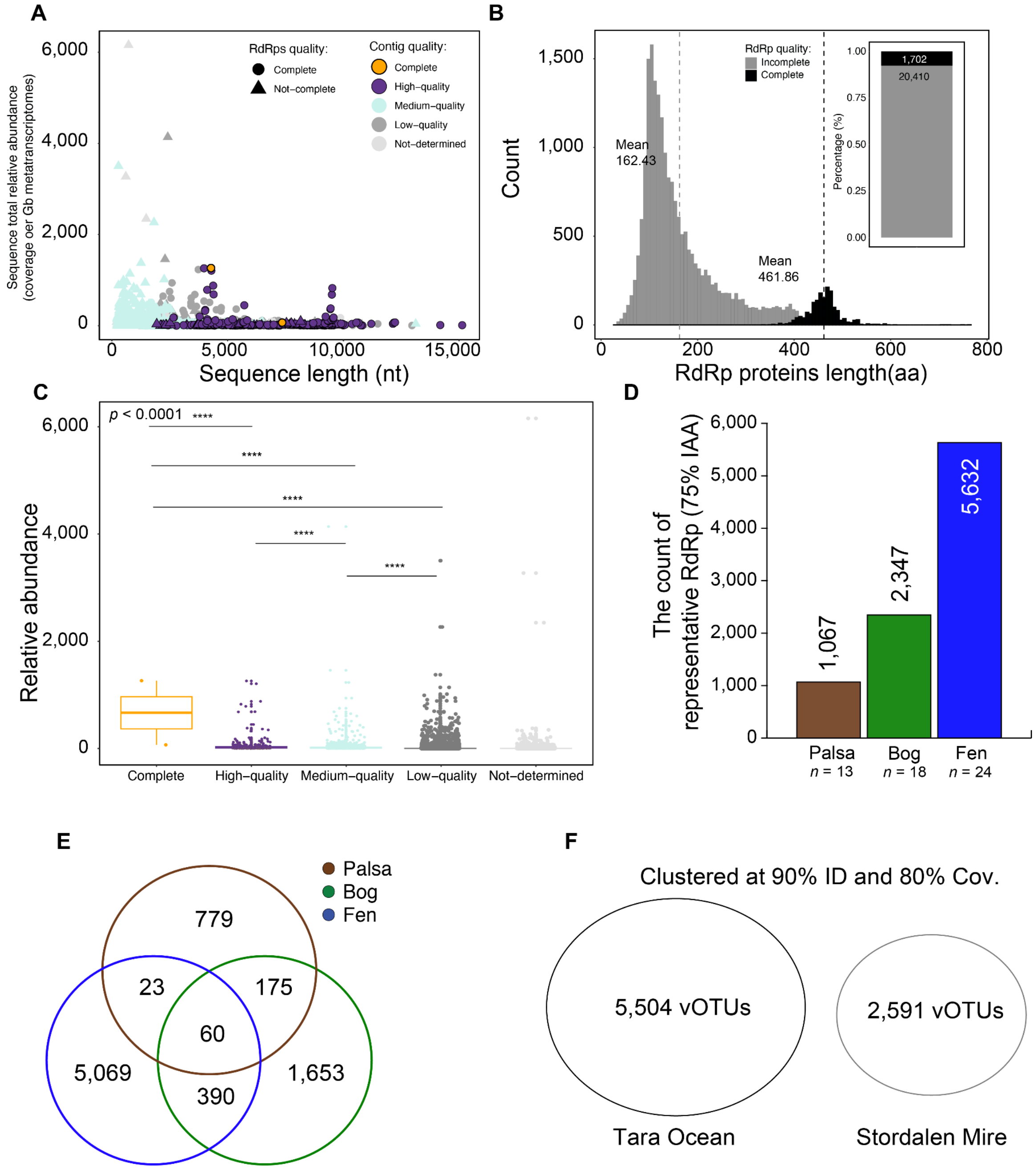


**Fig. S2. Completeness of Stordalen Mire orthornaviran genomes and RdRp domains.** (**A**) Lengths of 22,112 analyzed contigs encoding virus RdRps (*x* axis) and their cumulative coverage across Stordalen Mire (*y* axis), indicating completeness of virus genome sequences and RdRp domains. Contig quality analysis was performed using CheckV. (**B**) Histogram depicting the length distribution of complete and incomplete RdRp domain sequences. The inset stacked barplot indicates the percentage of complete RdRp domains. (**C**) Stordalen Mire orthornaviran contig quality (based on CheckV). Statistical analysis was performed using Kruskal–Wallis analysis, with *post hoc* Dunn-test and *p*-adjusted: Bonferroni. Only *p*-values ≤ 0.0001 are shown (****). (**D**) Counts of RdRp representative footprints per habitat at the 75% amino acid identity (AAI) level. (**E**) Counts of RdRp footprint representatives across habitats at 75% AAI (number of samples as indicated in D). (**F**) Clustering of 5,504 *Tara* ocean RNA viruses and 2,591 vOTUs of RNA viruses from Stordalen Mire at 90% identity and 80% coverage. No overlapping of viruses at the "species-level".

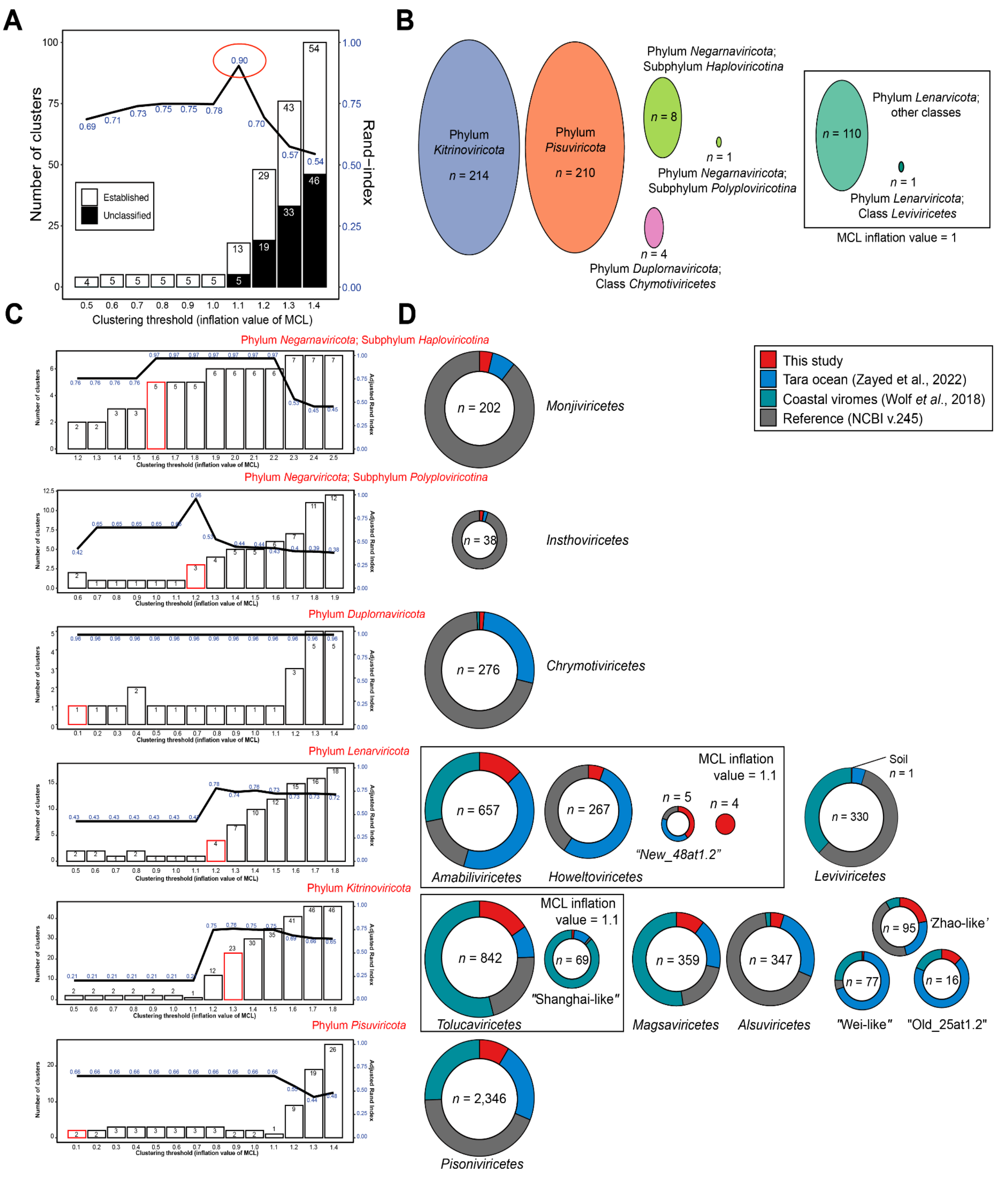


**Fig. S3. Establishment of RdRp domain-based phylum- and class-rank clusters.** (**A**) adjusted rand index (line) of the network-guided and phylogeny-based megataxonomy at different clustering thresholds. Stacked bars represent the number of taxonomic clusters of near-complete RdRp domains (at least 90% of the domain) at the depicted clustering thresholds. Only sequences representing established taxa (black) were used for calculating the agreement percentage. The highest agreement is shown at an inflation value of 1.1, forming a total of 18 clustered phyla; five were previously established by Wolf et al. (**B**) Established taxa at the Markov Clustering Algorithm (MCL) inflation value of 1.1, represented by ellipses in distinct colors. The box depicts taxa that were exclusively joined at lower inflation values. (**C**) adjusted rand index (line) of the network-guided and phylogeny-based at phylum/subphylum rank. (**D**) Pie charts represent the proportion of orthornaviraens of the "complete"/"near-complete" RdRp domains (of the network-guided phylogeny analysis) at class rank clusters, including sequences from this study (red), and references (NCBI—grey, *Tara* Ocean—blue, and coastal ocean viromes—green). The box depicts taxa that were exclusively joined at lower inflation values.

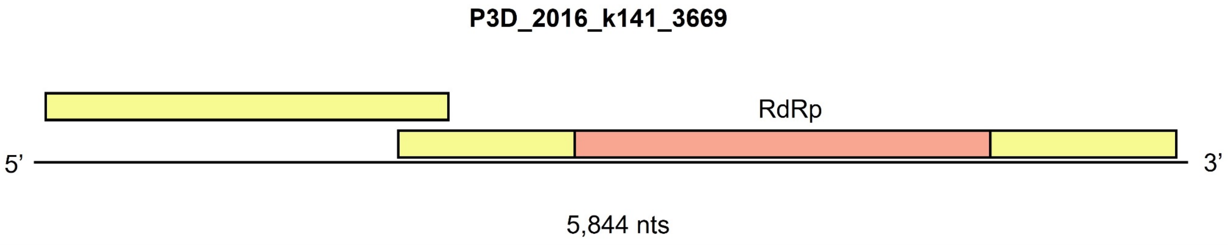


**Fig. S4. Genome of novel class ‘*Stormiviricetes*’.**
The longest genome of the newly identified class of ‘*Stormiviricetes*”. Yellow boxes represent ORFs in 5’→3’ sense and coral boxes represent the RdRp domain.

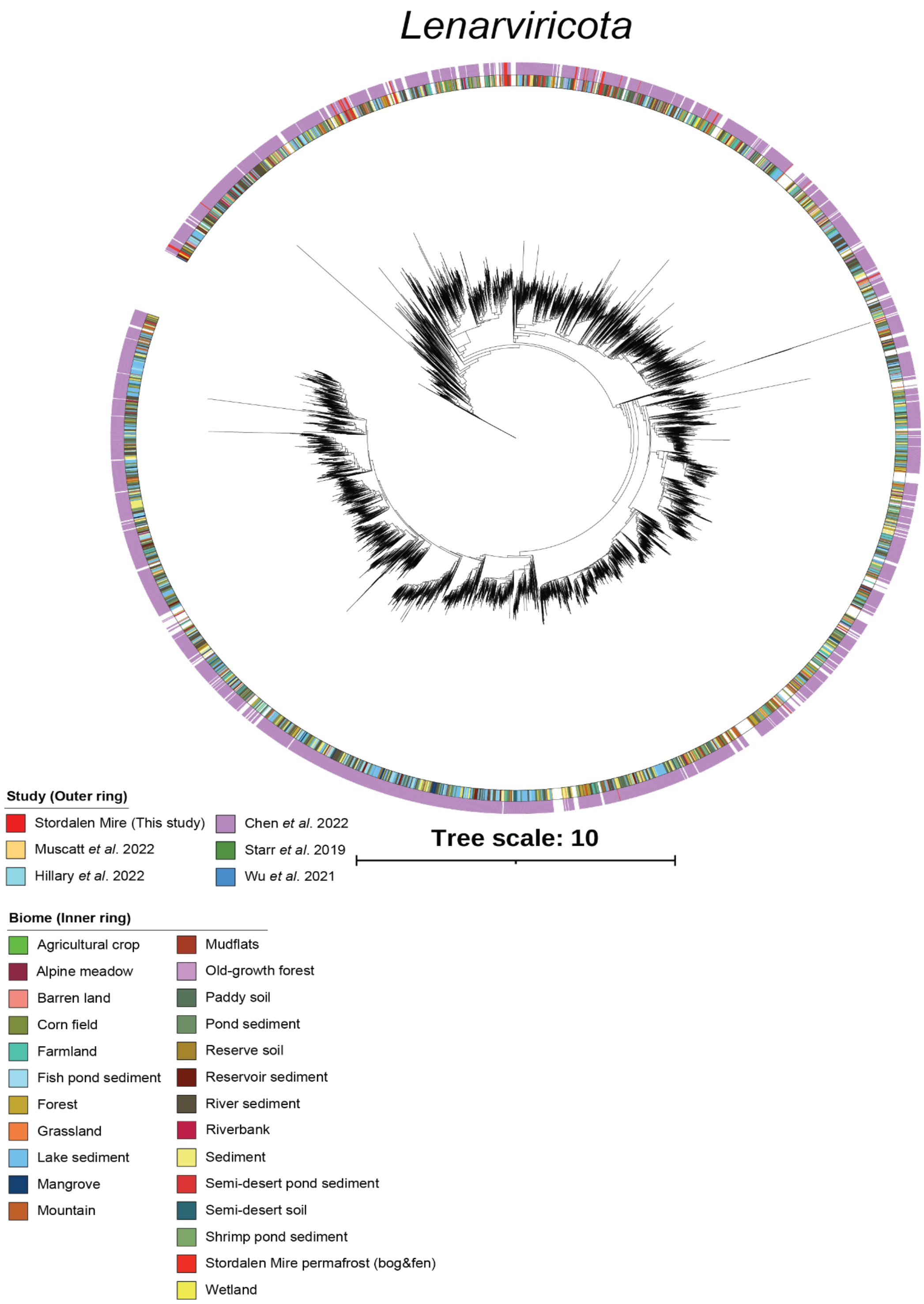


**Fig. S5. Thawing permafrost lenarviricots. RdRp-based phylogenies across RNA virus studies.**

A maximum-likelihood phylogenetic tree was built from the RdRp-guided taxonomy analysis of near-complete RdRp domain sequences. The scale bar indicates one amino acid residue substitution per site. Sequences used to build the trees were preclustered at 40% identity, and clades supported by 100% bootstrap values were collapsed. The inner ring represents the biomes of these viruses whereas the outer ring represents soil RNA virus studies.

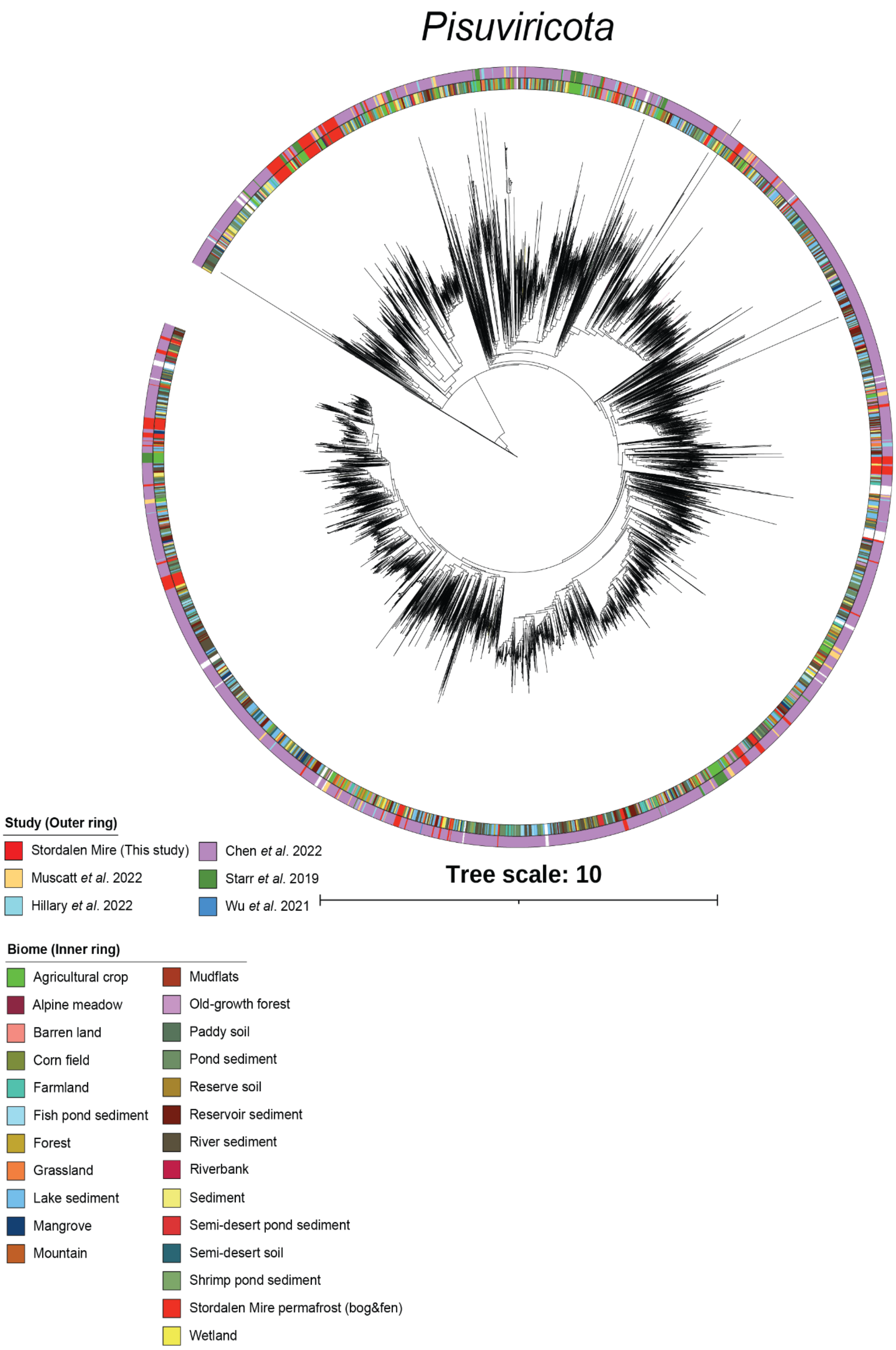


**Fig. S6. Thawing permafrost pisuviricots. RdRp-based phylogenies across RNA virus studies.**
A maximum-likelihood phylogenetic tree was built from the RdRp-guided taxonomy analysis of near-complete RdRp domain sequences. The scale bar indicates one amino acid residue substitution per site. Sequences used to build the trees were preclustered at 40% identity, and clades supported by 100% bootstrap values were collapsed. The inner ring represents the biomes of these viruses whereas the outer ring represents soil RNA virus studies.

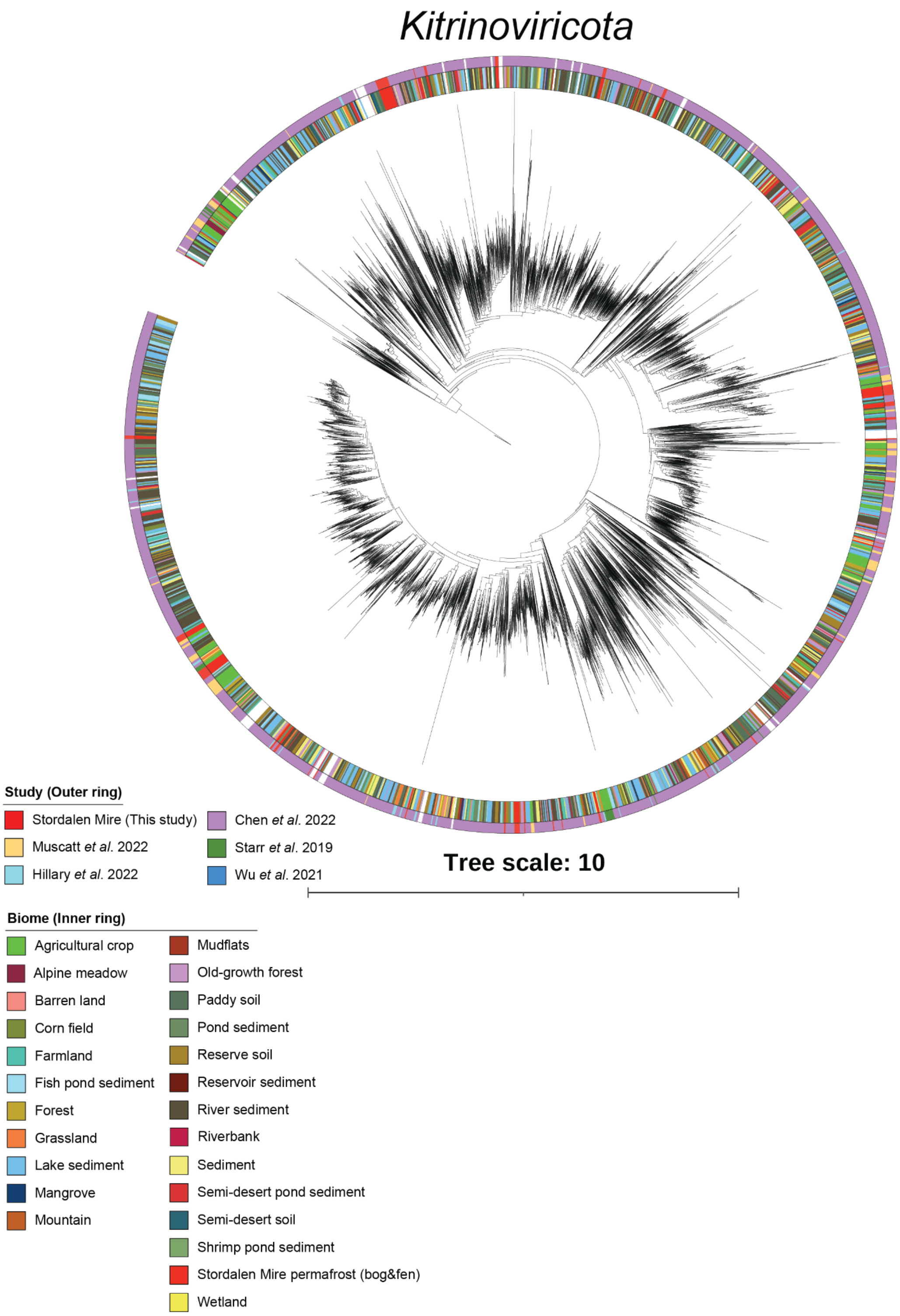


**Fig. S7. Thawing permafrost kitrinoviricots. RdRp-based phylogenies across RNA virus studies.**

A maximum-likelihood phylogenetic tree was built from the RdRp-guided taxonomy analysis of near-complete RdRp domain sequences. The scale bar indicates one amino acid residue substitution per site. Sequences used to build the trees were preclustered at 40% identity, and clades supported by 100% bootstrap values were collapsed. The inner ring represents the biomes of these viruses whereas the outer ring represents soil RNA virus studies.

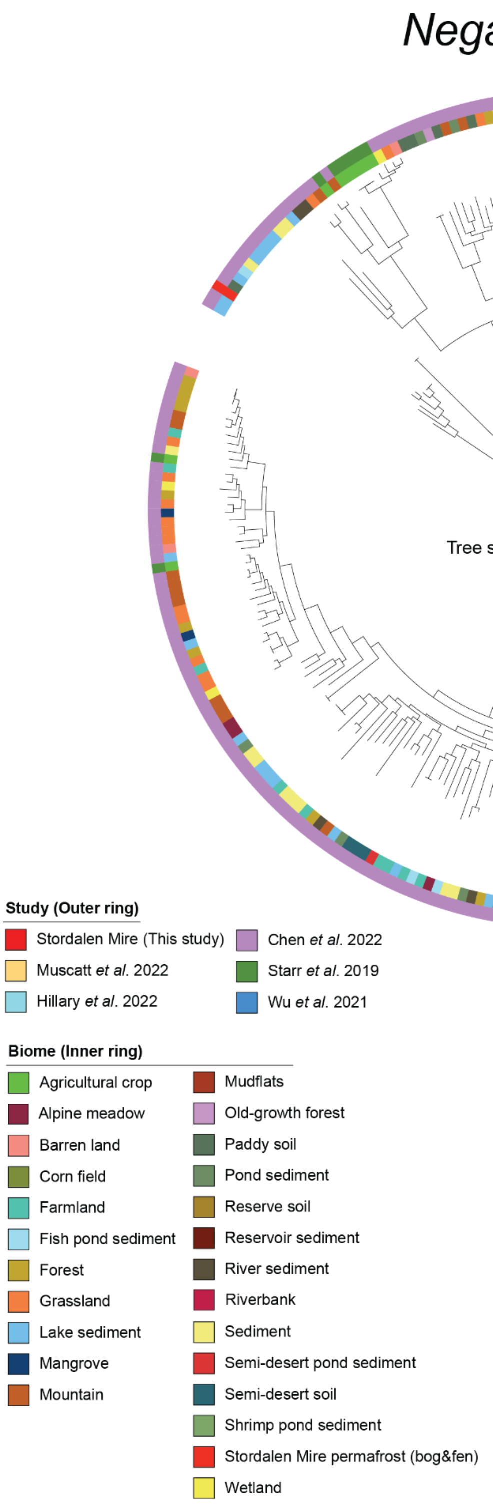


**Fig. S8. Thawing permafrost negarnaviricots. RdRp-based phylogenies across RNA virus studies.**

A maximum-likelihood phylogenetic tree was built from the RdRp-guided taxonomy analysis of near-complete RdRp domain sequences. The scale bar indicates one amino acid residue substitution per site. Sequences used to build the trees were preclustered at 40% identity, and clades supported by 100% bootstrap values were collapsed. The inner ring represents the biomes of these viruses whereas the outer ring represents soil RNA virus studies.

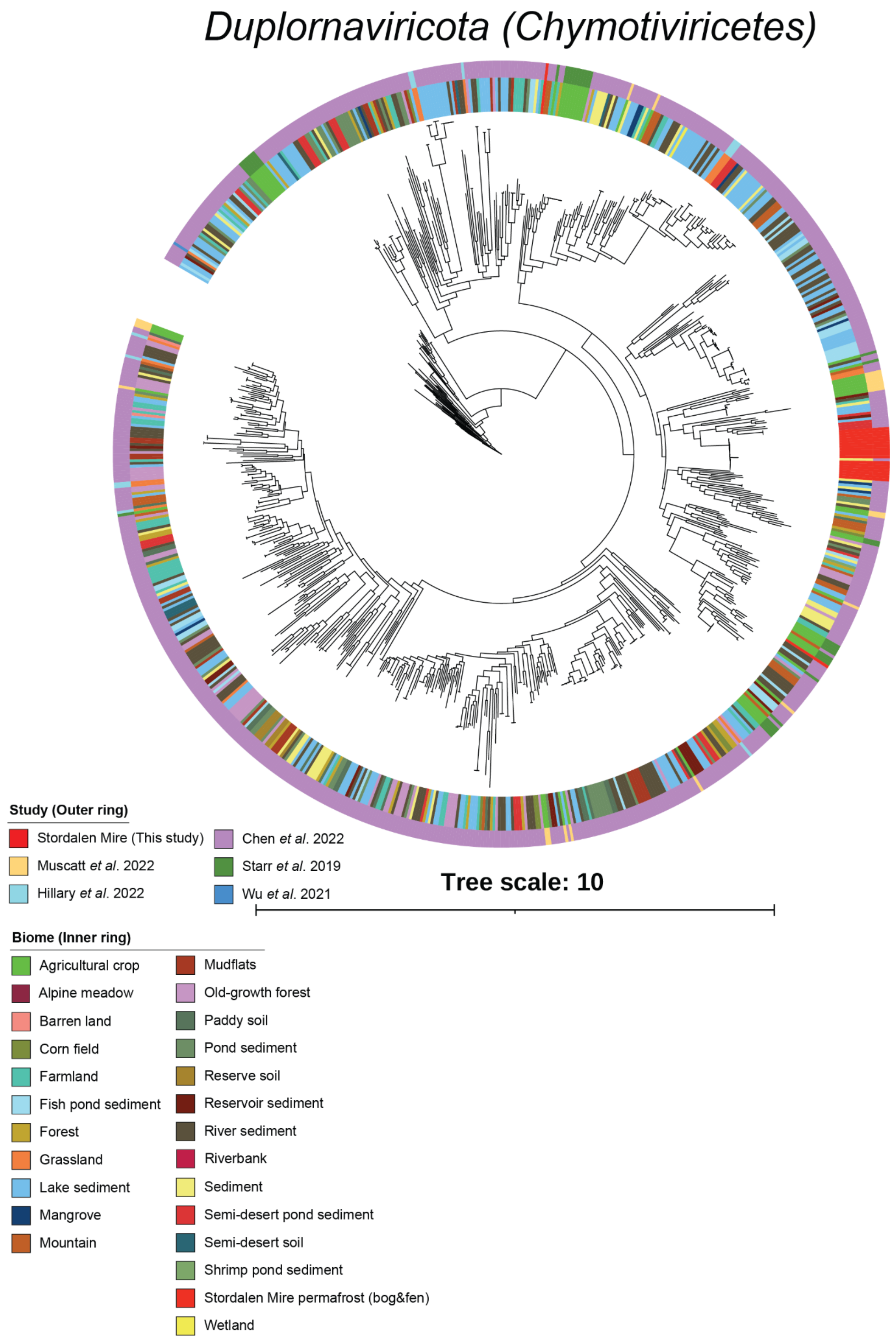


**Fig. S9. Thawing permafrost duplornaviricots (chrymotiviricetes). RdRp-based phylogenies across RNA virus studies.**

A maximum-likelihood phylogenetic tree was built from the RdRp-guided taxonomy analysis of near-complete RdRp domain sequences. The scale bar indicates one amino acid residue substitution per site. Sequences used to build the trees were preclustered at 40% identity, and clades supported by 100% bootstrap values were collapsed. The inner ring represents the biomes of these viruses whereas the outer ring represents soil RNA virus studies.

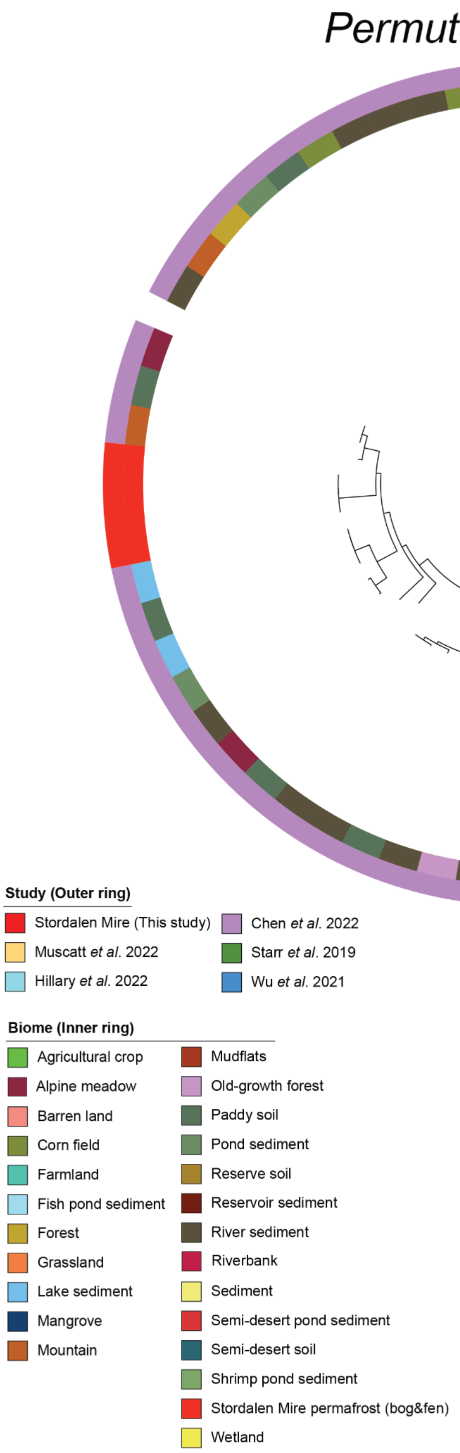


**Fig. S10. Thawing permafrost permutotetravirids. RdRp-based phylogenies across RNA virus studies.**

A maximum-likelihood phylogenetic tree was built from the RdRp-guided taxonomy analysis of near-complete RdRp domain sequences. The scale bar indicates one amino acid residue substitution per site. Sequences used to build the trees were preclustered at 40% identity, and clades supported by 100% bootstrap values were collapsed. The inner ring represents the biomes of these viruses whereas the outer ring represents soil RNA virus studies.

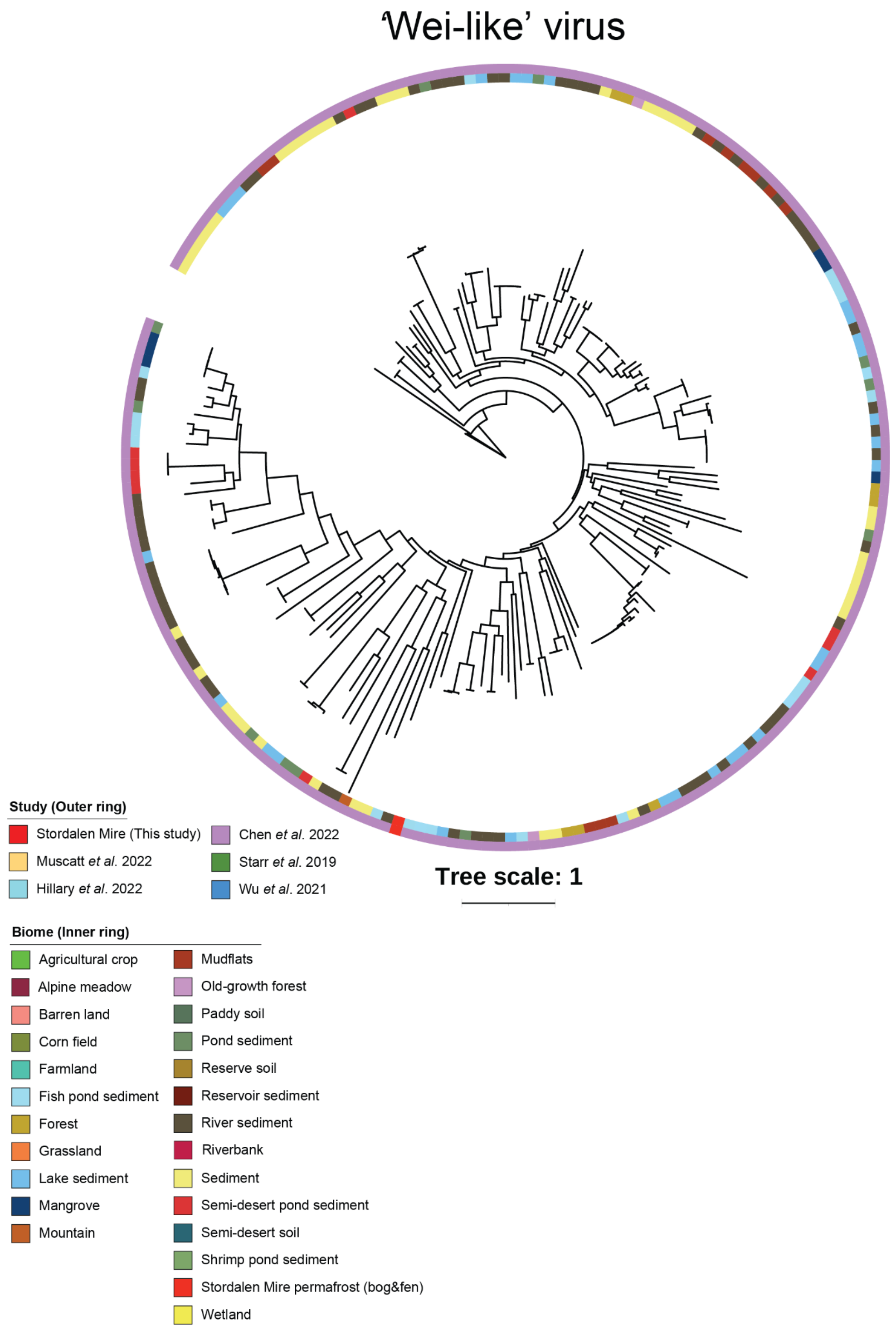


**Fig. S11. Thawing permafrost 'wei-like' viruses. RdRp-based phylogenies across RNA virus studies.**
A maximum-likelihood phylogenetic tree was built from the RdRp-guided taxonomy analysis of near-complete RdRp domain sequences. The scale bar indicates one amino acid residue substitution per site. Sequences used to build the trees were preclustered at 40% identity, and clades supported by 100% bootstrap values were collapsed. The inner ring represents the biomes of these viruses whereas the outer ring represents soil RNA virus studies.

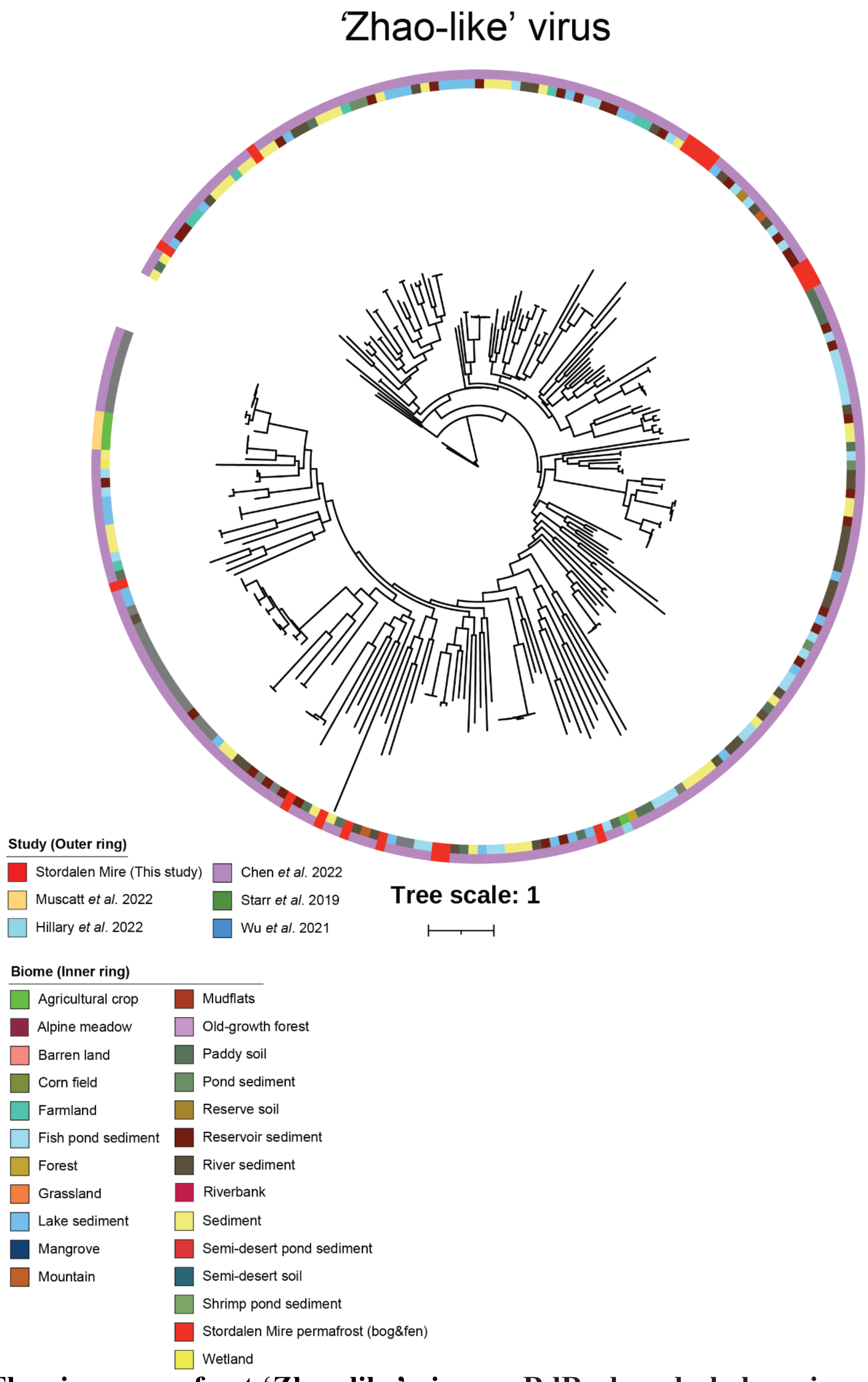


**Fig. S12. Thawing permafrost 'Zhao-like' viruses. RdRp-based phylogenies across RNA virus studies.**

A maximum-likelihood phylogenetic tree was built from the RdRp-guided taxonomy analysis of near-complete RdRp domain sequences. The scale bar indicates one amino acid residue substitution per site. Sequences used to build the trees were preclustered at 40% identity, and clades supported by 100% bootstrap values were collapsed. The inner ring represents the biomes of these viruses whereas the outer ring represents soil RNA virus studies.

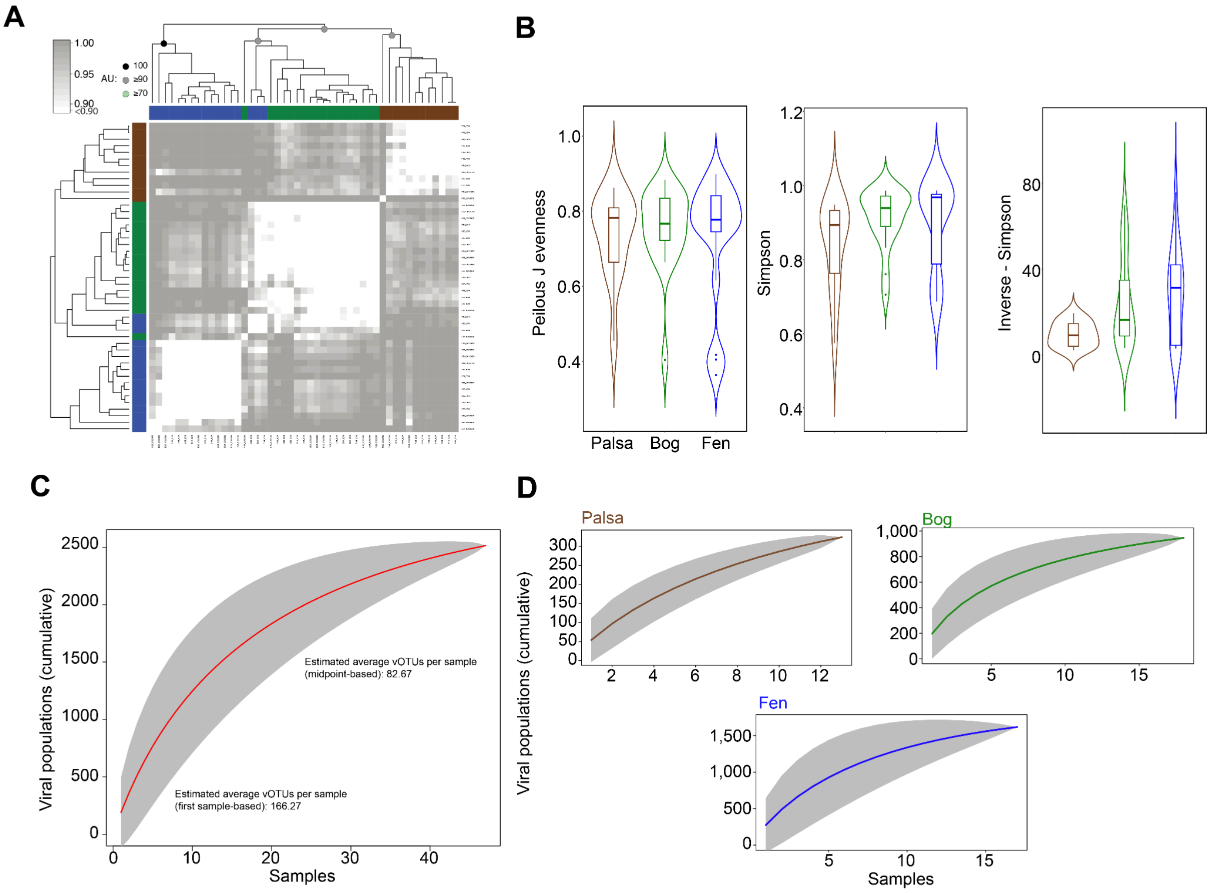


**Fig. S13. Stordalen Mire orthornaviraen ecology and diversity analysis.**
**(A)** Correlation-based hierarchical clustering of a Bray-Curtis dissimilarity matrix calculated from a randomly subsampled set from the vOTUs. The hierarchical clustering analysis structured orthornaviraens into three distinct meta-communities (brown; palsa, green; bog, and blue; fen) with an approximately unbiased (AU) bootstrap value ≥90. **(B)** Violin plots depict the diversity analysis of different metrics, including perilous J evenness, Simpson, and Inverse-Simpson. Diversity metrics were calculated using the "vegan" package in R. Statistical analysis was performed using Kruskal-Wallis analysis, with *post hoc* Dunn-test and *p*-adjusted: Bonferroni. Only the significant values are shown and donated as follows, *: $p$-value $\leq 0.05$; **: $p$-value $\leq 0.01$. **(C)** An accumulation curve of orthornaviraens in metatranscriptomes ($n = 49$, restricted to 2012 and 2016 data). Means are represented by red circles and 200 randomizations of sample order are shown in teal. Estimation of average vOTUs per sample from species accumulation curves. The first sample-based estimate (~166 vOTUs) represents an upper bound, while the midpoint-based estimate (~83 vOTUs) provides a more conservative average. and **(D)** Accumulative curve for palsa ($n = 13$), bog ($n = 18$), and fen ($n = 18$) (restricted to 2012 and 2016 data). Means are represented by brown, green, and blue circles (for palsa, bog, and fen, respectively), and the confidence interval (95%) of the sample order is shown in grey.

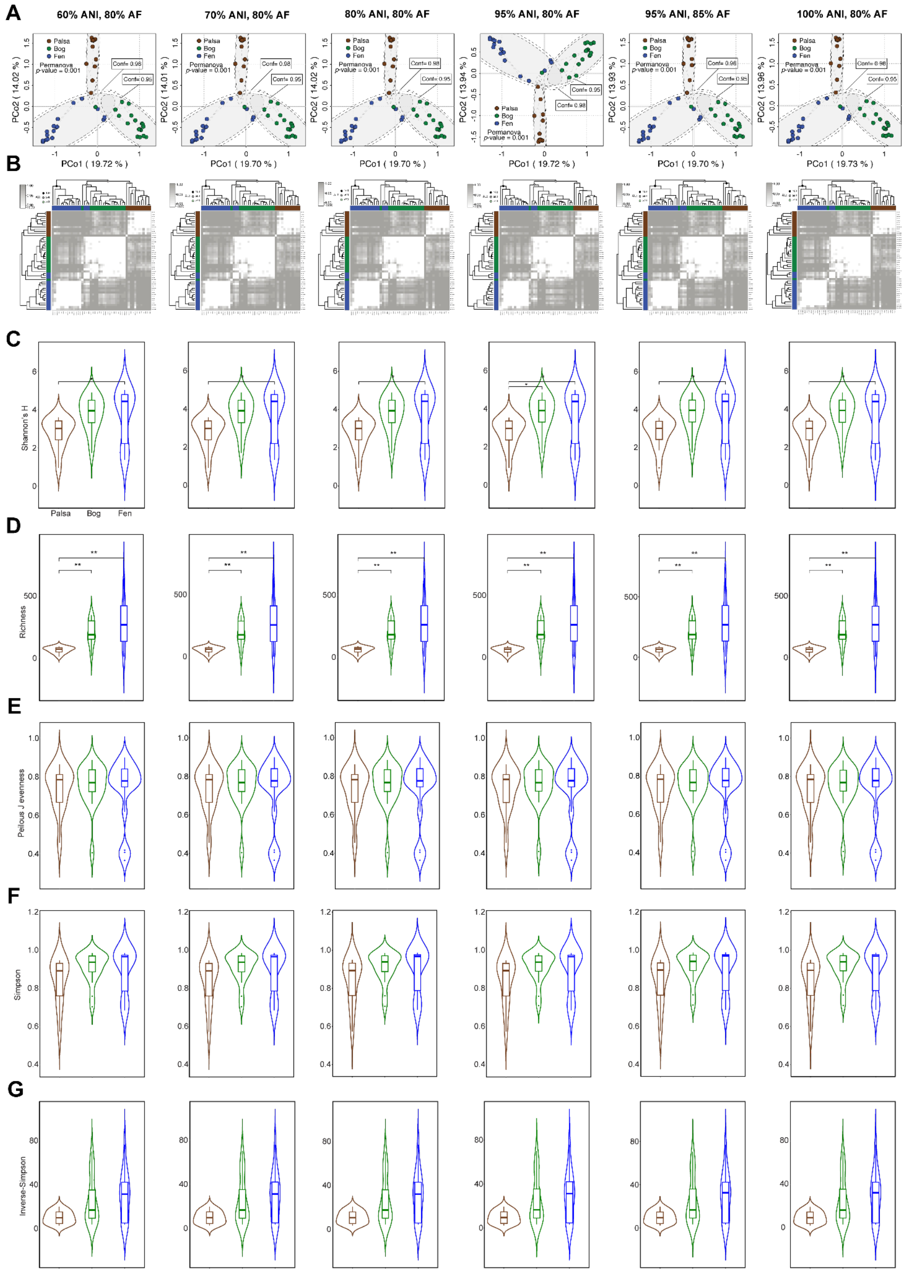


**Fig. S14. Sensitivity analyses for the robustness of ecological inferences under different vOTU definitions.**

(**A**) Principal component analysis (PCoA) of a Bray-Curtis dissimilarity matrix calculated from all vOTUs in this study. Dot colors correspond to Stordalen Mire habitats as in previous figures. (**B**) Correlation-based hierarchical clustering of a Bray-Curtis dissimilarity matrix calculated from a randomly subsampled set from vOTUs. Hierarchical clustering analysis sorted orthornaviraens into three distinct meta-communities (brown, palsa; green, bog; and blue, fen) with an approximately unbiased (AU) bootstrap value ≥90. (**C, D, E, F, G**) Violin plots (with boxplots) depict the diversity analysis of different metrics, including Shannon's *H*, richness, peilous J evenness, and Simpson and Inverse-Simpson. Statistical analysis was performed using Kruskal-Wallis analysis, with *post hoc* Dunn-test and *p*-adjusted: Bonferroni. Only significant values are shown and donated as follows, *: *p*-value ≤0.05; **: *p*-value ≤0.01.

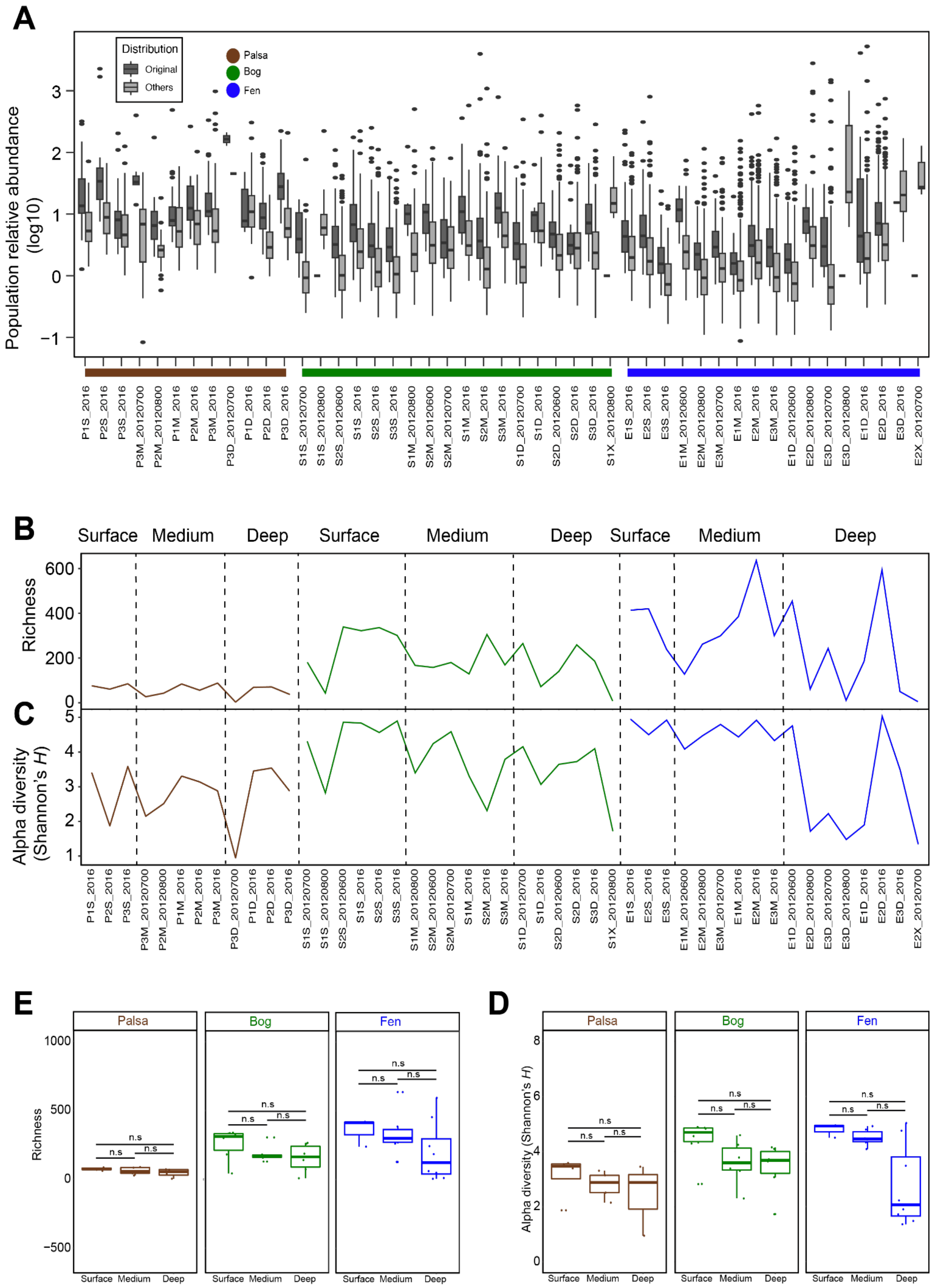


**Fig. S15. Ecological patterns of Stordalen Mire orthornaviraens across habitats.** (**A**) Relative abundance of vOTUs in original samples, from which vOTUs were assembled and compared in respect to abundance compared to other samples. (**B, C, D**, and **E**) Depth profiles of richness and alpha diversities (Shannon's *H*) of each sample, respectively. Samples were ordered based on depth, thaw gradient progress (palsa, bog, and fen) and sampling order. Statistical significance was evaluated with ANOVA (ns: not significant).

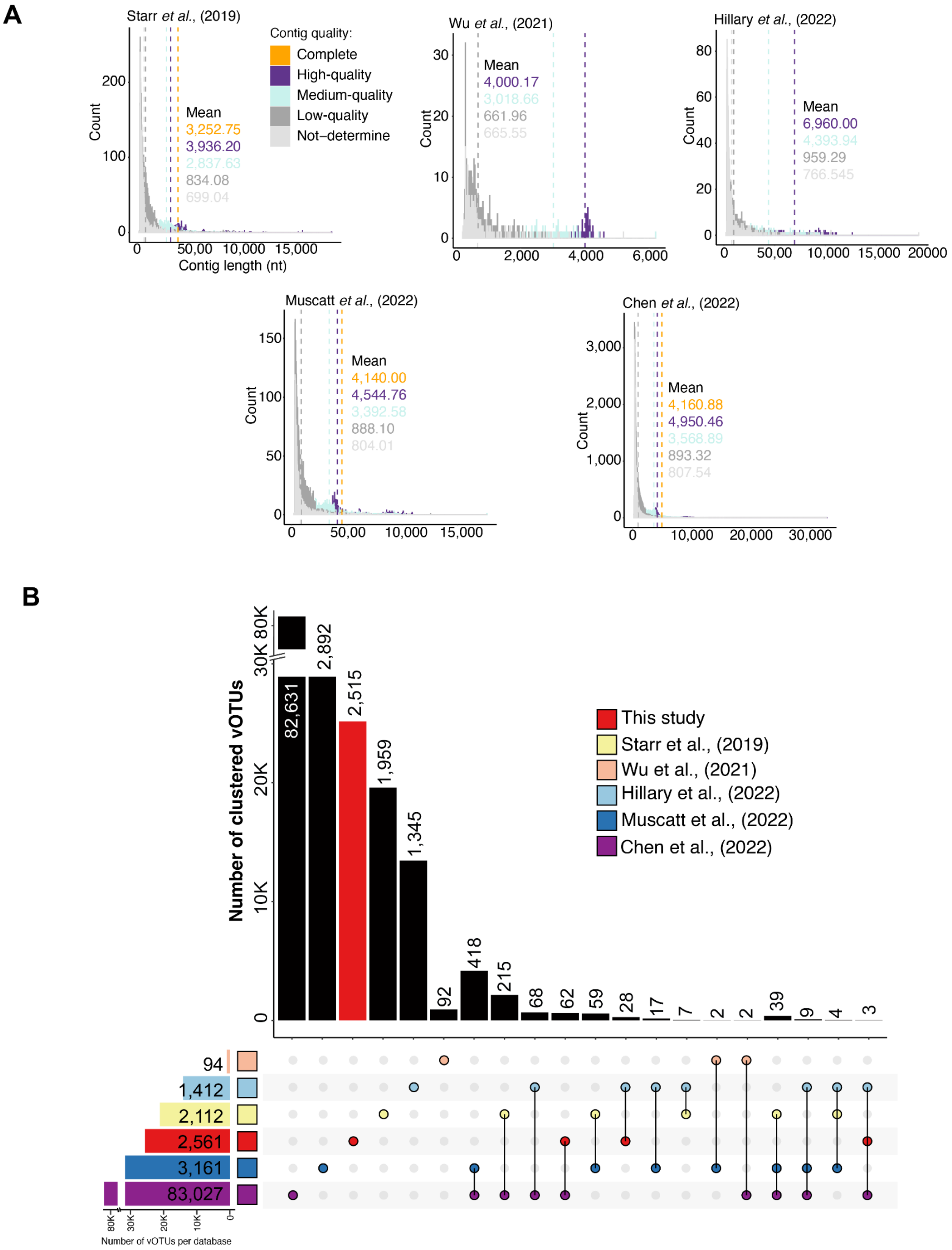


**Fig. S16. Comparison of orthornaviraen counts and RdRps across soil studies.** (**A**) The color dotted lines represent the length means of virus genomes based on contig quality. (**B**) UpSet plot depicts the number of shared and unique vOTUs (90% ID and 80% coverage).

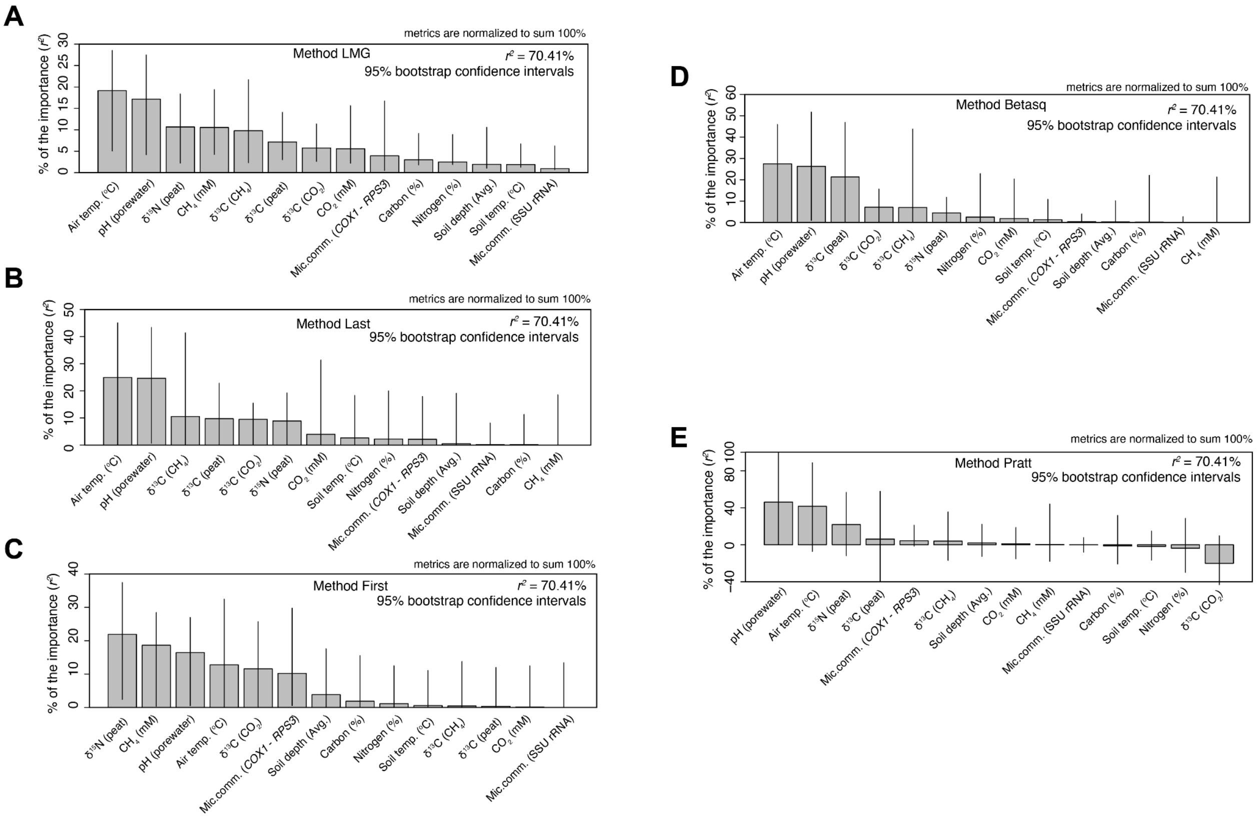


**Fig. S17. pH is a significant predictor of orthornaviraens in the Stordalen Mire thawing permafrost.**

Relative importance of environmental variables as predictors of estimated virus community (the first coordinate of a PCoA – Pco1). Analyses were performed using R package "relaimpo" v2.2-3 (*84*). This package enables univariate linear regression analyses using five different methods (**A**) LMG, (**B**) Last (**C**) First, (**D**) Betasq, and (**E**) Pratt, and 1,000 bootstraps for confidence estimates. $R^2$ denotes the proportion of response variance explained by the model. Bars show the mean proportion of response variance (i.e., relative contribution) of each variable based on 1,000 bootstrap replicates. Error bars indicate the 95% bootstrap confidence intervals of the response variance (in percentages). Only samples from 2011 and 2012 were used in the analysis, as no metadata were available for 2016.

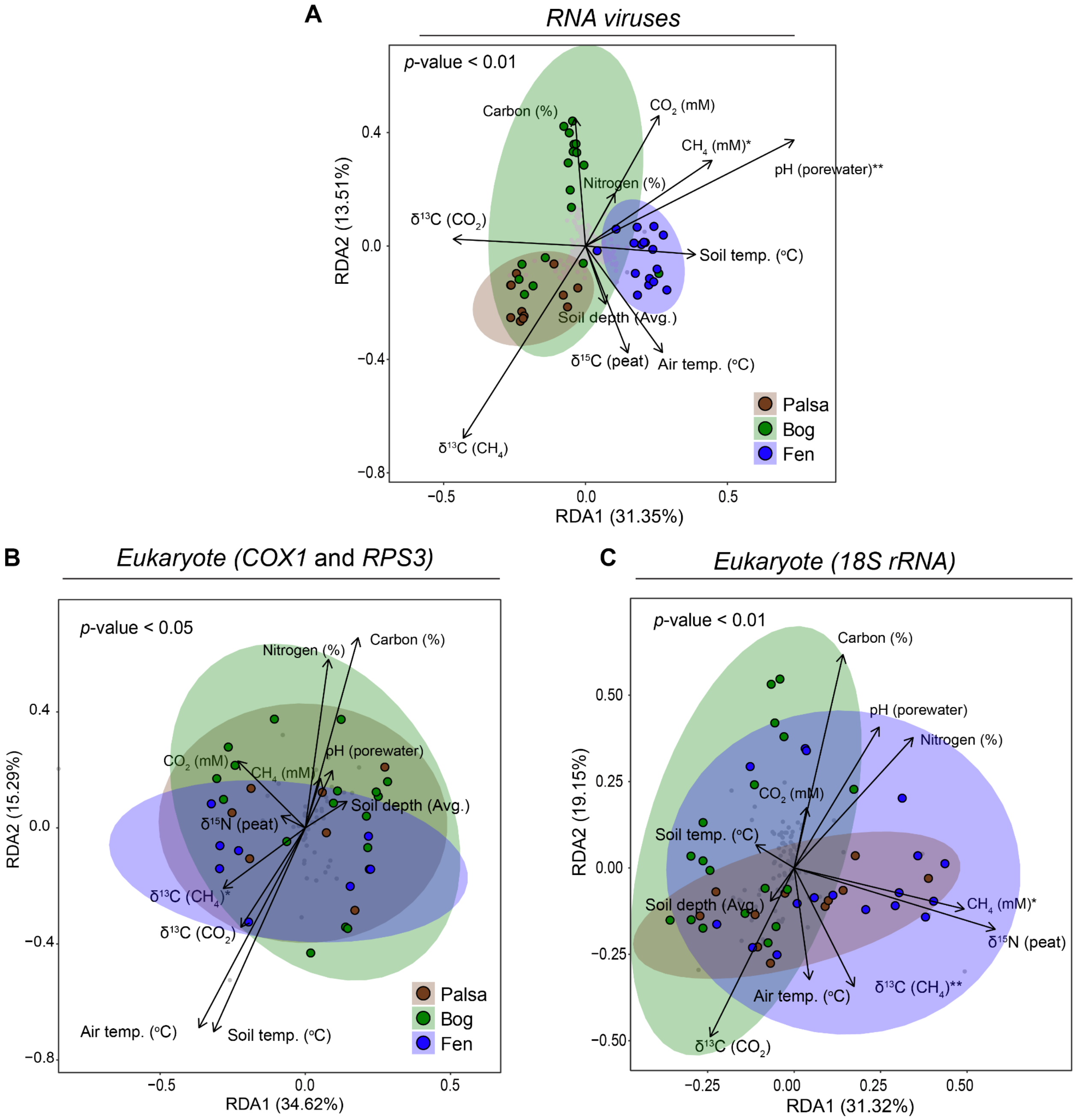


**Fig. S18. Redundancy Analysis (RDA) of RNA viruses and Stordalen Mire eukaryotes and prokaryotes.**

Redundancy analysis (RDA) of (**A**) RNA viruses, (**B**) Eukaryotes (based on *COX1* and *RPS3*) and (**C**) prokaryotes (based on 18S rRNA) of the relationship between the relative abundance of organisms across habitats and abiotic factors. Significance codes:; **: 0.01; *: 0.05.

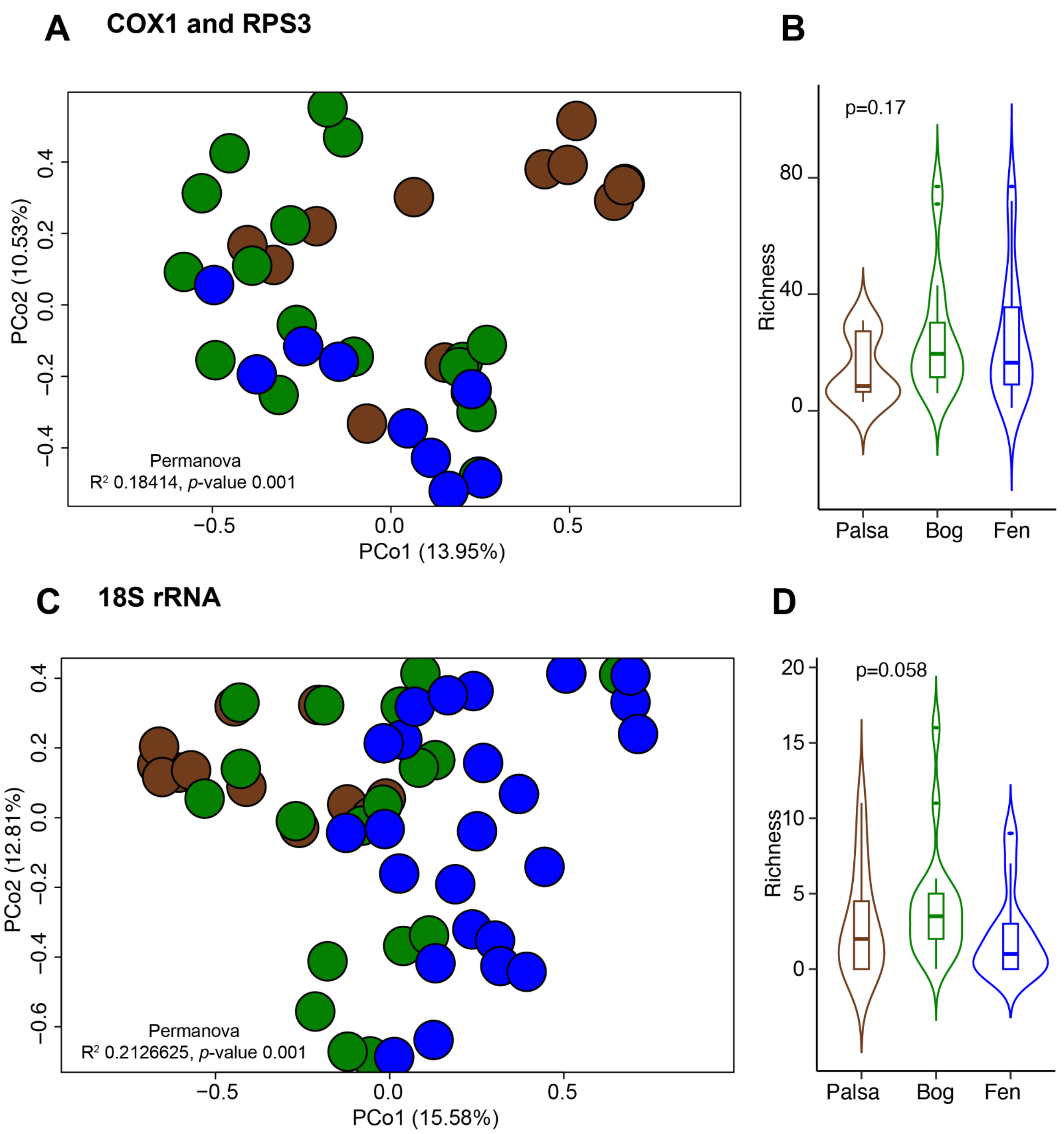


**Fig. S19. Organismal ecology.**
(**A** and **C**) Principal component analysis (PCoA) of a Bray-Curtis dissimilarity matrix calculated from all organism OTUs in this study deduced via marker gene and 18S rRNA sequencing, respectively. Violin plots (with boxplots) depict the Richness for (**B**) *COX1* and *RPS3* genes and (**D**) 18S rRNA, respectively. Statistical analysis was performed using Kruskal-Wallis analysis, with *post hoc* Dunn-test and *p*-adjusted/and ANOVA (for those with normal distribution data): Bonferroni.

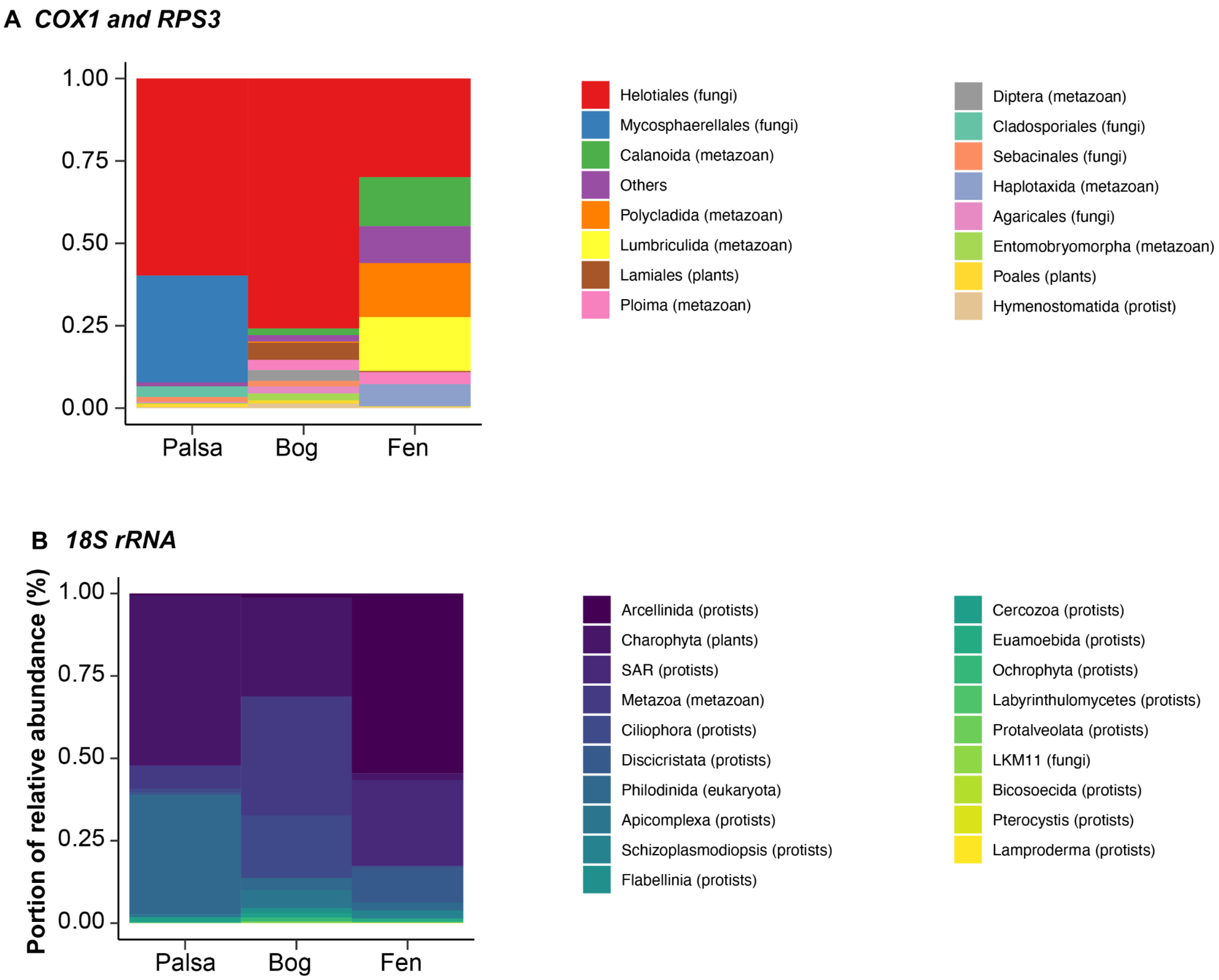


**Fig. S20. Microbial organism taxonomy across thawing gradients.**
The stacked-bar plots depict the proportion of relative abundance (%) based on metagenomics mapping. Taxonomy assignment at the order-level of organisms in Stordalen Mire identified by (**A**) *COX1* and *RPS3* genes, and (**B**) organisms identified by 18S rRNA gene.

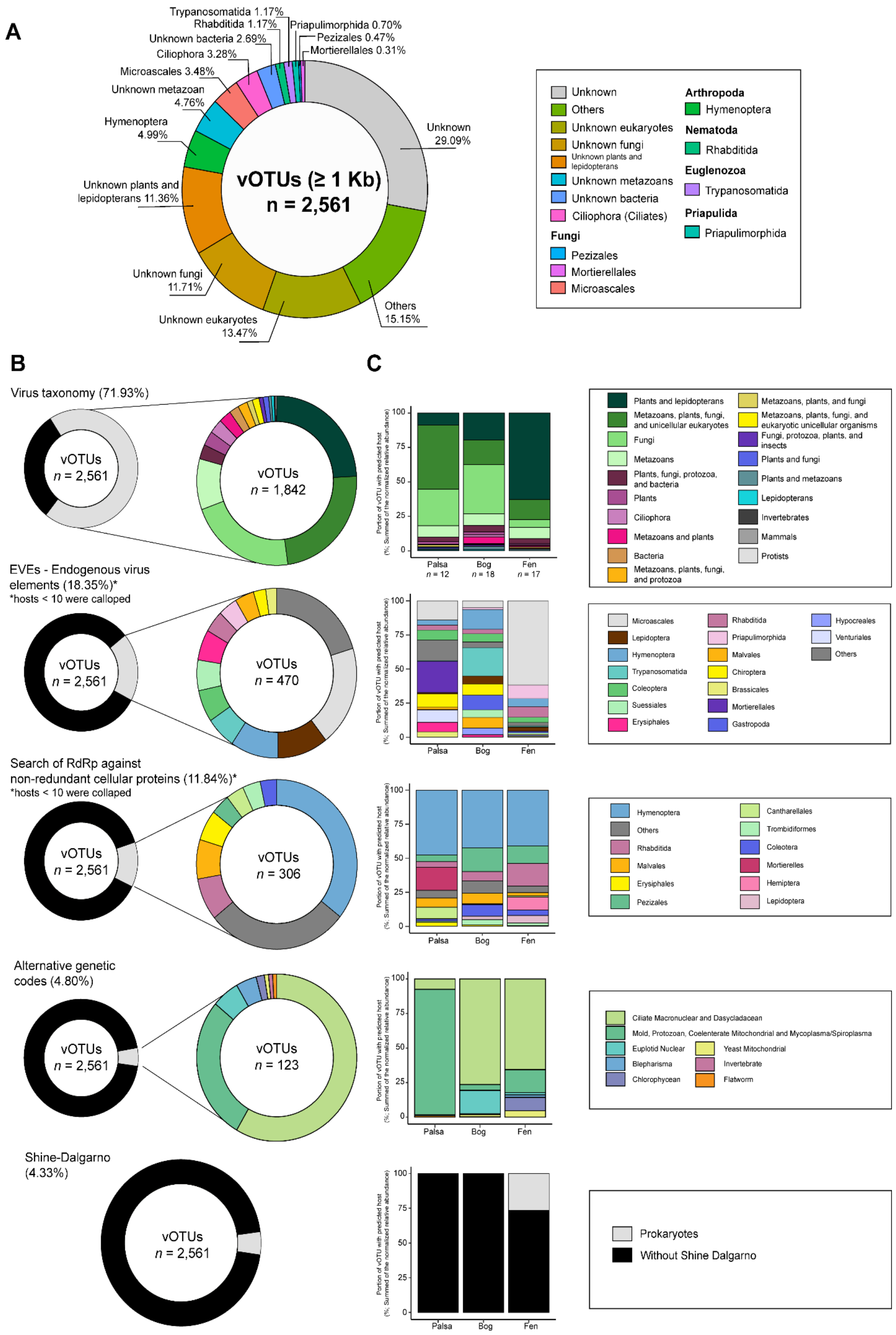


**Fig. S21. Inferred hosts of Stordalen Mire orthornaviraens.**
(**A**) Percent of inferred host based on the integrated approach. See analysis of inferred host using individual approach in panel B. (**B**) Host inferred analysis, including virus RdRp-based taxonomy, endogenous virus elements (EVEs), virus RdRps against non-redundant cellular proteins, alternative genetic codes and Shine-Dalgarno sequences. (**C**) Stacked bar plots depicting the percentage of vOTUs with the taxonomic affiliation of the inferred hosts per habitat. Only the top 10 taxonomy assignments are shown.

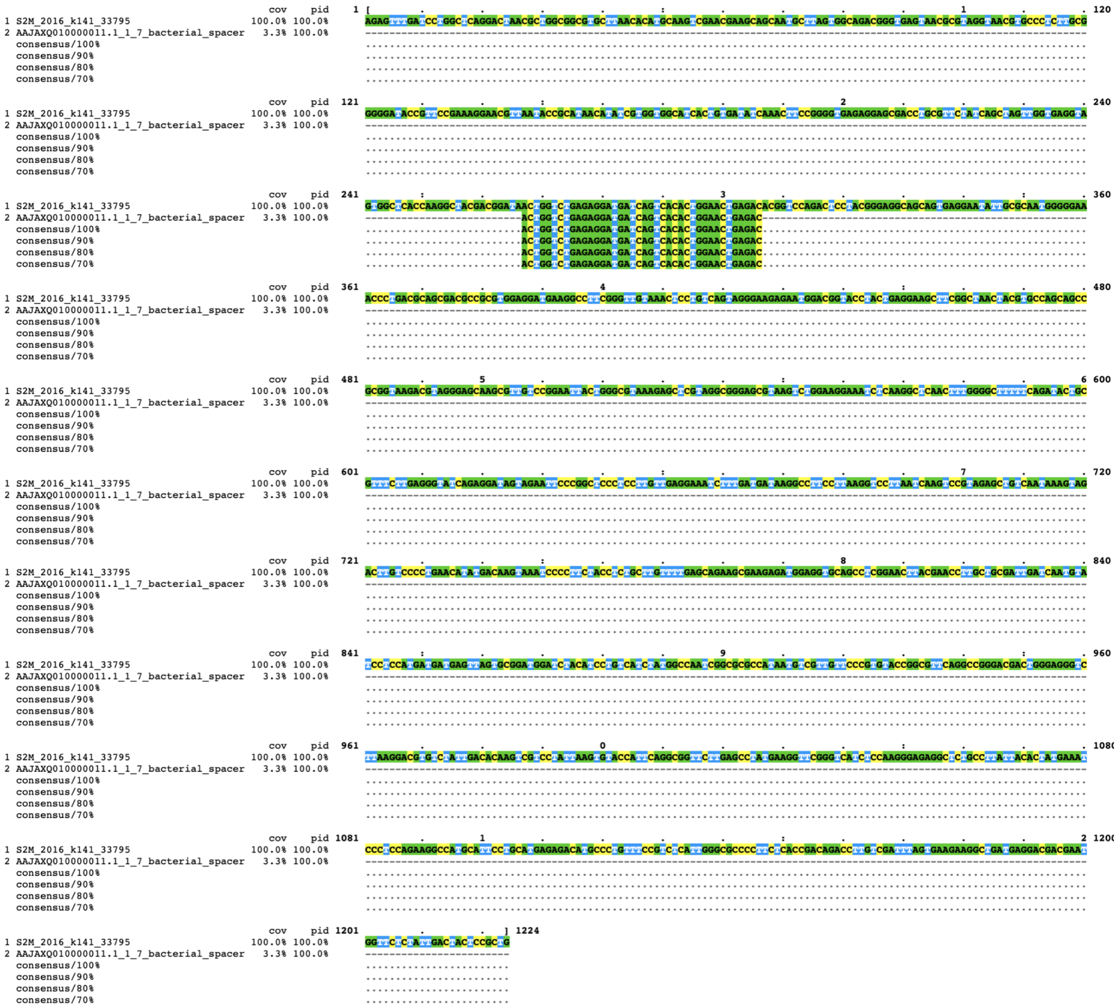

**Fig. S22. Search of CRISPR-Cas spacers across orthornaviran genomes.**
Global alignment of nucleotide sequences of an orthornaviraen contig (S2M_2016_k141_33795) and a CRISPR spacer of *Campylobacter lari* (AAJAXQ010000011.1) visualized using MView.

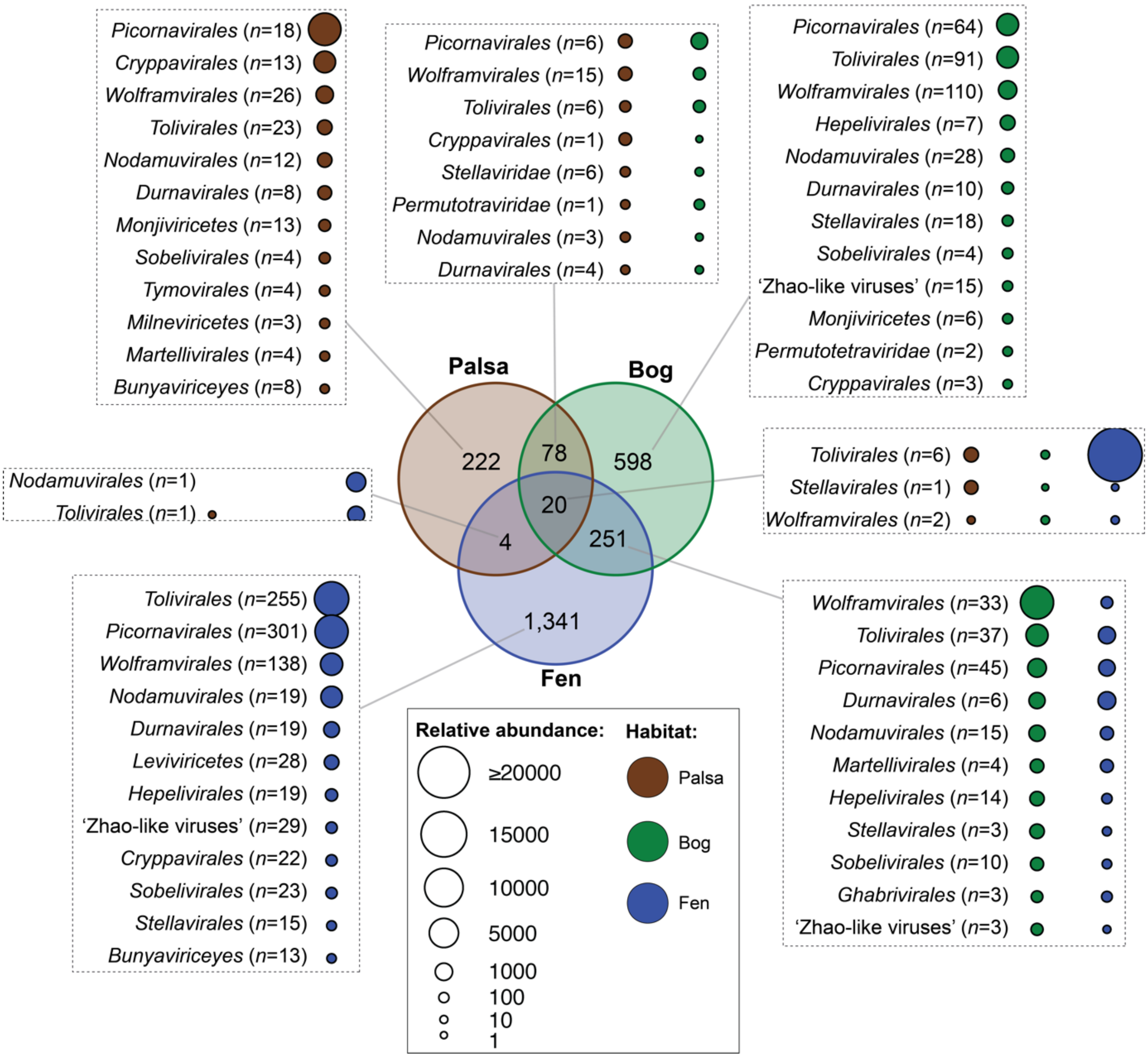


**Fig. S23. Venn diagram of Stordalen Mire orthornaviraens across habitats.**
A modified RdRp-scan (*80*) approach was used to assign order-ranked virus taxonomy to RdRp footprint sequences longer than 200 amino acid residues encoded by vOTUs. Dot sizes represent the summed relative abundances of vOTUs detected across habitats.

### Supplementary Tables

**S1 Table. Accession numbers Stordalen Mire metatranscriptomes used in this study.**
**S2 Table. Metadata: sampling IDs, locations, depth, biochemical measurements.**
**S3 Table. List of final RNA virus footprints and contigs.**
**S4 Table. Normalized relative abundances of vOTU.**
**S5 Table. MCL Network-based taxonomy assignment**
**S6 Table. Metagenomics assembed genomes were used in this analysis. Acquired from Woodcroft et al. 2018. Nature 560:49–54 ; 10.1038/s41586-018-0338-1**
**S7 Table. Integrated host interference analysis**
**S8 Table. Final auxiliary metabolic genes in RNA viruses**
